# Widespread divergent transcription from prokaryotic promoters

**DOI:** 10.1101/2020.01.31.928960

**Authors:** Emily Warman, David Forrest, Joseph T. Wade, David C. Grainger

**Affiliations:** Institute for Microbiology and Infection, School of Biosciences, University of Birmingham, Edgbaston, Birmingham, B15 2TT, UK; Wadsworth Centre, New York State Department of Health, Albany, NY, 12208, USA; Department of Biomedical Sciences, University at Albany, Albany, NY, 12201, USA

## Abstract

Promoters are DNA sequences that stimulate the initiation of transcription. In all prokaryotes, promoters are believed to drive transcription in a single direction. Here we show that prokaryotic promoters are frequently bidirectional and drive divergent transcription. Mechanistically, this occurs because key promoter elements have inherent symmetry and often coincide on opposite DNA strands. Reciprocal stimulation between divergent transcription start sites also contributes. Horizontally acquired DNA is enriched for bidirectional promoters suggesting that they represent an early step in prokaryotic promoter evolution.

---

Transcription initiation requires DNA sequences called promoters that interact with RNA polymerase (RNAP)^1^. Promoters consist of ordered core elements with distinct roles^2,3^. For example, most bacterial promoters contain a −10 element that interacts with the housekeeping RNAP σ^70^ subunit. This facilitates DNA unwinding^4,5^. In eukaryotes and archaea, the TBP binding TATA box has a similar role^6^. It has long been assumed that promoters are directional, driving transcription in a single orientation determined by promoter element arrangement^2,7^. This view has recently been challenged in eukaryotes^8–11^. Nonetheless, the consensus view is that prokaryotic promoters are unidirectional^12^.

Previous studies have mapped transcription start sites (TSSs) in *Escherichia coli* by detecting triphosphorylated RNA 5’ ends^13^. These TSSs can be assigned to σ^70^ binding events identified using ChIP-seq^13^. We noticed that not all σ^70^ binding sites were associated with detectable RNA 5’ ends. This was particularly evident for horizontally acquired genes silenced by histone-like nucleoid structuring (H-NS) protein (Figure S1). We reasoned that RNAP might initiate transcription but produce unstable RNAs. To test this, we fused 33 such σ^70^ targets to *lacZ*. Any transcripts produced should be stabilised, and detectable, due to translation. Transcription orientation cannot be directly inferred from σ^70^ ChIP-seq data. Hence, DNA sequences were cloned in both directions (Figure 1a). Surprisingly, over half of the fragments were transcriptionally active in both orientations (Figure 1b). We designated the direction of highest *lacZ* expression as “forward”. On average, “reverse” transcription neared half the “forward” activity (Figure 1c). For a subset of divergent transcript pairs, we mapped RNA 5’ ends (Figure 1d). Most reverse TSSs were upstream of the forward TSS and resulted from shared overlapping promoter elements (Figures 1e and S2). Mutations in shared promoter elements (Figure S2) reduced expression in both orientations (Figure 1f).

**Figure 1:**
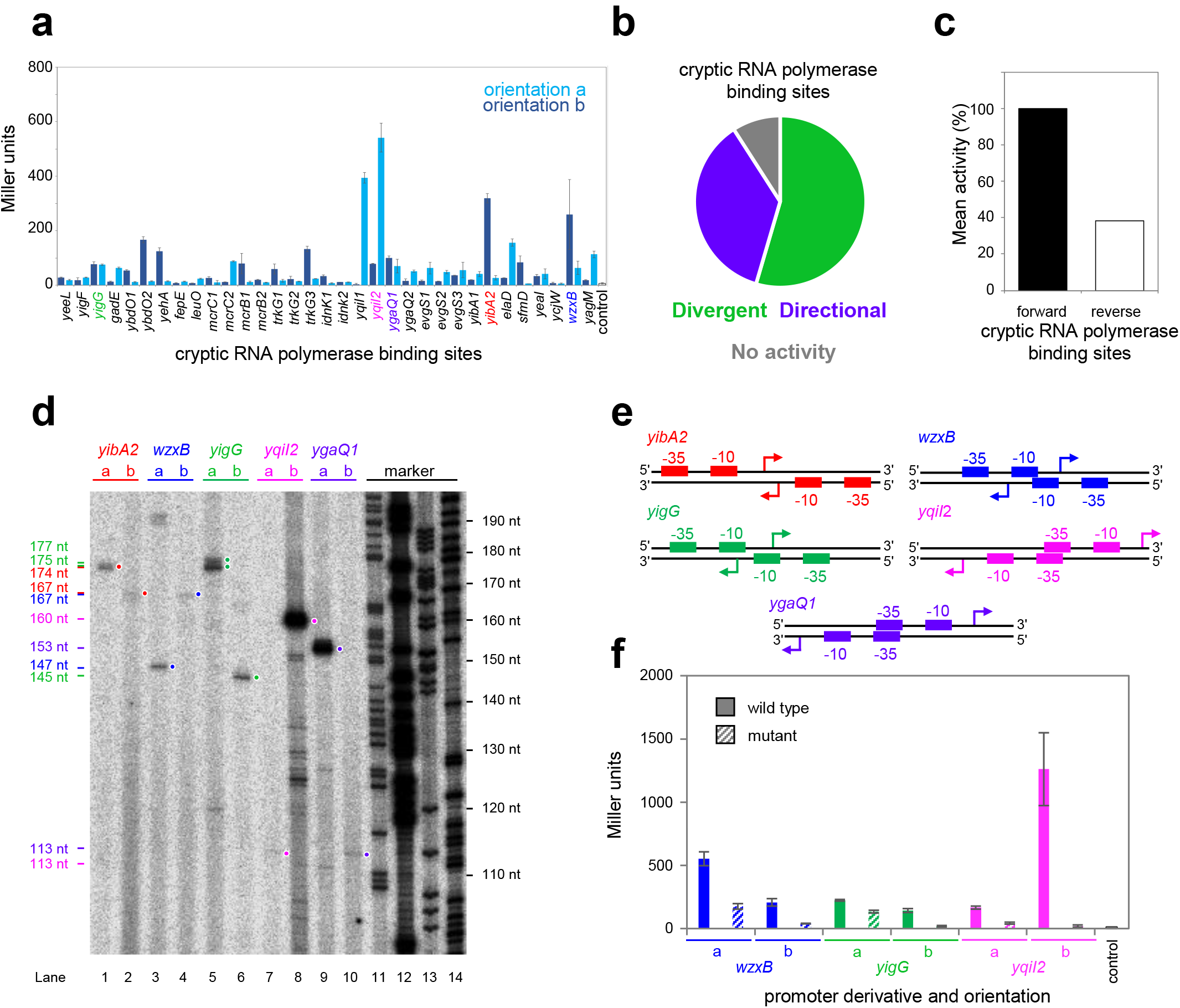
Divergent transcription within horizontally acquired genes. a) β-galactosidase activity derived from cryptic RNAP binding sites. b) Direction of transcription from cloned DNA fragments. c) Average forward or reverse β-galactosidase activity of all DNA fragments. d) Start sites mapped by primer extension for selected DNA fragments (orientations labelled a or b). Primer extension products in lanes 1 to 10, sizes in nucleotides (nt). Lanes 11-14 are Maxam-Gilbert sequencing reactions for calibration. e) Schematic representation of core promoter elements associated with divergent transcription. f) Effect of mutating shared core promoter elements.

To understand global patterns of divergent transcription we analysed TSSs independently mapped by RNA 5’ polyphosphatase sequencing (PPP-seq), dRNA-seq or cappable-seq^13–15^. In all cases, oppositely orientated TSSs tended to co-locate (Figure 2a). To increase sensitivity, we merged the datasets (Figure 2a, combined). This identified 5,292 divergent TSSs, defined as being separated by between 25 and 7 bp; 19 % of all detected TSSs in *E. coli*. We refer to the associated promoters as bidirectional. The most common distance between divergent TSSs was 18 bp; transcription initiates either side of overlapping −10 elements (Figure 2a, top expansion). We reasoned that promoter element symmetry must play a major role. To test this, we made a position weight matrix (PWM) describing all *E. coli* promoter sequences. If the PWM matched adjacent regions of DNA on opposite strands the symmetry score increased. Maximum symmetry correlated with divergent transcription (R^2^ = 0.85; Figure 2a bottom expansion). Consistent with this, a DNA sequence logo generated by aligning divergent TSSs, separated by 18 bp, was symmetrical (Figure 2b). Contrastingly, TSSs with no divergent transcript generated an asymmetrical motif (Figure 2c). Note that the first, second and sixth positions of promoter −10 elements (consensus 5’-TATAAT-3’) are key for transcription initiation^5^ (Figure 2c). At divergent TSS offset by 18 bp, nucleotides two and six of −10 elements on opposite DNA strands base pair. Hence, these positions are most strongly conserved (Figure 2b). Example −10 elements arranged in this way are shown in Figure 2d. Divergent transcription also increased at TSSs separated by 29, 23, 12, 10 or 7 bp (Figure S3a). These configurations also correspond to symmetrical base pairing between key −10 element nucleotides, and TSSs, on opposite DNA strands (Figure S3b). The distribution of all bidirectional promoters with respect to genes is shown in Figure 2e.

**Figure 2:**
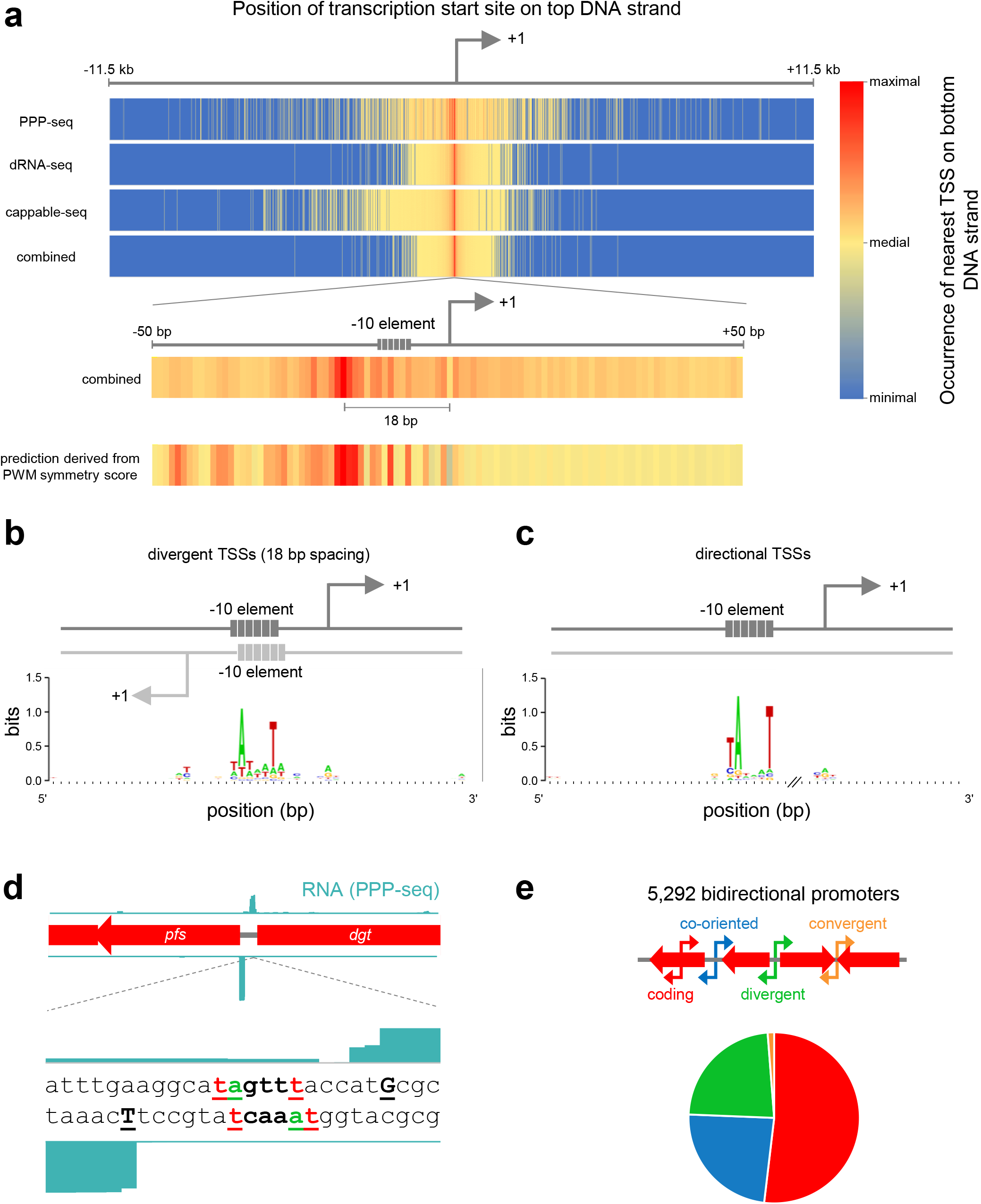
Widespread divergent transcription from bidirectional promoters in *Escherichia coli*. a)Heatmaps made using global transcription start site (TSS) data^13–15^ or position weight matrix analysis. TSSs on the top chromosome strand are aligned at the centre of the heatmap (bent arrow, labelled +1). Heatmap colour indicates abundance of bottom strand TSSs at that position. The expansion shows the occurrence of bottom strand TSSs in a 50 bp window either side of all top strand promoters. b) Predominant DNA sequence motif associated with bidirectional or c) directional promoters. The x-axis break indicates the variable distance between −10 element and TSS at directional promoters. d) a bidirectional promoter between the *E. coli pfs* and *dgt* genes. Promoter −10 elements are bold. TSSs are in uppercase. e) Relative position of all bidirectional *E. coli* promoters with respect to genes.

In *E. coli* transcription preferentially initiates at an adenine (Figure 2c). For divergent TSSs 18 bp apart, the +1 nucleotide corresponds to position −18on the opposite DNA strand. Hence, −18is often a thymine (Figure 2b). A thymine at position −18can increase transcription by improving interaction between σ^70^ residue R451 and the DNA backbone^16^ (Figure 3a). We speculated that the +1/-18 overlap could explain why this configuration is so frequently detected. To test this, we cloned a bidirectional promoter, with 18 bp between TSSs, in both orientations upstream of the *λoop* transcriptional terminator (Figure 3bi). We also made derivatives where the A•T at each +1/-18 position was replaced with C•G (Figure 3bii-iii). We measured RNA synthesis terminated by *λoop* using *in vitro* transcription (Figure 3c). As expected, altering the TSS reduced production of the associated RNA (compare lane 1 with 5 and 3 with 11); the same mutations also reduced transcription in the opposite direction (compare lane 1 with 9 and 3 with 7). Though σ^70^ RA451 was defective at the bidirectional promoters (even lane numbers to 12) it was unimpaired at a control promoter not requiring this contact (lanes 13-14).

**Figure 3:**
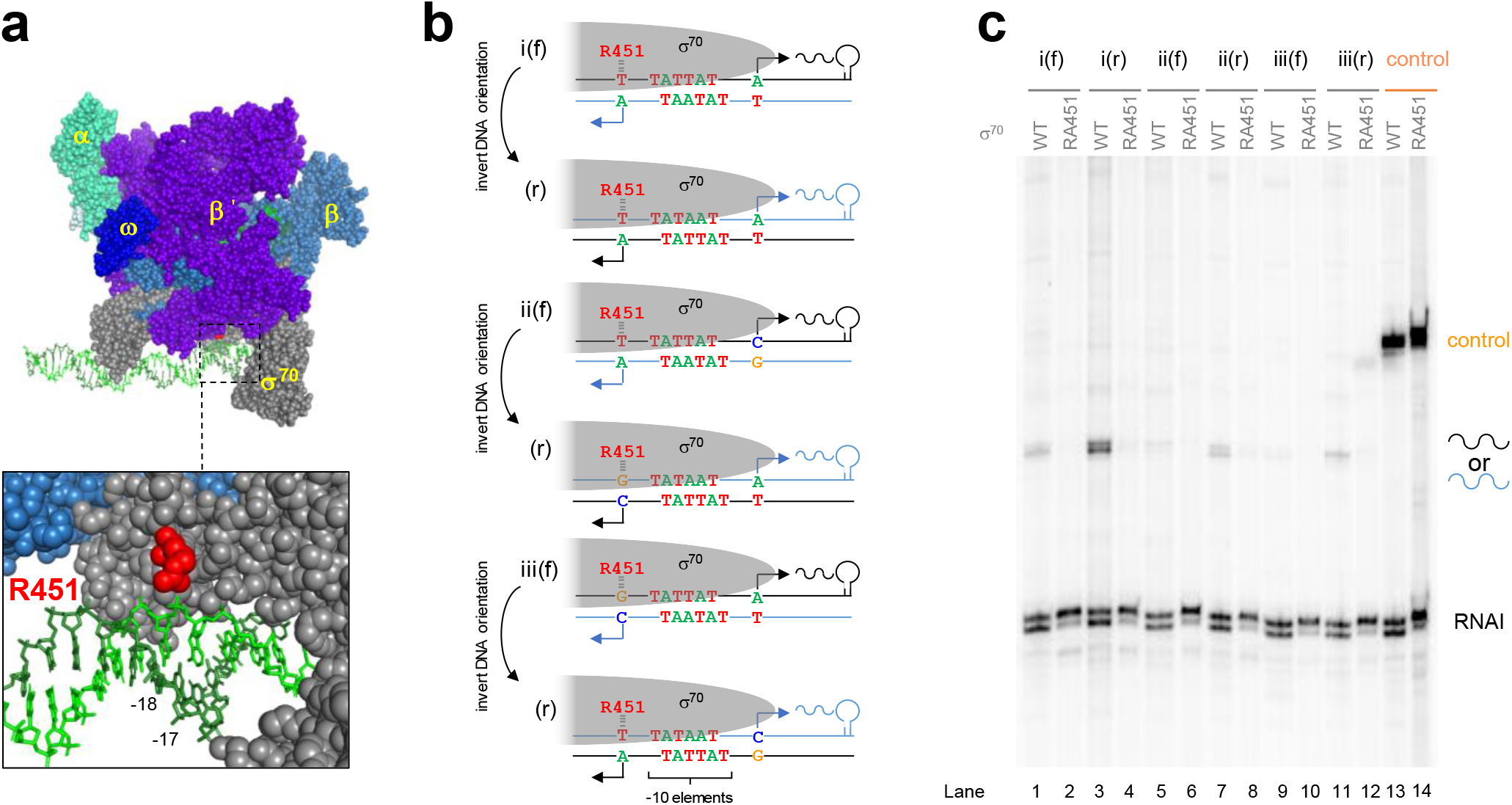
Reciprocal stimulation between divergent transcription start sites. a) Structure of RNAP bound to DNA (PDB: 6CA0)^28^. Relevant features labelled. b) DNA templates used for *in vitro* transcription. For simplicity, only the sequences of promoter −10 elements (labelled) and TSSs (bent arrows) are shown. Opposing DNA strands represented by black or blue lines. Interaction between σ^70^ R451 and the DNA backbone indicated by dashes. c) Products of *in vitro* transcription (using templates in panel b) using either σ^70^ or the R451A derivative. The RNAI transcript is derived from the replication origin of the plasmid DNA template.

To determine the prevalence of bidirectional promoters in bacteria we analysed TSS maps for proteobacteria^13–15,17–20^, actinobacteria^21,22^, and a firmicute^23^. We also mapped TSSs in an additional firmicute, *Bacillus subtilis*, using cappable-seq (summarised in Figure S4 and Table S1). Co-localised divergent TSSs were abundant in all bacteria analysed (Figure 4a). Proteobacteria and actinobacteria were most similar; divergent TSSs were most frequently separated by 18 or 19 bp, and shared a near-identical symmetrical −10 element with *E. coli* (Figure S5). Firmicutes used the same range of −10 element configurations illustrated in Figure S3 for *E. coli*, albeit with little preference for a single arrangement (Figure S5). For all bacteria, spacing intervals associated with divergent transcription scored highest for symmetry (Figure S5)

**Figure 4:**
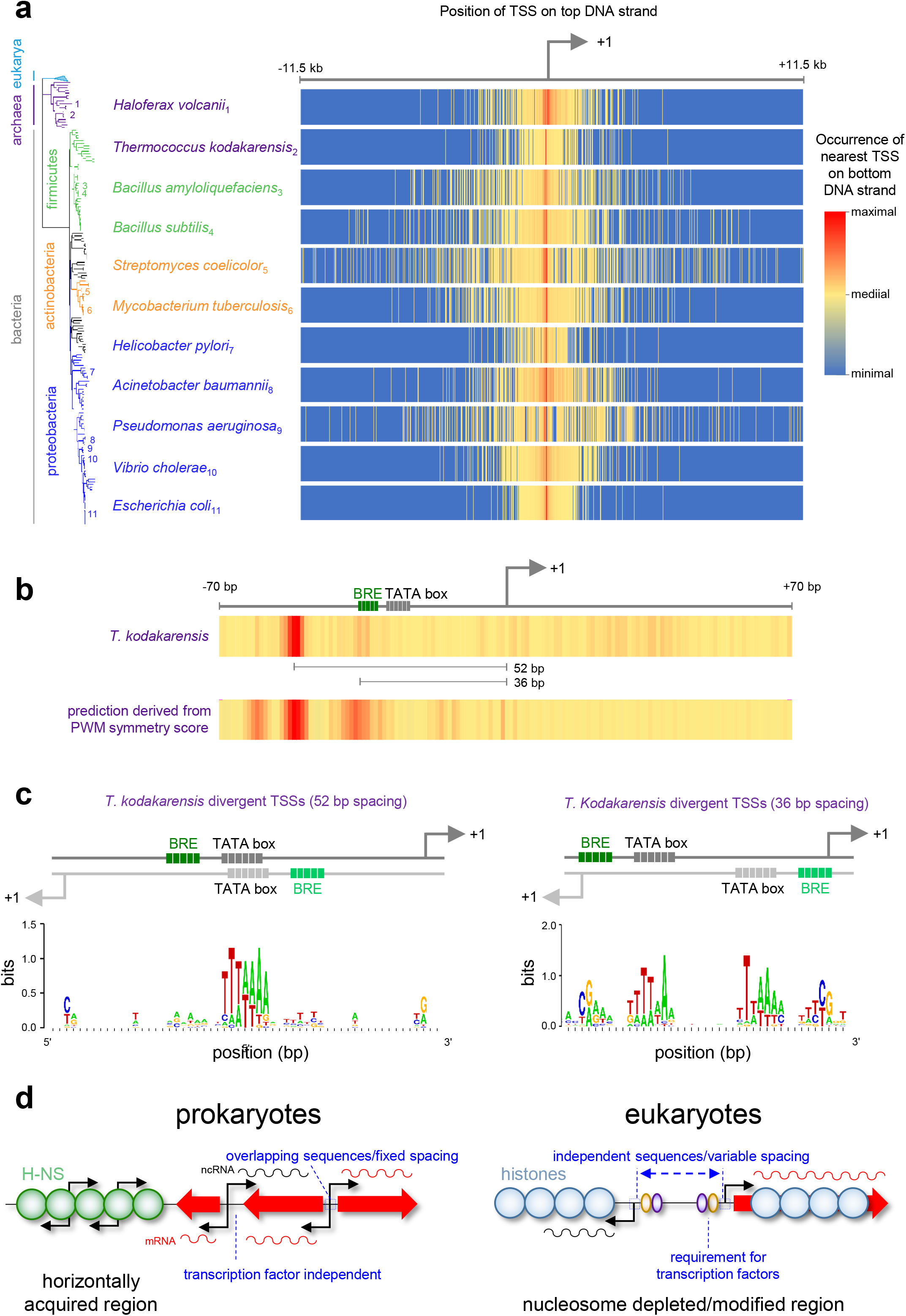
Bidirectional promoters are widespread in prokaryotes. a,b) Heatmaps indicate abundance and position of TSSs on the bottom DNA strand, relative to the nearest top strand promoter (bent arrow). Species and phylogenetic relationships are indicated to left of heatmaps. c) DNA sequence motifs derived from divergent TSSs in *T. kodakarensis*. d) Bidirectional promoters have a different basis in prokaryotes and eukaryotes.

Archaeal transcription is closely related to that of eukaryotes; promoters have a TATA box and B recognition element (BRE), located a narrow range of distances from the TSS^24^. We analysed TSS maps for the archaea *Thermococcus kodakarensis* and *Haloferax volcanii*^25,26^. We observed strong signatures of promoter bidirectionality (Figure 4a). In *T. kodakarensis*, divergent TSSs were predominantly separated by 52 bp and located either side of a shared TATA box element (5’-TTATAAA-3’) (Figure 4b,c and S6a). Less frequently, TSSs separated by 36 bp were used (Figures 4b and S6a). Here, the B recognition element (BRE; 5’-CGAAA-3’) is positioned so the initial C•G bp can also act as the TSS on the opposite DNA strand (Figure 4c). Similar observations were made for *H. volcanii* despite the unusual TATA box consensus (5’-TTWT-3’) of haloarchaea (Figure S6b,c). For both species, an independent promoter PWM search identified near identical spacing rules (Figure S6).

Our data demonstrate that divergent transcription from promoters is a process conserved in all life forms. The phenomenon is similarly frequent in diverse prokaryotes (Figure S7) and superficially resembles the situation in eukaryotes. However, the mechanistic basis is fundamentally different (Figure 4d). In eukaryotes, bidirectionality is generated by an activator protein creating two adjacent regions of nucleosome depletion^27^. Thus, divergent TSSs use separate core promoter elements that can be separated by thousands of bp, with no distance optimal. By contrast, divergent transcription in prokaryotes depends on symmetry of key promoter elements; TSSs on opposite strands are closely spaced at preferred intervals. Consequently, for prokaryotes, divergent transcription can be predicted using DNA sequence and recapitulated *in vitro* with purified components. In eukaryotes, recently acquired DNA is enriched for bidirectional promoters^27^. This has been attributed to pervasive transcription factor binding^27^. We initially identified divergent transcription in horizontally acquired *E. coli* DNA (Figure 1). Furthermore, detection of bidirectional promoters increased in cells lacking H-NS (Figure S8a). This results from elevated promoter frequency and symmetry in foreign genes (Figure S8b). Hence, divergent transcription and promoter evolution are linked in prokaryotes. Strikingly, the proportion of bidirectional promoters used for mRNA production is higher than the equivalent fraction of canonical promoters (Figures 2e, S4d). We conclude that divergent transcription plays a key role in prokaryotic cells.

