## Supplementary material for "Widespread divergent transcription from prokaryotic promoters": Materials and Methods

### MATERIAL AND METHODS

*Strains, plasmids and oligonucleotides*

All strains plasmids and oligonucleotides used are listed in Table S2. Standard procedures for strain and DNA manipulation were used throughout. All bacterial cultures were grown in LB media.

*Transcription start site mapping*

Transcript start sites were mapped for individual promoters using primer extension as described by Haycocks and Grainger[1]. The RNA was purified from indicated *E. coli* strains carrying different DNA fragments cloned in pRW50. The 5' end-labelled primer D49724, which anneals downstream of the *Hin*dIII site in pRW50, was used in all experiments. Primer extension products were analysed on denaturing 6% polyacrylamide gels, calibrated with size standards, and visualized using a Fuji phosphor screen and Bio-Rad Molecular Imager FX. To map TSSs globally we used cappable-seq. Duplicate cultures of *B. subtilis* strain 168 ca were grown until mid-exponential phase in LB media with shaking at 37ºC. Cells were harvested and flash frozen in liquid nitrogen. Total RNA was isolated as described previously with the exception that RNA concentration and quality was determined on an Agilent 2200 Tapestation following the manufacturer's instructions[2]. Library preparation and sequencing was done by Vertis Technologie AG according to the protocol described by Ettwiller *et al.*[3]. Briefly, 5' triphosphorylated RNA was capped with 3'- desthiobiotin-TEG-guanosine 5' triphosphate (DTBGTP) using Vaccinia capping enzyme (New England Biolabs). Biotinylated RNA was captured and eluted from streptavidin beads to obtain 5' fragments of primary transcripts. These transcripts were poly(A) tailed with poly(A) polymerase before conversion of the 5' CAP moiety to a 5' monophosphate using CAP-clip Acid pyrophosphatase (Cellscript). An RNA adapter was ligated to the 5' monophosphate and cDNA synthesis was done with an oligo(dT)-adapter primer and M-MLV reverse transcriptase. cDNAs were amplified by PCR to a final concentration of 10-20 ng $\mu l^{-1}$. Full length cDNAs were fragmented and immobilised with streptavidin magnetic beads for blunting and ligation of the 3' Illumina sequencing adapter. The immobilised cDNA fragments were amplified via PCR. The sample libraries were mixed in equimolar amounts 200-500 bp fragments were purified from an agarose gel after electrophoresis. The libraries were sequenced on an Illumina Nextseq 500 system with a read length of 75 bp. Fastq files were deposited in Array Express (accession number E-MTAB-8582).

*in vitro transcription assays*

*In vitro* transcription reactions used the method of Kolb *et al.*[4] as described by Savery *et al.*[5]. Plasmid template DNA was isolated from *E. coli* transformed with pSR containing the appropriate promoter DNA fragment. Reaction buffer contained 20 mM Tris pH 7.9, 200 mM GTP/ATP/CTP, 10 mM UTP, 5 μCi ($\alpha^{32}$P) UTP, 500 mM DTT, 5 mM MgCl2, 100 μg $ml^{-1}$ BSA and 0.2 mM cAMP. Template DNA (at a final concentration of 16 μg $ml^{-1}$) was incubated with RNA polymerase holoenzyme to start the reaction.

*β-galactosidase assays*

Assays were done according to the method of Miller >. Cells were grown in LB media supplemented with appropriate antibiotics to mid-log phase. Values shown are the mean of three independent experiments. Error bars represent the standard deviation of three independent experiments. Promoters were characterised as active if they stimulated β-galactosidase activity >2-fold over background levels generated by promoterless *lacZ*.

*Bioinformatics*

Individual sequence reads were mapped against the *B. subtilis* genome (Genbank accession number NC_000964.3) using Bowtie2[7]. Resulting Binary Alignment Map (BAM) files were used to generate wiggle plots using bam2wig.py[8,9]. For each strand of the chromosome, we assigned TSSs to base positions where the read depth increased more than 3-fold, compared to the previous base, in both experimental replicates. To compare TSSs in wild type *E. coli*, and the Δ*hns* derivative, we used our previously generated data[10] and remapped TSSs. This was done using TSSpredator (version 1.06)[11] with the following settings: step height 0.1, step height reduction 0.09, step factor 1.5, step factor reduction 0.5, enrichment factor 3, normalisation percentile 0.9, enrichment normalisation percentile 0.5, UTR length 300 and antisense UTR length 100. Cluster method was set to HIGHEST and all other parameters were set to 0.

Bidirectional promoters were identified by the distance between each promoter on the top DNA strand and the nearest promoter on the bottom DNA strand. Promoters were classified as bidirectional if the distance between divergent start sites was between 10 and 25 bp. To determine the distance between TSSs and promoter -10 elements were searched for the sequence TANNNT in the 17 bp region upstream of the TSS. If this sequence did not occur, or occurred multiple times, the TSS was excluded to avoid ambiguities. To generate DNA sequence motifs we used Weblogo[12]. For directional *E. coli* promoters we created two alignments, anchored by either the position of the TSS or -10 element, that were then spliced together in the intervening DNA. This was required because the spacing between the +1 and -10 entities is variable (Figure S4a) and results in improper alignment unless taken into account (compare Figure 2c and S4b). This adjustment was not required for bidirectional promoters with TSSs separated by 18 bp (Figure 2b). In this situation, juxtaposition of the TSSs and -10 elements are "locked" in place in accordance with Figure 3 and the associated description. We identified horizontally acquired genes with DarkHorse using genus level phylogenetic granularity[13]. Sections of DNA with high or low H-NS binding were identified using the ChIP-seq analysis of Kahramanoglou *et al*[14].

*Analysis of symmetry scores*

To determine symmetry scores, we derived a PWM corresponding to sequences from -100 to +50 bp relative to each TSSs for each species test. We refer to this as the “forward PWM”. (Note that for the heatmap in Figure 2a, the forward PWM was derived from sequences from -100 to +100 to facilitate analysis over a longer range of spacings; importantly, this does not impact the calculated scores). We then made a “reverse PWM” that corresponds to the reverse complement of the forward PWM, but was limited to sequences from -37 to +5 relative to the TSSs, since this is the range that includes all key promoter elements for all species tested. We aligned the forward and reverse PWMs across all possible spacings. For each spacing, we calculated a symmetry score by (i) multiplying the fraction of each of the four nucleotides at each position of the forward PWM with the fraction of each of the complementary nucleotides at the overlapping position of the reverse PWM, and (ii) multiplying this value by 4, taking the log (base 2), and summing for all positions within the overlapping PWM positions.

Symmetry scores were also calculated for individual *E. coli* promoter sequences, to compare promoter sequences in horizontally acquired versus non-horizontally acquired regions, and to compare promoter sequences in H-NS-bound versus unbound regions. In these cases, we analysed individual promoter regions from position -100 to +50 relative to the TSS. We aligned the reverse PWM for *E. coli* (derived as described above) with each promoter sequence across all possible spacings. For each spacing, we determined the frequency of the nucleotide found in the promoter with the corresponding nucleotide frequency in the reverse PWM. We then multiplied these values for every position within the PWM. The final symmetry score for each promoter sequence was calculated as the maximum score across all possible spacings multiplied by a constant (to avoid extremely small numbers).
