## Supplementary Figures and Legends for "Widespread divergent transcription from prokaryotic promoters"

### Figure S1

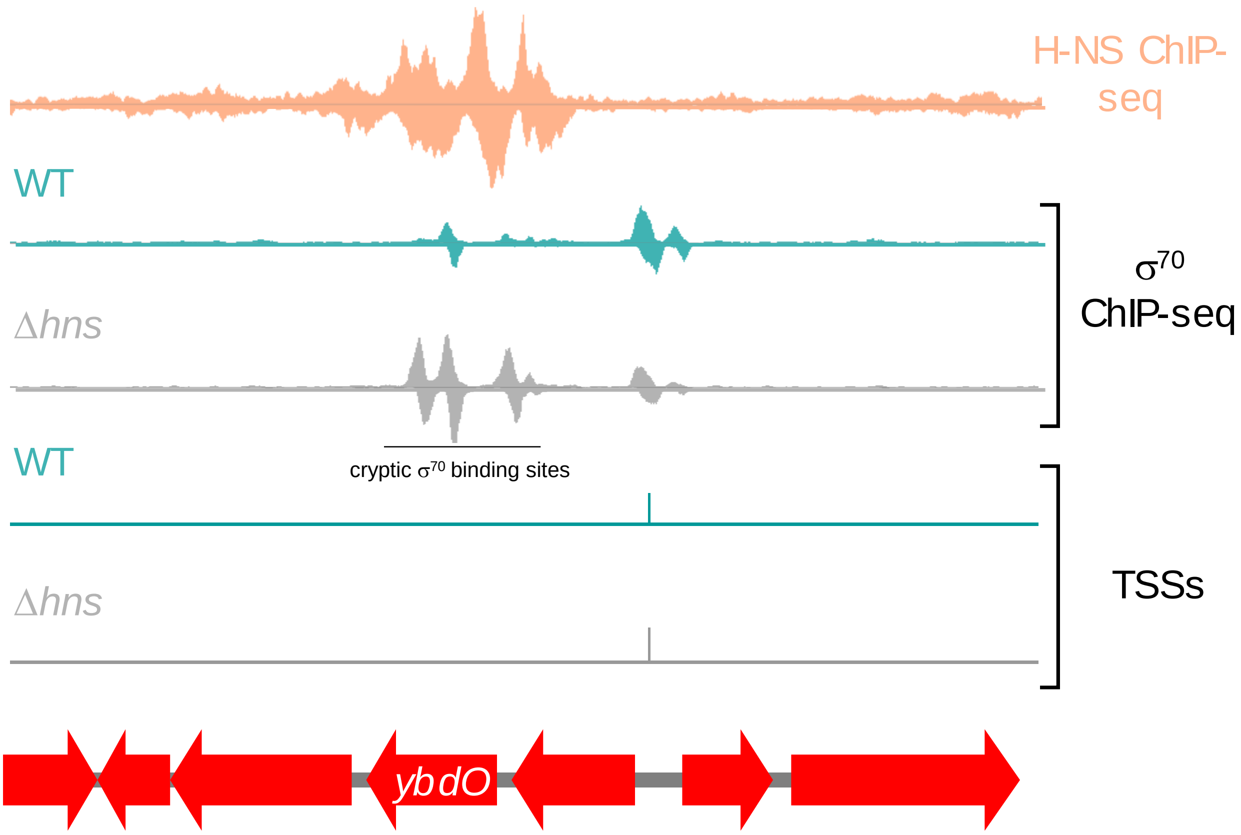
H-NS ChIP-seq
WT
Δhns
cryptic σ70 binding sites
σ70 ChIP-seq
WT
Δhns
TSSs
ybdO

### Figure S2

***yibA2* in "a" orientation***

-35 -10

gggttttatctgttttatgcgatgagttaaaaaaaatactgtttatgatgatatcattg

cccaaaatagacaaaatacgctactcaatttttttttatgacaaatactactatagtaac

aagctgcgggtgaattaggtgataaaacgctacttcctgttttagatactatgttgtaca

ttcgacgcccacttaatccactatttttgcgatgaaggacaaaatctatgatacaacatgt

-10 -35

agtttgatgacaatgaaattataacttccgaggaggaagcttttgtcagtgcgcaaaaag

tcaaactactgttactttaatattgaaggctcctccttcgaaaacagtcacgcgtttttc

D49724 (oligo for primer extension)

atcctgaatttcaggctcagttcaacgacctgctgaaaaactatgccgggcgtccaaccg

taggacttaaagtccgagtcaagttgctggacgacttttgatacggcccgcaggttggc

cgctgaccaaatgccagaacattacagccgggacga 3′

gcgactggtttacggtcttgtaatgtcggccctgct 5′

***wzxB* in "a" orientation***

-35

5′ acgttactttatctttactatctgctgctttggcaatactctgagttgctgtgagattga

3′ tgcaatgaaatagaaatgatagacgacgaaaccgttatgagactcaacgacactctaact

-10

ggg

aaccatattcaacaaacattatcatatatagcatagtcgcttggcaaaaaccgaatatac

ttggtataagttgtttgtaatagtatatatcgtatcagcgaaccgtttttggcttatatg

ccc -10 -35

cgaaattttcaggaccaagtgttcttacaaaggaggaagcttttgtcagtgcgcaaaaag

gctttaaaagtcctggttcacaagaatgtttcctccttcgaaaacagtcacgcgtttttc

D49724 (oligo for primer extension)

atcctgaatttcaggctcagttcaacgacctgctgaaaaactatgccgggcgtccaaccg

taggacttaaagtccgagtcaagttgctggacgacttttgatacggcccgcaggttggc

cgctgaccaaatgccagaacattacagccgggacga 3′

gcgactggtttacggtcttgtaatgtcggccctgct 5′

***yigG* in "a" orientation***

-35 -10

5′ cattgcctgaacaggcaaaatcttcggctatcattgtgatgatagagatgatatatactg

3′ gtaacggacttgtccgttttagaagccgatagtaacactactatctctactatatatgac

cc

ctaatgtaccaaaaacataagtttttatatagatgaaaccactatcacggagtcgctggc

gattacatggttttgtattcaaaaatatatctactttggtgatagtgcctcagcgaccg

gg -10 -35

aattcatgttgatgacgagataatggagtaaggaggaagctttgtcagtgcgcaaaaag

ttaagtacaactactgctctattacctcattcctccttcgaaacagtcacgcgtttttc

D49724 (oligo for primer extension)

atcctgaatttcaggctcagttcaacgacctgctgaaaaactatgccgggcgtccaaccg

taggacttaaagtccgagtcaagttgctggacgactttttgatacggcccgcaggttggc

cgctgaccaaatgccagaacattacagccgggacga 3′

gcgactggtttacggtcttgtaatgtcggccctgct 5′

***yqiI2* in "a" orientation***

5′ gaatattttatgaatgttttctgtaataatgcactaccaggcccatctccaggagaagaa

3′ cttataaaatacttacaaaagacattattacgtgatggtccgggtagaggtcctcttctt

-35 Δ -10

tacccatctgcatgggcaaatataatgatgttgttagcatcaggtcaagactttttataat

atgggtagacgtacccgtttatattactacaacaatcgtagtccagttctgaaaatatta

-10 -35

caaaactcttatactttcggtgtaacttataggaggaagctttgtcagtgcgcaaaaag

gttttgagaatatgaaagccacattgaatatcctccttcgaaacagtcacgcgtttttc

D49724 (oligo for primer extension)

atcctgaatttcaggctcagttcaacgacctgctgaaaaactatgccgggcgtccaaccg

taggacttaaagtccgagtcaagttgctggacgactttttgatacggcccgcaggttggc

cgctgaccaaatgccagaacattacagccgggacga 3′

gcgactggtttacggtcttgtaatgtcggccctgct 5′

***ygaQ1*** **in “a” orientation***

-35

cggttacacaatactaacttatttaacccaaaatatcataaaaaagccgtt**atgaat**tac

gccaatgtgttatgattgaataa**a**ttggg**ttttat**a**gt**atttttcggcaat**acttaa**tg

-10 -35

-10

atggaatatc**tg**g**taactt**gtcagtt**g**gatgaacaacaaatgtcatcactgctttatgaa

taccttatagaccattgaacagtcaacctacttgttgtttacagtagtgacgaaatactt

agagatgatttaagcgccattgatttttcaaggagg*aagctt*ttgtcagtgcgcaaaaag

tctctactaaattcgcggtaactaaaaagttcctcc*ttcgaa*aacagtcacgcgtttttc

D49724 (oligo for primer extension)

atcctgaatttcaggctcagtt**caacgacctgctgaaaaactatgccgggcgtccaacc**g

taggacttaaagtccgagtcaagttgctggacgactttttgatacggcccgcaggttggc

cgctgaccaaatgccagaacattacagccgggacga 3’

gcgactggtttacggtcttgtaatgtcggccctgct 5’

* For the “b” orientation read the reverse complement sequence of the bases in black. The sequences in grey remain unchanged

### Figure S3

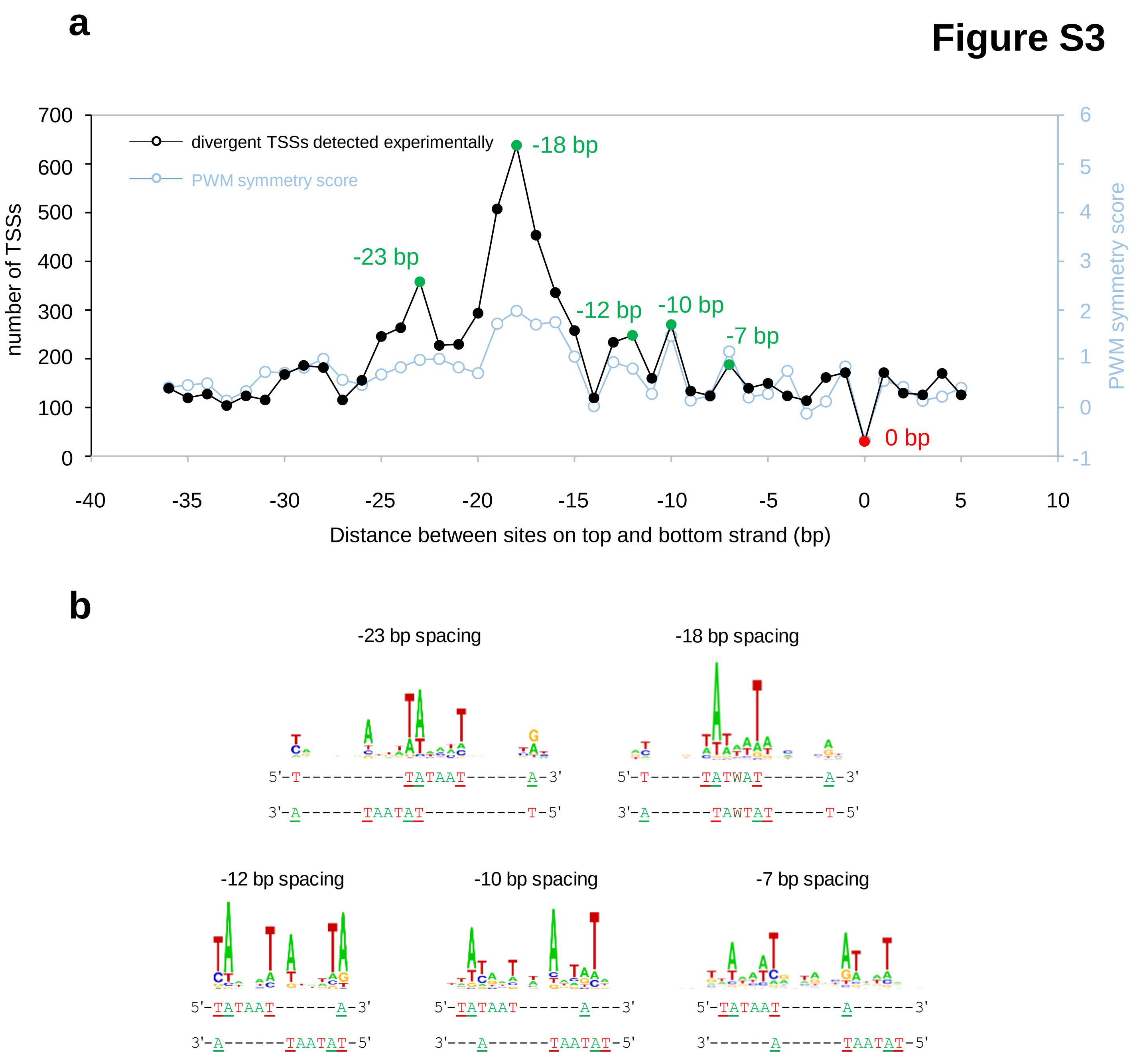
a
divergent TSSs detected experimentally
PWM symmetry score
number of TSSs
PWM symmetry score
Distance between sites on top and bottom strand (bp)
-18 bp
-23 bp
-12 bp
-10 bp
-7 bp
0 bp
b
-23 bp spacing
5'-T----------TATAAT------A-3'
3'-A------TAATAT----------T-5'
-18 bp spacing
5'-T-----TATWAT------A-3'
3'-A------TAWTAT-----T-5'
-12 bp spacing
5'-TATAAT------A-3'
3'-A------TAATAT-5'
-10 bp spacing
5'-TATAAT------A---3'
3'---A------TAATAT-5'
-7 bp spacing
5'-TATAAT------A------3'
3'------A------TAATAT-5'

### Figure S4

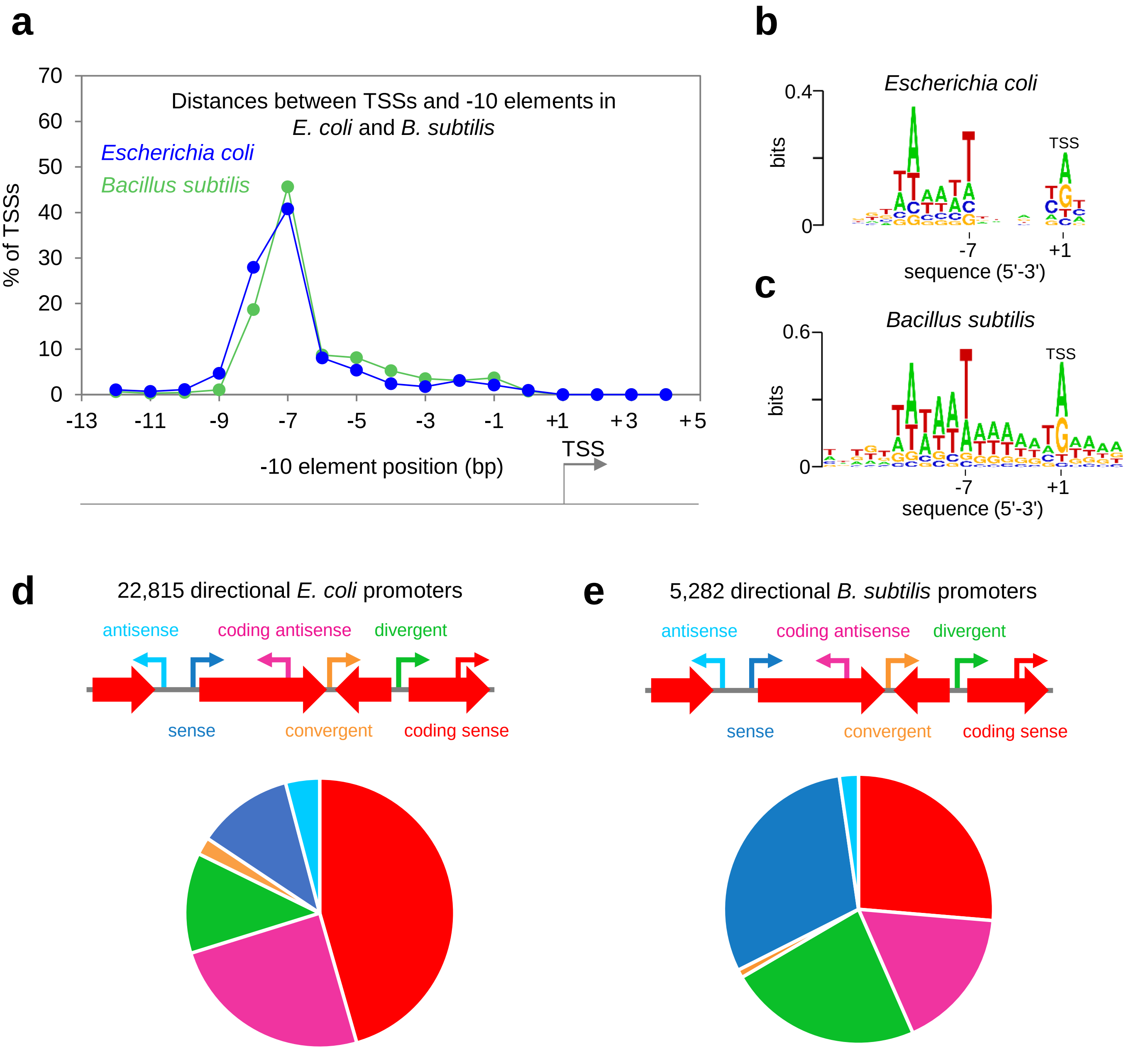
a
b
c
d
e
Distances between TSSs and -10 elements in
E. coli and B. subtilis
Escherichia coli
Bacillus subtilis
% of TSSs
70
60
50
40
30
20
10
0
-13
-11
-9
-7
-5
-3
-1
+1
+3
+5
-10 element position (bp)
TSS
Escherichia coli
0.4
bits
0
TSS
-7
+1
sequence (5'-3')
Bacillus subtilis
0.6
bits
0
TSS
-7
+1
sequence (5'-3')
22,815 directional E. coli promoters
antisense
coding antisense
divergent
sense
convergent
coding sense
5,282 directional B. subtilis promoters
antisense
coding antisense
divergent
sense
convergent
coding sense

### Figure S5

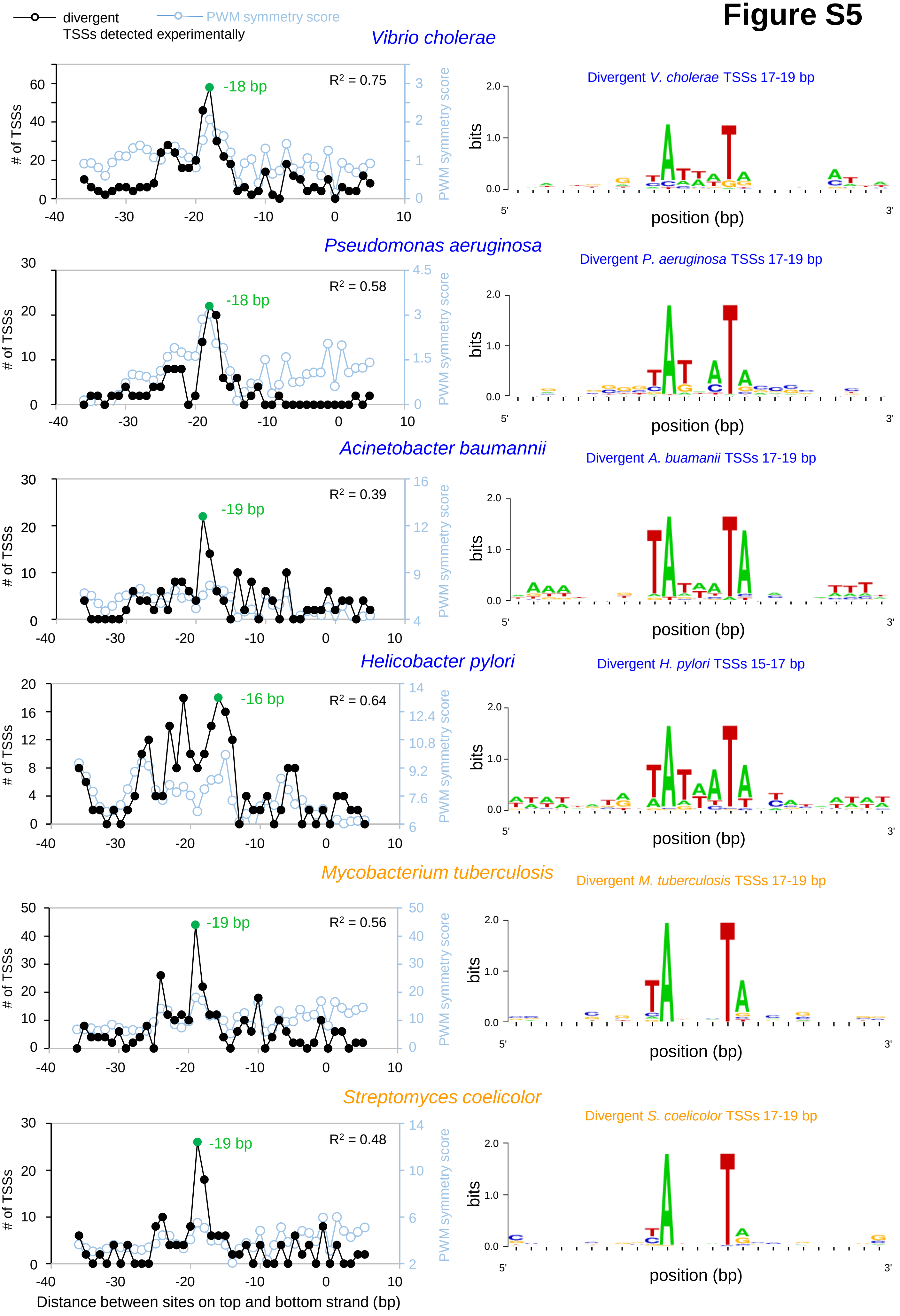
divergent
TSSs detected experimentally
PWM symmetry score
Vibrio cholerae
R² = 0.75
-18 bp
Divergent V. cholerae TSSs 17-19 bp
Pseudomonas aeruginosa
R² = 0.58
-18 bp
Divergent P. aeruginosa TSSs 17-19 bp
Acinetobacter baumannii
R² = 0.39
-19 bp
Divergent A. buamanii TSSs 17-19 bp
Helicobacter pylori
R² = 0.64
-16 bp
Divergent H. pylori TSSs 15-17 bp
Mycobacterium tuberculosis
R² = 0.56
-19 bp
Divergent M. tuberculosis TSSs 17-19 bp
Streptomyces coelicolor
R² = 0.48
-19 bp
Divergent S. coelicolor TSSs 17-19 bp
### of TSSs
PWM symmetry score
bits
position (bp)
5'
3'
Distance between sites on top and bottom strand (bp)

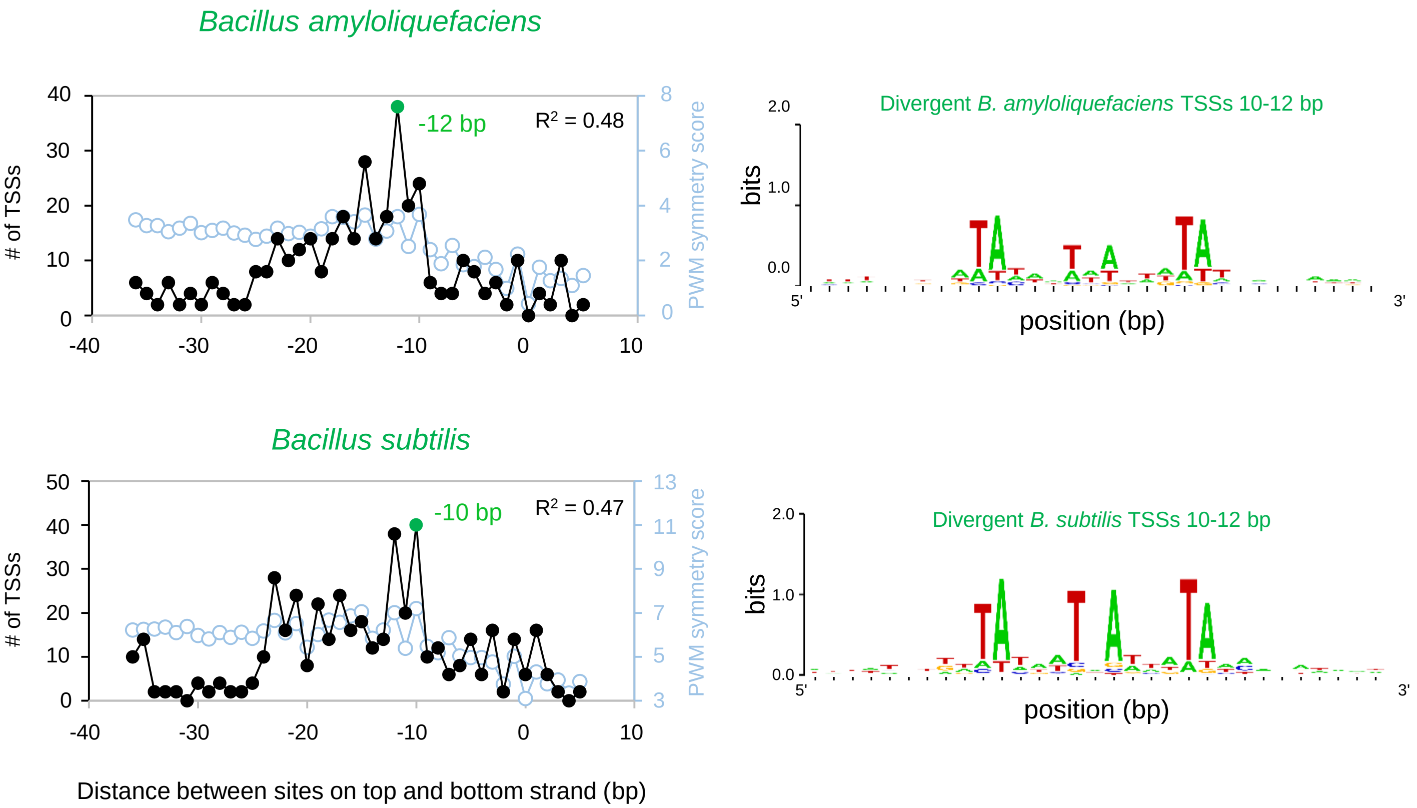

Bacillus amyloliquefaciens
-12 bp
R2 = 0.48
### of TSSs
PWM symmetry score
Divergent B. amyloliquefaciens TSSs 10-12 bp
bits
position (bp)
Bacillus subtilis
-10 bp
R2 = 0.47
Divergent B. subtilis TSSs 10-12 bp
Distance between sites on top and bottom strand (bp)

### Figure S6

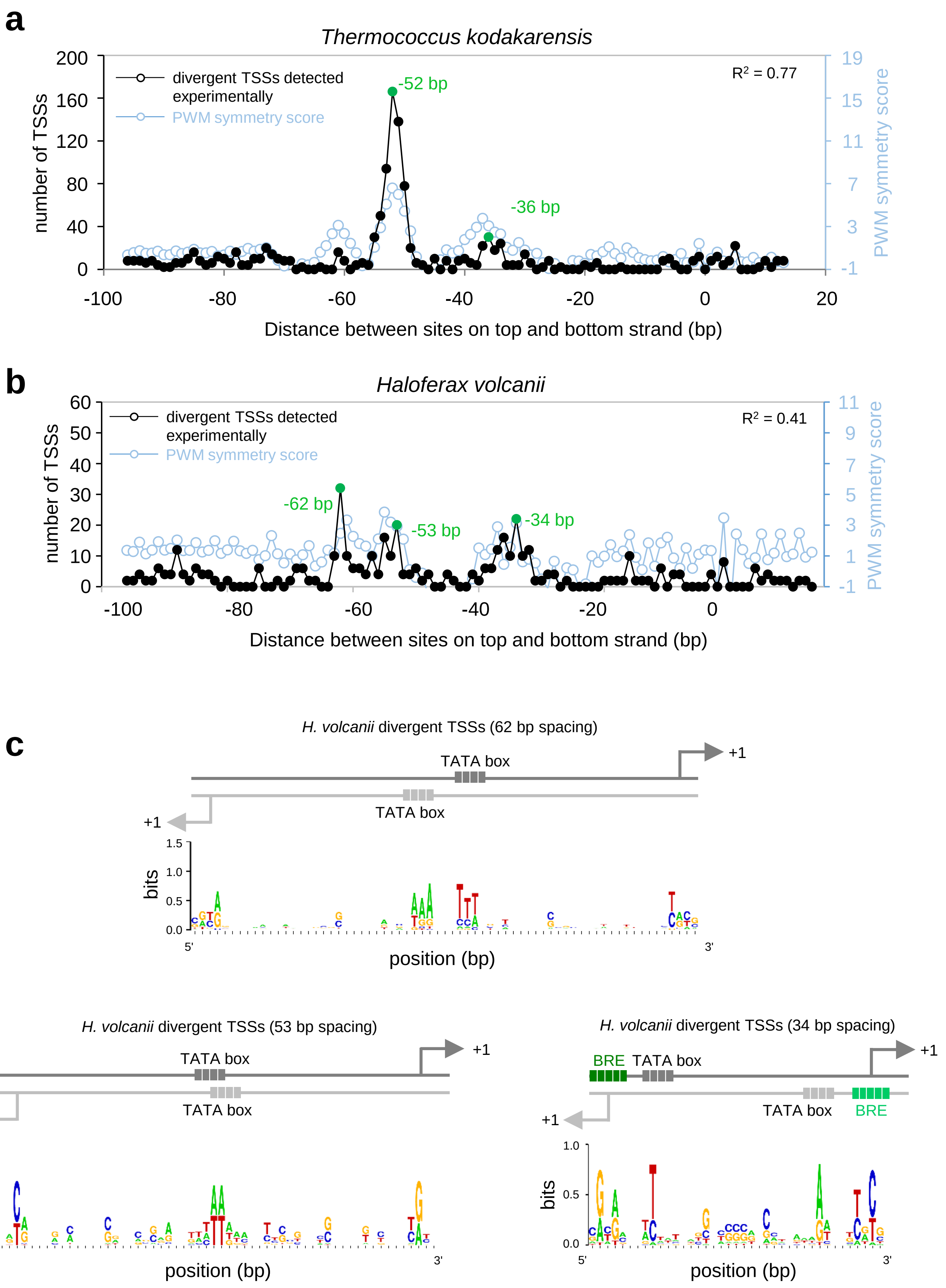

a
Thermococcus kodakarensis
divergent TSSs detected experimentally
PWM symmetry score
R² = 0.77
-52 bp
-36 bp
number of TSSs
PWM symmetry score
200
160
120
80
40
0
19
15
11
7
3
-1
-100
-80
-60
-40
-20
0
20
Distance between sites on top and bottom strand (bp)
b
Haloferax volcanii
divergent TSSs detected experimentally
PWM symmetry score
R² = 0.41
-62 bp
-53 bp
-34 bp
number of TSSs
PWM symmetry score
60
50
40
30
20
10
0
11
9
7
5
3
1
-1
-100
-80
-60
-40
-20
0
Distance between sites on top and bottom strand (bp)
c
H. volcanii divergent TSSs (62 bp spacing)
TATA box
TATA box
+1
+1
bits
1.5
1.0
0.5
0.0
5'
3'
position (bp)
H. volcanii divergent TSSs (53 bp spacing)
TATA box
TATA box
+1
+1
bits
1.5
1.0
0.5
0.0
5'
3'
position (bp)
H. volcanii divergent TSSs (34 bp spacing)
BRE
TATA box
TATA box
BRE
+1
+1
bits
1.0
0.5
0.0
5'
3'
position (bp)

### Figure S7

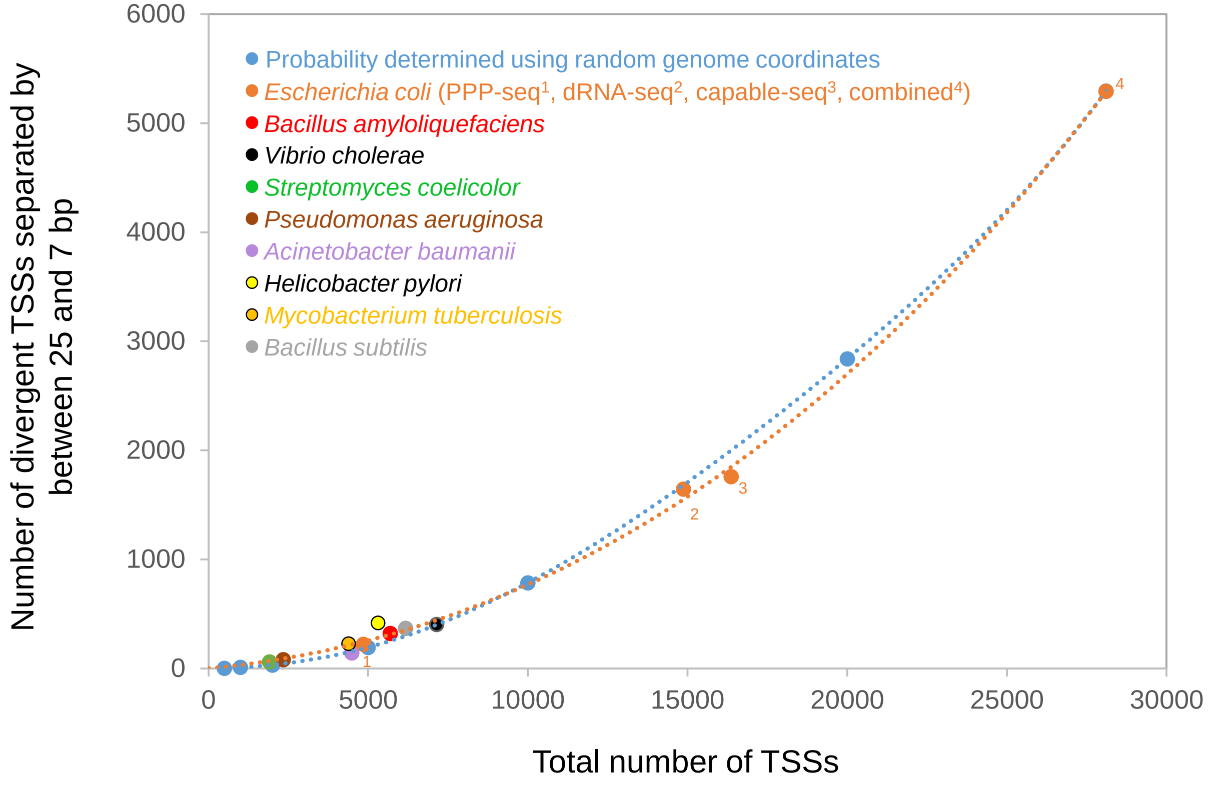
Probability determined using random genome coordinates
Escherichia coli (PPP-seq1, dRNA-seq2, capable-seq3, combined4)
Bacillus amyloliquefaciens
Vibrio cholerae
Streptomyces coelicolor
Pseudomonas aeruginosa
Acinetobacter baumanii
Helicobacter pylori
Mycobacterium tuberculosis
Bacillus subtilis
Number of divergent TSSs separated by between 25 and 7 bp
Total number of TSSs
0
1000
2000
3000
4000
5000
6000
5000
10000
15000
20000
25000
30000
1
2
3
4

### Figure S8

**a**

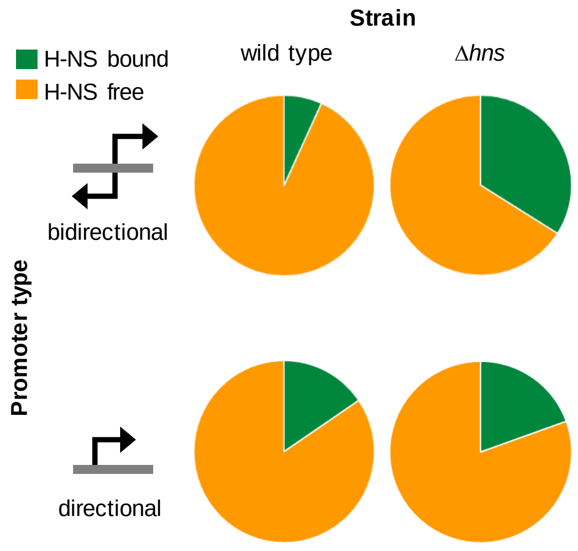
Strain
wild type
Δhns
H-NS bound
H-NS free
Promoter type
bidirectional
directional

**b**

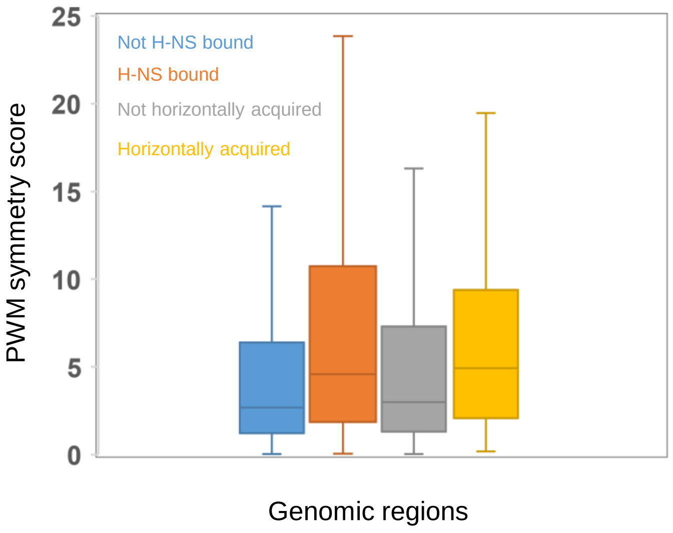
PWM symmetry score
25
20
15
10
5
0
Not H-NS bound
H-NS bound
Not horizontally acquired
Horizontally acquired
Genomic regions

### Figure S9

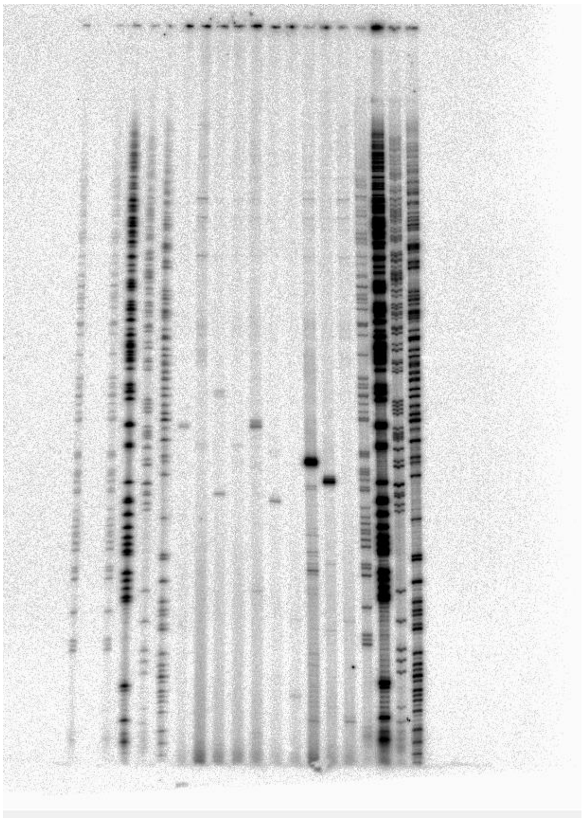

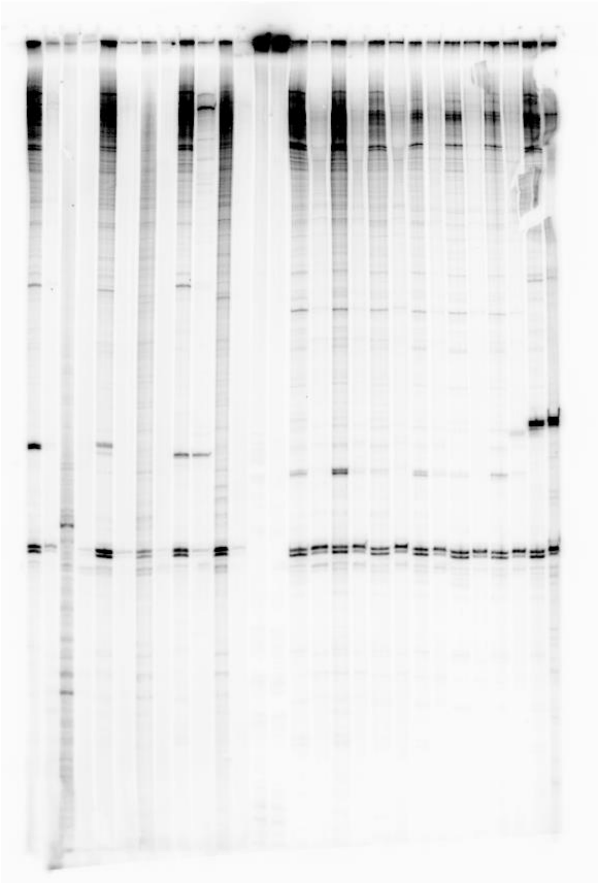

#### SUPPLEMENTARY FIGURE LEGENDS

**Figure S1: Not all $\sigma^{70}$ binding sites align with an RNA 5' end.** Binding patterns for H-NS (peach) and $\sigma^{70}$ (teal or grey) are derived from ChIP-seq assays [1]. The RNA 5' ends associated with $\sigma^{70}$ binding (teal or grey) were identified by PPP-seq [1]. Genes are shown as red block arrows.

**Figure S2: Sequences of cryptic RNA polymerase binding sites associated with divergent transcription.** The figure shows promoter DNA sequences (black typeface) and part of the plasmid DNA backbone (grey typeface). The promoter -10 (red) and -35 (green) elements are highlighted on each DNA strand and transcription start sites are denoted by a bent arrow. Sites of mutations and deletions (Δ) are boxed. The sequences are in the "a" orientation as indicated in Figure 1. When in the "b" orientation the DNA sequence encompassed by black typeface is the reverse complement. Oligonucleotide D49724, used in primer extension analysis, is indicated by a half arrow and binds to the corresponding sequence in grey bold typeface.

**Figure S3: Spacing optima between divergent transcription start sites.** a) The graph shows the number of divergent TSSs separated by different distances. The majority of bottom strand transcription start sites occur 18 bp upstream of top strand RNA initiation sites. However, peaks in the occurrence of divergent TSSs also occur elsewhere. These positions are denoted by green data points. The red data point indicates a sharp decrease in the occurrence of divergent TSSs. The symmetry score increases at spacing intervals where the promoter PWM identifies matching sequences overlapping on each DNA strand.  b) DNA sequence motifs associated with each preferred spacing between divergent TSSs. Motifs were generated by aligning sequences according to the top strand TSS. The configuration of promoter -10 elements and TSSs, indicated by each motif, is shown below the respective sequence logo. Key positions within the -10 elements, and TSSs, are underlined.

**Figure S4: Properties of transcription start sites in *Bacillus subtilis*.** We mapped transcription start sites globally in *Bacillus subtilis* using cappable-seq. The general properties of *B. subtilis* promoters were compared with those identified in *E. coli*.  a) Distances between promoter -10 elements and transcription start sites (TSSs) in *Escherichia coli* and *B. subtilis*. b) DNA sequence motifs associated with unidirectional TSSs in *E. coli* and *B. subtilis*. d,e) Positioning of TSSs with respect to coding DNA sequences in *E. coli* and *B. subtilis*.

**Figure S5: Properties of bidirectional promoters in different bacteria.** The figure shows the number of divergent transcription start sites separated by different distances (black line in graph). The data point in each graph, corresponding to the preferred configuration of divergent start sites, is green. The pale blue data indicate predicted promoter overlap (i.e. symmetry) derived from a position weight matrix (PWM) search of each DNA strand. The $R^2$ values indicate the degree of correlation between computational prediction and experimental data shown. The DNA motifs adjacent to each graph were generated by aligning promoter -10 hexamer sequences. For *V. cholerae, P. aeruginosa, A. baumanii, M. tuberculosis,* and *S. coelicolour* we aligned -10 elements from start sites separated by 17, 18 and 19 bp. In the case of *H. pylori,* we aligned -10 elements from those start sites 15, 16 or 17 bp apart. Note that all of these distances typically involve the same configuration of -10 elements because the distance between the -10 hexamer and transcription start site is variable (see Figure S4). For *B. subtilis* and *B. amyloliquefaciens* we aligned start sites separated by 10, 11 or 12 bp.

**Figure S6: Bidirectional promoters in the archaea *Thermococcus kodakarensis* and *Haloferax volcanii* have a shared TATA box.** a,b) The figure shows the number of divergent transcription start sites separated by different distances (black line in graph). The data points in each graph, corresponding to the preferred configurations of divergent start sites, are green. The pale blue data indicate promoter sequence symmetry. The $R^2$ values indicate the degree of correlation between computational prediction and experimental data shown. b) DNA sequences associated with divergent TSSs in *Haloferax volcanii* separated by 62, 53 or 34 bp were aligned according to the position of the TSS on the top DNA strand. The inferred configuration of key elements is shown above each motif.

**Figure S7: The ratio of directional to bidirectional promoters is similar in different bacteria.** We used multiple *Escherichia coli* TSS maps to identify bidirectional promoters (corresponding to divergent TSSs separated by between 25 and 7 bp). We noticed that the proportion of TSSs from bidirectional promoters was much smaller in datasets with fewer total TSSs. We reasoned that this was logical; the chance of detecting both transcripts from a given bidirectional promoter is much smaller for less complete TSS maps. For instance, 19 % of TSSs in the combined *E. coli* TSS map (28,107 TSSs in total) were derived from a bidirectional promoter. In contrast, this value was only 5 % for TSSs identified by PPP-seq[1] (4,846 TSSs in total). The number of total and divergent TSSs, for each *E. coli* TSS map, is plotted in orange; the relationship is not linear. For comparison, we generated a probability model using a mock TSS map for *E. coli*. The artificial map consisted of 28,107 randomly selected *E. coli* genome co-ordinates as TSSs. Of these, 19 % of positions on the bottom strand were set to be between 7 and 25 bp upstream of a top strand co-ordinate (i.e. the mock data exactly emulated the combined TSS composition of the genuine experimental data for *E. coli*). We then randomly selected sub-populations of genome co-ordinates from the mock TSS map and determined how many pairs of top and bottom stand positions remained separated by between 7 and 25 bp. These data are plotted in pale blue. Consistent with our logic, the relationship was not linear and resembled the real experimental data in orange. Finally, we plotted experimentally determined TSS maps for different bacteria (all TSS numbers were normalised for genome size). Crucially, these organisms have been subject to much less scrutiny than *E. coli*. Hence, the total number of TSSs identified for each bacterium is comparatively small. Even so, it is clear that all data points fall close to the orange and pale blue trend lines. Hence, the fraction of promoters that are bidirectional must be broadly similar in different bacterial species.

**Figure S8: Bidirectional promoters are enriched in horizontally acquired genes.** a) Detection of directional and bidirectional promoters by PPP-seq[1]. The pie charts show the fractions of each promoter type detected in H-NS bound or H-NS free regions of the *E. coli* genome in the presence and absence of H-NS. b) Distribution of position weight matrix (PWM) scores for bidirectional promoters in different sections of the *E. coli* genome. Higher scores indicate a better match to the PWM describing bidirectional promoters.

**Figure S9: Raw gel images.** Full uncropped images of gel used in this work.
