## Supplementary Tables for "Widespread divergent transcription from prokaryotic promoters"

**Table S1: Positions of TSSs across the Bacillus subtilis genome (NC_000964.3).**

| Strand | Position |
|---|---|
| + | 155 |
| + | 157 |
| - | 288 |
| - | 910 |
| - | 3168 |
| + | 3176 |
| + | 3177 |
| + | 3180 |
| + | 3649 |
| + | 3652 |
| - | 4364 |
| - | 4365 |
| - | 4810 |
| + | 6916 |
| + | 6918 |
| - | 7778 |
| - | 9123 |
| + | 9537 |
| + | 9540 |
| + | 9634 |
| + | 9637 |
| + | 11065 |
| + | 11643 |
| + | 11711 |
| + | 14671 |
| + | 14692 |
| + | 15879 |
| + | 15884 |
| - | 17359 |
| + | 17462 |
| + | 17465 |
| + | 18977 |
| + | 18979 |
| + | 19022 |
| - | 20584 |
| + | 20597 |
| + | 20599 |
| + | 22281 |
| + | 22284 |
| - | 23012 |
| + | 23121 |
| - | 23812 |
| - | 24473 |
| + | 24574 |
| + | 24667 |
| - | 25403 |
| + | 25667 |

| | |
|---|---|
| + | 25668 |
| - | 25794 |
| + | 25804 |
| + | 26377 |
| + | 26740 |
| + | 26788 |
| - | 27200 |
| - | 27544 |
| - | 27549 |
| + | 28275 |
| + | 28382 |
| + | 28385 |
| + | 28708 |
| + | 30168 |
| + | 31533 |
| + | 32111 |
| + | 32179 |
| + | 35138 |
| + | 35237 |
| + | 35802 |
| + | 35805 |
| + | 36549 |
| + | 36737 |
| + | 36739 |
| + | 37690 |
| + | 37692 |
| + | 38556 |
| + | 40673 |
| + | 41328 |
| + | 41329 |
| - | 41869 |
| + | 42435 |
| + | 43196 |
| + | 43509 |
| - | 45207 |
| - | 45212 |
| - | 45398 |
| + | 45527 |
| + | 47526 |
| + | 48617 |
| + | 48686 |
| + | 49999 |
| + | 50002 |
| - | 50013 |
| - | 50015 |
| + | 50420 |
| + | 50754 |
| - | 52597 |
| + | 52748 |
| - | 52910 |

| | |
|---|---|
| + | 53483 |
| + | 54408 |
| + | 54411 |
| + | 54412 |
| + | 55094 |
| - | 55750 |
| + | 55830 |
| + | 55831 |
| + | 56322 |
| + | 56324 |
| + | 57327 |
| - | 57500 |
| + | 58744 |
| + | 58746 |
| + | 58748 |
| + | 58878 |
| - | 59392 |
| + | 59440 |
| + | 59450 |
| + | 59451 |
| + | 60257 |
| + | 60759 |
| - | 62019 |
| + | 63995 |
| + | 64752 |
| - | 66302 |
| + | 66325 |
| + | 67079 |
| + | 67082 |
| + | 67410 |
| + | 68458 |
| + | 68789 |
| + | 68790 |
| + | 69053 |
| + | 69055 |
| + | 69581 |
| + | 69583 |
| - | 69618 |
| + | 70153 |
| + | 70155 |
| + | 71009 |
| - | 71031 |
| + | 71818 |
| - | 72580 |
| + | 72841 |
| + | 73064 |
| + | 74891 |
| + | 74893 |
| - | 75745 |
| + | 75761 |

| | |
|---|---|
| + | 76313 |
| + | 76672 |
| + | 76950 |
| + | 76954 |
| - | 78399 |
| + | 79040 |
| + | 79162 |
| - | 79617 |
| + | 80659 |
| + | 81243 |
| + | 81731 |
| + | 82073 |
| + | 82833 |
| - | 82986 |
| - | 83262 |
| + | 84068 |
| + | 84070 |
| + | 85064 |
| - | 85799 |
| + | 87267 |
| + | 87270 |
| + | 87387 |
| + | 87467 |
| + | 87817 |
| + | 87897 |
| - | 87915 |
| - | 87917 |
| + | 88203 |
| - | 89570 |
| + | 90421 |
| + | 90424 |
| + | 91790 |
| + | 92156 |
| + | 92256 |
| + | 95216 |
| + | 95237 |
| + | 96146 |
| + | 96276 |
| + | 97646 |
| + | 98012 |
| + | 98112 |
| + | 101072 |
| + | 101093 |
| + | 101294 |
| + | 101295 |
| + | 102004 |
| + | 103138 |
| + | 106008 |
| + | 106200 |
| + | 106583 |

| | |
|---|---|
| + | 106585 |
| + | 106586 |
| + | 107626 |
| + | 107830 |
| + | 108466 |
| + | 108598 |
| - | 108753 |
| + | 109205 |
| + | 109206 |
| + | 109207 |
| + | 109941 |
| + | 110295 |
| - | 110767 |
| + | 110994 |
| + | 110998 |
| + | 112238 |
| + | 114482 |
| + | 114524 |
| + | 114598 |
| + | 115551 |
| + | 116571 |
| + | 116575 |
| - | 117292 |
| + | 117310 |
| + | 117736 |
| + | 117739 |
| - | 118451 |
| + | 118469 |
| + | 118470 |
| + | 119845 |
| + | 119848 |
| - | 121643 |
| + | 121704 |
| + | 121705 |
| + | 121842 |
| - | 122263 |
| - | 123052 |
| + | 123753 |
| + | 123755 |
| + | 129077 |
| + | 129298 |
| + | 129299 |
| + | 129301 |
| - | 132777 |
| + | 132799 |
| + | 132942 |
| + | 134147 |
| + | 135168 |
| + | 135169 |
| + | 135238 |

| | |
|---|---|
| - | 135497 |
| + | 140982 |
| + | 141982 |
| - | 142368 |
| + | 143866 |
| + | 144021 |
| - | 144965 |
| + | 145552 |
| + | 146571 |
| + | 146716 |
| + | 147265 |
| + | 147759 |
| + | 147761 |
| - | 148022 |
| + | 149470 |
| + | 149519 |
| + | 149908 |
| + | 152068 |
| + | 152795 |
| + | 153221 |
| + | 153732 |
| + | 153735 |
| + | 153737 |
| + | 154847 |
| + | 155122 |
| + | 155124 |
| + | 155135 |
| - | 155531 |
| - | 156233 |
| + | 156431 |
| - | 157380 |
| + | 157385 |
| + | 157391 |
| - | 158940 |
| + | 159101 |
| - | 159120 |
| + | 160036 |
| + | 160686 |
| + | 162146 |
| + | 162512 |
| + | 162612 |
| + | 165570 |
| + | 165591 |
| + | 165707 |
| + | 165750 |
| + | 166253 |
| + | 166384 |
| + | 167754 |
| + | 168120 |
| + | 168220 |

| | |
|---|---|
| + | 171176 |
| + | 171197 |
| + | 171382 |
| + | 172751 |
| + | 173116 |
| + | 173216 |
| + | 176197 |
| + | 176606 |
| + | 176930 |
| + | 177109 |
| + | 178594 |
| + | 178597 |
| - | 179682 |
| + | 181986 |
| + | 181988 |
| - | 182315 |
| - | 183358 |
| - | 185047 |
| + | 186121 |
| + | 186638 |
| + | 188529 |
| + | 191968 |
| + | 192351 |
| - | 192998 |
| - | 192999 |
| + | 193035 |
| + | 193222 |
| + | 193341 |
| + | 193345 |
| - | 194052 |
| + | 194182 |
| + | 194183 |
| + | 194775 |
| + | 194777 |
| + | 194821 |
| - | 195283 |
| + | 196140 |
| + | 197216 |
| + | 199857 |
| + | 199967 |
| - | 200017 |
| + | 200267 |
| + | 200600 |
| + | 201521 |
| + | 201873 |
| - | 203507 |
| + | 203608 |
| + | 204331 |
| + | 204991 |
| - | 206251 |

| | |
|---|---|
| + | 209904 |
| + | 210516 |
| + | 210766 |
| + | 211425 |
| + | 211427 |
| + | 211429 |
| - | 211603 |
| + | 212080 |
| + | 212414 |
| + | 213890 |
| + | 216339 |
| - | 217372 |
| + | 219511 |
| - | 220159 |
| + | 220169 |
| + | 220602 |
| + | 221098 |
| + | 221201 |
| + | 222110 |
| + | 222598 |
| + | 222951 |
| + | 222958 |
| + | 222959 |
| + | 223389 |
| + | 223720 |
| - | 224013 |
| - | 224149 |
| - | 224950 |
| + | 225015 |
| + | 225034 |
| + | 225037 |
| + | 225457 |
| - | 226057 |
| + | 226438 |
| + | 226439 |
| + | 227346 |
| - | 227770 |
| + | 228917 |
| - | 229376 |
| + | 230449 |
| + | 230641 |
| - | 231115 |
| - | 231116 |
| + | 231270 |
| + | 231309 |
| - | 231532 |
| - | 235481 |
| - | 235483 |
| + | 235604 |
| + | 235935 |

| | |
|---|---|
| - | 238385 |
| + | 238687 |
| - | 239602 |
| + | 239651 |
| - | 239851 |
| - | 240951 |
| + | 241162 |
| - | 241350 |
| - | 241871 |
| + | 242538 |
| - | 242760 |
| + | 243140 |
| - | 243690 |
| + | 243870 |
| + | 243871 |
| + | 245198 |
| - | 246479 |
| + | 246569 |
| + | 246584 |
| + | 246926 |
| + | 246928 |
| + | 247607 |
| + | 247612 |
| + | 247711 |
| + | 248156 |
| - | 248585 |
| + | 248732 |
| + | 249950 |
| + | 249952 |
| + | 251406 |
| + | 252466 |
| + | 252467 |
| + | 252468 |
| - | 254797 |
| + | 256686 |
| + | 256836 |
| + | 257706 |
| + | 257750 |
| + | 258423 |
| + | 258495 |
| + | 258888 |
| + | 259571 |
| + | 261126 |
| - | 261625 |
| - | 262645 |
| + | 264167 |
| + | 265413 |
| + | 266078 |
| + | 266483 |
| + | 267794 |

| | |
|---|---|
| + | 271231 |
| + | 271419 |
| + | 273039 |
| + | 273217 |
| + | 273218 |
| + | 273684 |
| + | 274110 |
| - | 274378 |
| + | 275609 |
| + | 275610 |
| + | 276757 |
| + | 276796 |
| + | 276797 |
| + | 276798 |
| - | 277286 |
| - | 277288 |
| + | 278321 |
| + | 280058 |
| - | 282184 |
| - | 282186 |
| - | 283207 |
| - | 284829 |
| - | 287037 |
| + | 287482 |
| + | 287483 |
| - | 287850 |
| - | 289504 |
| - | 290740 |
| + | 290888 |
| + | 292170 |
| + | 292171 |
| + | 292706 |
| - | 293296 |
| + | 293471 |
| + | 293472 |
| - | 296311 |
| + | 296401 |
| + | 296402 |
| + | 298418 |
| - | 299318 |
| - | 300681 |
| - | 300682 |
| - | 302293 |
| - | 302295 |
| + | 302395 |
| + | 302978 |
| + | 303298 |
| - | 304382 |
| + | 304393 |
| - | 304727 |

| | |
|---|---|
| + | 304777 |
| + | 305629 |
| - | 305999 |
| + | 307561 |
| - | 308055 |
| + | 308272 |
| + | 308273 |
| - | 308451 |
| + | 308615 |
| + | 308921 |
| + | 308923 |
| - | 309231 |
| + | 309419 |
| + | 310657 |
| + | 310982 |
| + | 311870 |
| - | 311909 |
| + | 312069 |
| + | 315420 |
| + | 317694 |
| + | 319333 |
| - | 320708 |
| + | 320942 |
| + | 320989 |
| - | 321681 |
| - | 322064 |
| - | 325266 |
| + | 325281 |
| + | 325283 |
| - | 326004 |
| + | 326114 |
| + | 326597 |
| + | 326854 |
| + | 326855 |
| - | 327081 |
| + | 327482 |
| + | 327497 |
| + | 329242 |
| + | 329713 |
| + | 329715 |
| + | 334030 |
| - | 334062 |
| + | 334440 |
| + | 334530 |
| - | 336460 |
| + | 338202 |
| + | 338203 |
| + | 338204 |
| + | 338907 |
| + | 339195 |

| | |
|---|---|
| + | 339510 |
| - | 339765 |
| + | 339894 |
| + | 339895 |
| - | 339955 |
| + | 339990 |
| + | 339992 |
| - | 341409 |
| + | 341444 |
| + | 342499 |
| + | 344511 |
| + | 346486 |
| + | 346489 |
| + | 346595 |
| + | 346596 |
| + | 346597 |
| + | 346599 |
| + | 348161 |
| - | 348444 |
| + | 348857 |
| - | 349522 |
| + | 351325 |
| - | 351800 |
| + | 352803 |
| + | 352804 |
| - | 355645 |
| - | 355646 |
| + | 355713 |
| - | 358212 |
| - | 358213 |
| - | 358214 |
| + | 360815 |
| + | 361471 |
| - | 362797 |
| + | 362897 |
| + | 364223 |
| + | 365802 |
| - | 366867 |
| - | 368042 |
| - | 368883 |
| - | 368884 |
| + | 369194 |
| + | 369196 |
| + | 369299 |
| - | 369873 |
| + | 370231 |
| + | 371104 |
| + | 372088 |
| - | 372620 |
| + | 372822 |

| | |
|---|---|
| + | 373376 |
| - | 373742 |
| - | 374519 |
| + | 374762 |
| - | 375824 |
| - | 375827 |
| - | 375828 |
| + | 375837 |
| + | 375956 |
| + | 375958 |
| + | 376678 |
| + | 377081 |
| - | 377087 |
| - | 384869 |
| - | 391380 |
| - | 394402 |
| + | 403188 |
| + | 403189 |
| + | 405225 |
| - | 405491 |
| - | 406049 |
| + | 406087 |
| + | 406606 |
| + | 407717 |
| - | 408164 |
| - | 408917 |
| - | 408918 |
| - | 409544 |
| - | 411513 |
| - | 412191 |
| - | 412521 |
| + | 412542 |
| - | 415131 |
| + | 415328 |
| + | 415332 |
| - | 416568 |
| - | 417745 |
| - | 417748 |
| + | 417969 |
| + | 419733 |
| - | 420054 |
| + | 423209 |
| + | 423210 |
| - | 423543 |
| - | 426429 |
| + | 426540 |
| + | 427232 |
| + | 428359 |
| - | 428777 |
| + | 428799 |

| | |
|---|---|
| + | 429534 |
| + | 429878 |
| - | 430388 |
| - | 430395 |
| - | 430487 |
| + | 431053 |
| + | 432215 |
| + | 432277 |
| - | 432397 |
| + | 433220 |
| + | 433567 |
| - | 433824 |
| - | 435249 |
| - | 438420 |
| + | 439562 |
| - | 439615 |
| + | 439704 |
| + | 439709 |
| - | 441508 |
| + | 442923 |
| - | 449057 |
| + | 449166 |
| + | 449171 |
| + | 449172 |
| + | 449648 |
| + | 449649 |
| + | 449650 |
| + | 449651 |
| + | 449652 |
| - | 451217 |
| - | 452416 |
| - | 452783 |
| + | 452804 |
| + | 452806 |
| + | 453997 |
| + | 453998 |
| - | 454105 |
| - | 455283 |
| - | 455284 |
| - | 455285 |
| - | 455948 |
| + | 456025 |
| - | 456352 |
| + | 456926 |
| - | 459902 |
| + | 460056 |
| - | 460549 |
| + | 462183 |
| + | 463184 |
| - | 463277 |

| | |
|---|---|
| + | 463289 |
| - | 463474 |
| - | 463807 |
| - | 463808 |
| + | 463914 |
| + | 463916 |
| + | 465998 |
| - | 466327 |
| + | 467053 |
| + | 468521 |
| + | 469367 |
| + | 469368 |
| - | 471009 |
| - | 471211 |
| + | 471680 |
| + | 472575 |
| + | 472663 |
| - | 473245 |
| + | 473776 |
| + | 474264 |
| + | 474266 |
| + | 474706 |
| - | 475902 |
| + | 475919 |
| + | 476033 |
| + | 476524 |
| + | 476527 |
| + | 478860 |
| + | 478901 |
| - | 482828 |
| + | 484526 |
| + | 486089 |
| + | 486090 |
| + | 486091 |
| + | 488811 |
| + | 488812 |
| - | 491066 |
| - | 491068 |
| - | 492489 |
| - | 492493 |
| - | 492937 |
| - | 492938 |
| - | 492939 |
| + | 493530 |
| + | 494479 |
| + | 494480 |
| - | 496586 |
| - | 496587 |
| + | 496995 |
| - | 497713 |

| | |
|---|---|
| + | 498608 |
| + | 500124 |
| + | 500852 |
| + | 501530 |
| - | 502958 |
| + | 504661 |
| + | 504670 |
| + | 505760 |
| - | 506050 |
| - | 506666 |
| - | 506670 |
| + | 506846 |
| - | 507125 |
| + | 507135 |
| + | 507138 |
| + | 507762 |
| - | 508091 |
| - | 508096 |
| + | 508119 |
| + | 508197 |
| + | 508200 |
| + | 509356 |
| + | 509746 |
| - | 509806 |
| + | 510878 |
| + | 511030 |
| - | 511779 |
| + | 512755 |
| + | 512788 |
| + | 515111 |
| + | 515112 |
| - | 515459 |
| + | 515511 |
| - | 515641 |
| - | 516416 |
| + | 517339 |
| + | 517340 |
| + | 518505 |
| + | 518604 |
| + | 518607 |
| + | 518608 |
| + | 519357 |
| + | 519362 |
| + | 519365 |
| - | 519545 |
| - | 520153 |
| - | 520343 |
| + | 522055 |
| + | 522056 |
| - | 523680 |

| | |
|---|---|
| - | 524095 |
| + | 524331 |
| + | 524677 |
| + | 524787 |
| + | 524788 |
| + | 524789 |
| + | 525708 |
| + | 525711 |
| + | 525714 |
| + | 526011 |
| + | 527371 |
| + | 527907 |
| + | 528621 |
| + | 528624 |
| + | 528783 |
| - | 531328 |
| - | 531550 |
| + | 531750 |
| - | 532639 |
| - | 532929 |
| + | 533248 |
| - | 536158 |
| + | 536347 |
| - | 539675 |
| - | 545063 |
| + | 545572 |
| + | 545573 |
| - | 546414 |
| - | 547011 |
| - | 547013 |
| + | 547261 |
| + | 547773 |
| + | 548016 |
| + | 548197 |
| + | 548392 |
| + | 548701 |
| + | 549976 |
| - | 550158 |
| - | 550517 |
| + | 551075 |
| - | 552349 |
| - | 552532 |
| + | 552590 |
| - | 553524 |
| + | 553664 |
| + | 554209 |
| - | 554337 |
| + | 554874 |
| + | 555019 |
| + | 556415 |

| | |
|---|---|
| + | 557046 |
| + | 557209 |
| + | 557826 |
| + | 558222 |
| + | 559149 |
| + | 559152 |
| - | 559608 |
| - | 559609 |
| - | 560552 |
| - | 560638 |
| + | 560646 |
| + | 561516 |
| - | 561884 |
| + | 562469 |
| + | 562472 |
| - | 562888 |
| + | 563203 |
| + | 563371 |
| - | 564571 |
| - | 564573 |
| + | 564669 |
| - | 565618 |
| + | 565724 |
| + | 566163 |
| - | 566301 |
| + | 567769 |
| + | 568226 |
| - | 568483 |
| + | 568504 |
| - | 570000 |
| - | 571303 |
| + | 572940 |
| + | 573352 |
| - | 574050 |
| + | 574287 |
| - | 574458 |
| + | 574655 |
| + | 574659 |
| + | 575521 |
| - | 575899 |
| + | 575951 |
| - | 576011 |
| + | 576141 |
| - | 576154 |
| + | 576189 |
| - | 576404 |
| - | 577175 |
| - | 578223 |
| + | 578304 |
| + | 578306 |

| | |
|---|---|
| + | 578654 |
| + | 579337 |
| + | 580108 |
| + | 581676 |
| + | 583228 |
| + | 584885 |
| - | 585614 |
| - | 585804 |
| + | 585847 |
| + | 586213 |
| - | 587096 |
| - | 587115 |
| + | 587462 |
| - | 587508 |
| + | 587659 |
| + | 588915 |
| - | 589631 |
| + | 589681 |
| + | 591946 |
| - | 594123 |
| - | 595029 |
| + | 595030 |
| - | 595031 |
| - | 596080 |
| + | 596321 |
| + | 598140 |
| + | 598676 |
| + | 600571 |
| + | 601267 |
| - | 602007 |
| + | 602970 |
| + | 606371 |
| + | 606565 |
| - | 606605 |
| + | 607238 |
| + | 607319 |
| - | 608790 |
| - | 608792 |
| + | 608884 |
| - | 609320 |
| + | 611227 |
| - | 612850 |
| - | 613325 |
| + | 613563 |
| + | 615282 |
| - | 615751 |
| + | 615831 |
| + | 618042 |
| + | 618045 |
| - | 618184 |

| | |
|---|---|
| - | 618187 |
| + | 618652 |
| - | 619752 |
| - | 621522 |
| + | 621559 |
| + | 621675 |
| + | 621728 |
| - | 622566 |
| + | 622999 |
| - | 623320 |
| + | 624413 |
| + | 624414 |
| - | 626457 |
| + | 626591 |
| - | 628802 |
| + | 629078 |
| + | 630128 |
| + | 630129 |
| + | 630665 |
| + | 632173 |
| + | 634103 |
| + | 634823 |
| + | 634842 |
| + | 634929 |
| + | 634932 |
| + | 635098 |
| + | 636688 |
| + | 637054 |
| + | 637157 |
| + | 640117 |
| + | 640138 |
| + | 640268 |
| + | 640617 |
| + | 640619 |
| - | 641622 |
| - | 641624 |
| + | 641633 |
| + | 641634 |
| + | 643224 |
| + | 644881 |
| + | 646322 |
| + | 646412 |
| - | 646478 |
| + | 646484 |
| + | 646567 |
| + | 646883 |
| + | 647469 |
| - | 649703 |
| + | 649727 |
| + | 649838 |

| | |
|---|---|
| - | 652260 |
| + | 653988 |
| + | 655121 |
| + | 655123 |
| + | 655298 |
| + | 655816 |
| + | 656675 |
| + | 656889 |
| + | 658501 |
| - | 659336 |
| - | 659547 |
| + | 659566 |
| + | 659571 |
| + | 660391 |
| - | 660992 |
| - | 662625 |
| + | 663347 |
| + | 663440 |
| + | 663610 |
| - | 663732 |
| + | 664164 |
| - | 664897 |
| + | 667195 |
| - | 667282 |
| + | 667402 |
| - | 668745 |
| - | 668746 |
| - | 668990 |
| + | 669421 |
| + | 670055 |
| + | 671198 |
| + | 671209 |
| - | 672369 |
| + | 674519 |
| - | 675895 |
| - | 678512 |
| + | 679069 |
| - | 679219 |
| - | 679240 |
| + | 679365 |
| + | 679513 |
| - | 680555 |
| + | 680673 |
| - | 681082 |
| - | 681500 |
| - | 681502 |
| - | 683392 |
| - | 683395 |
| + | 685051 |
| - | 686608 |

| | |
|---|---|
| + | 686933 |
| + | 688145 |
| + | 688861 |
| - | 689657 |
| + | 690829 |
| + | 691684 |
| + | 692607 |
| + | 692609 |
| + | 692707 |
| + | 694425 |
| - | 695417 |
| + | 696069 |
| + | 696134 |
| + | 697100 |
| + | 697101 |
| + | 697411 |
| + | 697996 |
| + | 698372 |
| + | 699274 |
| - | 700539 |
| - | 700592 |
| - | 701618 |
| + | 710618 |
| + | 711115 |
| - | 711947 |
| - | 711948 |
| - | 711949 |
| + | 712477 |
| - | 713627 |
| + | 713634 |
| + | 713636 |
| + | 713639 |
| + | 715414 |
| + | 715415 |
| + | 715416 |
| - | 715921 |
| - | 715923 |
| + | 716218 |
| + | 716819 |
| + | 718589 |
| + | 718591 |
| + | 719342 |
| - | 720269 |
| - | 722272 |
| + | 722284 |
| + | 724878 |
| + | 724879 |
| - | 726769 |
| - | 726778 |
| - | 728444 |

| | |
|---|---|
| + | 728617 |
| + | 728618 |
| + | 732657 |
| + | 732659 |
| - | 732895 |
| - | 732897 |
| + | 732914 |
| - | 733836 |
| + | 735328 |
| + | 736155 |
| + | 736385 |
| + | 736386 |
| + | 738623 |
| + | 739664 |
| - | 740096 |
| + | 740960 |
| + | 741203 |
| - | 741251 |
| + | 742321 |
| - | 742954 |
| - | 743180 |
| - | 743617 |
| - | 744422 |
| + | 744827 |
| + | 745223 |
| + | 746215 |
| - | 747922 |
| - | 747924 |
| - | 749592 |
| + | 750932 |
| + | 751449 |
| + | 752359 |
| + | 752361 |
| + | 752363 |
| - | 753227 |
| + | 753800 |
| + | 753801 |
| + | 754141 |
| + | 755077 |
| + | 757073 |
| + | 757075 |
| + | 758056 |
| + | 759544 |
| + | 777497 |
| - | 778160 |
| - | 780858 |
| + | 782256 |
| - | 783394 |
| - | 784345 |
| + | 784363 |

| | |
|---|---|
| - | 785467 |
| - | 785469 |
| - | 785471 |
| - | 785930 |
| + | 786653 |
| - | 786707 |
| + | 787668 |
| - | 787810 |
| + | 787955 |
| + | 787958 |
| - | 790182 |
| + | 790270 |
| + | 790747 |
| + | 791302 |
| - | 792565 |
| + | 795214 |
| + | 796024 |
| + | 796025 |
| + | 796130 |
| - | 796691 |
| + | 801927 |
| + | 803863 |
| - | 804579 |
| - | 804581 |
| - | 805389 |
| - | 806897 |
| - | 806899 |
| - | 806902 |
| + | 807045 |
| + | 809528 |
| - | 810714 |
| + | 811521 |
| + | 811523 |
| + | 811525 |
| + | 812112 |
| + | 812116 |
| - | 813905 |
| - | 815991 |
| - | 815992 |
| + | 816085 |
| + | 816087 |
| - | 817917 |
| - | 818852 |
| - | 820024 |
| - | 820572 |
| + | 821002 |
| + | 821410 |
| + | 822713 |
| - | 823839 |
| - | 824304 |

| | |
|---|---|
| - | 824307 |
| - | 826777 |
| - | 826783 |
| - | 827315 |
| + | 827425 |
| + | 827427 |
| + | 827941 |
| - | 828745 |
| - | 828746 |
| - | 829462 |
| - | 830310 |
| + | 830926 |
| + | 834313 |
| + | 834613 |
| + | 836625 |
| + | 836626 |
| - | 838956 |
| - | 839259 |
| - | 839264 |
| + | 839269 |
| - | 839676 |
| - | 840515 |
| + | 840624 |
| + | 840627 |
| - | 842881 |
| + | 843141 |
| + | 843890 |
| - | 844050 |
| + | 844229 |
| + | 844231 |
| + | 844745 |
| - | 845866 |
| + | 846238 |
| + | 846446 |
| + | 846594 |
| + | 846980 |
| + | 847211 |
| + | 848293 |
| + | 849882 |
| + | 850319 |
| + | 850321 |
| - | 852158 |
| + | 853544 |
| + | 854383 |
| + | 856240 |
| - | 859535 |
| + | 859624 |
| + | 859713 |
| + | 859715 |
| + | 859863 |

| | |
|---|---|
| - | 860633 |
| - | 861833 |
| - | 861835 |
| - | 861839 |
| + | 861953 |
| + | 862515 |
| + | 864511 |
| - | 865064 |
| + | 866342 |
| - | 867663 |
| - | 868039 |
| - | 869172 |
| + | 869248 |
| + | 869249 |
| + | 869250 |
| + | 869251 |
| + | 870022 |
| - | 871272 |
| - | 871273 |
| - | 872318 |
| - | 872346 |
| - | 873279 |
| + | 873335 |
| + | 873336 |
| + | 875146 |
| + | 875626 |
| - | 875646 |
| + | 876184 |
| + | 877561 |
| + | 877572 |
| - | 882229 |
| + | 883721 |
| + | 885637 |
| - | 886196 |
| - | 886201 |
| - | 886412 |
| - | 887703 |
| - | 888108 |
| - | 889734 |
| + | 889996 |
| + | 890151 |
| + | 890855 |
| + | 891513 |
| - | 891581 |
| + | 893864 |
| + | 893866 |
| - | 894231 |
| - | 894476 |
| + | 897551 |
| + | 897552 |

| | |
|---|---|
| - | 898643 |
| + | 898937 |
| + | 899980 |
| - | 901515 |
| + | 903574 |
| + | 903785 |
| + | 903786 |
| + | 905018 |
| - | 909817 |
| - | 909818 |
| + | 909987 |
| - | 910722 |
| + | 910802 |
| + | 911728 |
| - | 912179 |
| - | 912609 |
| + | 913894 |
| + | 913897 |
| + | 915852 |
| - | 916039 |
| - | 916633 |
| - | 916635 |
| - | 916636 |
| + | 916729 |
| + | 916730 |
| - | 917761 |
| - | 920374 |
| + | 920394 |
| + | 920396 |
| + | 921391 |
| + | 922589 |
| + | 923547 |
| + | 923548 |
| - | 924397 |
| - | 924603 |
| + | 924615 |
| + | 925467 |
| - | 925569 |
| - | 925580 |
| + | 926853 |
| - | 926908 |
| - | 927523 |
| + | 928331 |
| + | 928773 |
| + | 929389 |
| + | 929390 |
| + | 930737 |
| + | 930739 |
| + | 933054 |
| + | 935582 |

| | |
|---|---|
| - | 935588 |
| - | 935590 |
| + | 936638 |
| + | 936639 |
| + | 936906 |
| + | 936907 |
| + | 937001 |
| + | 938229 |
| + | 938516 |
| + | 938658 |
| + | 938660 |
| + | 938661 |
| + | 939088 |
| - | 943768 |
| - | 945326 |
| - | 945328 |
| + | 945427 |
| + | 945428 |
| + | 946518 |
| + | 946521 |
| + | 947951 |
| + | 948317 |
| + | 948420 |
| + | 951442 |
| + | 951444 |
| + | 952397 |
| + | 952580 |
| + | 953185 |
| + | 953996 |
| + | 953997 |
| - | 954178 |
| + | 954269 |
| + | 954561 |
| + | 954860 |
| + | 954861 |
| - | 955621 |
| + | 955647 |
| + | 955649 |
| + | 956731 |
| + | 956923 |
| + | 957122 |
| + | 959217 |
| - | 961008 |
| - | 961010 |
| + | 961273 |
| + | 965197 |
| + | 965874 |
| + | 965875 |
| - | 966270 |
| - | 966902 |

| | |
|---|---|
| + | 967031 |
| + | 967202 |
| + | 967651 |
| - | 967878 |
| - | 967880 |
| + | 968931 |
| + | 970116 |
| + | 970118 |
| + | 970724 |
| - | 973005 |
| - | 973006 |
| - | 973119 |
| + | 973647 |
| + | 975398 |
| + | 975471 |
| - | 976205 |
| + | 976533 |
| + | 976535 |
| + | 977731 |
| - | 978476 |
| + | 980440 |
| - | 981024 |
| + | 984030 |
| + | 984035 |
| - | 984584 |
| - | 984588 |
| + | 984797 |
| + | 984798 |
| + | 985399 |
| + | 985847 |
| - | 985860 |
| - | 986483 |
| + | 986956 |
| + | 986958 |
| - | 988364 |
| - | 988891 |
| + | 989253 |
| + | 991372 |
| - | 991454 |
| - | 991776 |
| + | 993513 |
| + | 996626 |
| + | 996627 |
| - | 997085 |
| + | 997204 |
| + | 997683 |
| + | 997685 |
| + | 999100 |
| + | 1000286 |
| + | 1001206 |

| | |
|---|---|
| + | 1001724 |
| + | 1001848 |
| + | 1002353 |
| + | 1002354 |
| + | 1003265 |
| + | 1004181 |
| + | 1004860 |
| + | 1006748 |
| + | 1008640 |
| + | 1008642 |
| + | 1008746 |
| - | 1010175 |
| + | 1012488 |
| - | 1013302 |
| - | 1013313 |
| - | 1013879 |
| - | 1013880 |
| - | 1013881 |
| - | 1013882 |
| - | 1013883 |
| + | 1016366 |
| + | 1017861 |
| - | 1018150 |
| + | 1018660 |
| - | 1018694 |
| + | 1018934 |
| + | 1018937 |
| - | 1020046 |
| + | 1020870 |
| - | 1021020 |
| - | 1021022 |
| + | 1021032 |
| + | 1022203 |
| - | 1022443 |
| - | 1022453 |
| + | 1022608 |
| + | 1023222 |
| + | 1024845 |
| - | 1027737 |
| + | 1027749 |
| - | 1029591 |
| - | 1030085 |
| - | 1030162 |
| - | 1030163 |
| + | 1030242 |
| - | 1030271 |
| - | 1030860 |
| + | 1031355 |
| + | 1031359 |
| + | 1033460 |

| | |
|---|---|
| - | 1033903 |
| + | 1034008 |
| + | 1034010 |
| - | 1034859 |
| + | 1035405 |
| + | 1035407 |
| - | 1037627 |
| - | 1037863 |
| - | 1038448 |
| - | 1038449 |
| - | 1038450 |
| + | 1038633 |
| + | 1038635 |
| - | 1038857 |
| + | 1038872 |
| + | 1039491 |
| - | 1041799 |
| + | 1041939 |
| + | 1041942 |
| + | 1041945 |
| + | 1042453 |
| + | 1042937 |
| - | 1044898 |
| - | 1044899 |
| + | 1044986 |
| + | 1048869 |
| - | 1049776 |
| + | 1049831 |
| + | 1050271 |
| - | 1050704 |
| + | 1051479 |
| - | 1052578 |
| + | 1053464 |
| + | 1054708 |
| + | 1054709 |
| + | 1055635 |
| + | 1056147 |
| - | 1056240 |
| - | 1056243 |
| + | 1056390 |
| + | 1057649 |
| - | 1058656 |
| + | 1058678 |
| - | 1058895 |
| - | 1059883 |
| + | 1060409 |
| - | 1061340 |
| - | 1061343 |
| + | 1061453 |
| - | 1062501 |

| | |
|---|---|
| - | 1062502 |
| - | 1062504 |
| + | 1062574 |
| + | 1062575 |
| + | 1064232 |
| + | 1064823 |
| - | 1065517 |
| - | 1068087 |
| + | 1069013 |
| + | 1070147 |
| + | 1070600 |
| - | 1071282 |
| - | 1071313 |
| - | 1071967 |
| - | 1072595 |
| - | 1072597 |
| + | 1072740 |
| + | 1072741 |
| + | 1072742 |
| + | 1072743 |
| - | 1074337 |
| - | 1075246 |
| - | 1076409 |
| - | 1076412 |
| - | 1076987 |
| - | 1076989 |
| - | 1081369 |
| - | 1081371 |
| - | 1081372 |
| + | 1082766 |
| - | 1083743 |
| + | 1084535 |
| + | 1085345 |
| + | 1086068 |
| + | 1086069 |
| + | 1087204 |
| + | 1087223 |
| + | 1087495 |
| + | 1087703 |
| + | 1088061 |
| + | 1089683 |
| + | 1089707 |
| + | 1090093 |
| + | 1090924 |
| - | 1091254 |
| + | 1092287 |
| - | 1093513 |
| - | 1093763 |
| + | 1095039 |
| - | 1097916 |

| | |
|---|---|
| - | 1097917 |
| - | 1097952 |
| - | 1098146 |
| - | 1098281 |
| - | 1098284 |
| + | 1098378 |
| + | 1098379 |
| + | 1099539 |
| + | 1100206 |
| + | 1100922 |
| + | 1100924 |
| + | 1100927 |
| + | 1102131 |
| - | 1102499 |
| + | 1103076 |
| + | 1105984 |
| + | 1105985 |
| + | 1106507 |
| + | 1107061 |
| + | 1107688 |
| - | 1109343 |
| + | 1109656 |
| - | 1111938 |
| - | 1111940 |
| - | 1112514 |
| + | 1112854 |
| - | 1116841 |
| - | 1116842 |
| - | 1116843 |
| - | 1116844 |
| + | 1117073 |
| + | 1117075 |
| - | 1118662 |
| + | 1118802 |
| + | 1118804 |
| + | 1121272 |
| - | 1121432 |
| + | 1121526 |
| + | 1122372 |
| + | 1122766 |
| + | 1123491 |
| + | 1124403 |
| + | 1124404 |
| + | 1125907 |
| - | 1126040 |
| - | 1126887 |
| + | 1129279 |
| + | 1129282 |
| + | 1129811 |
| + | 1129813 |

| | |
|---|---|
| + | 1130148 |
| - | 1132128 |
| - | 1132130 |
| - | 1132132 |
| + | 1133422 |
| + | 1134850 |
| + | 1136292 |
| + | 1136294 |
| + | 1144400 |
| + | 1149113 |
| + | 1150478 |
| + | 1150479 |
| + | 1150838 |
| - | 1151066 |
| - | 1151077 |
| + | 1152208 |
| + | 1152209 |
| + | 1153235 |
| + | 1153236 |
| + | 1153237 |
| + | 1153760 |
| + | 1153762 |
| + | 1155941 |
| - | 1156689 |
| - | 1159332 |
| - | 1159715 |
| + | 1159728 |
| + | 1159918 |
| + | 1160170 |
| + | 1162052 |
| + | 1162213 |
| + | 1163017 |
| - | 1164181 |
| + | 1164250 |
| + | 1165499 |
| - | 1165975 |
| - | 1165976 |
| - | 1168960 |
| - | 1168962 |
| - | 1168963 |
| - | 1175067 |
| - | 1180860 |
| - | 1180861 |
| - | 1180862 |
| - | 1181428 |
| + | 1181451 |
| + | 1181453 |
| - | 1182567 |
| + | 1183132 |
| + | 1184017 |

| | |
|---|---|
| + | 1184657 |
| - | 1184976 |
| + | 1185070 |
| + | 1187527 |
| + | 1188135 |
| - | 1188594 |
| + | 1188662 |
| + | 1188665 |
| - | 1189287 |
| + | 1190105 |
| - | 1191342 |
| - | 1191373 |
| + | 1191394 |
| + | 1191397 |
| + | 1192192 |
| + | 1192231 |
| + | 1192833 |
| + | 1194161 |
| + | 1194999 |
| - | 1195767 |
| - | 1200224 |
| - | 1202391 |
| + | 1202868 |
| - | 1203116 |
| + | 1203925 |
| + | 1203926 |
| - | 1204946 |
| - | 1204947 |
| - | 1204948 |
| + | 1205053 |
| + | 1206200 |
| + | 1206574 |
| + | 1206580 |
| - | 1206581 |
| + | 1206582 |
| + | 1207283 |
| + | 1207929 |
| - | 1208097 |
| - | 1208100 |
| + | 1208155 |
| + | 1208157 |
| + | 1208334 |
| + | 1208825 |
| + | 1211373 |
| + | 1211375 |
| + | 1215356 |
| + | 1219218 |
| - | 1219388 |
| - | 1219399 |
| - | 1219401 |

| | |
|---|---|
| - | 1219402 |
| + | 1219702 |
| - | 1221640 |
| + | 1225515 |
| - | 1226192 |
| + | 1226863 |
| + | 1227616 |
| + | 1227617 |
| - | 1228089 |
| + | 1228935 |
| + | 1228940 |
| - | 1229634 |
| + | 1229863 |
| + | 1231194 |
| + | 1231197 |
| - | 1232846 |
| + | 1233222 |
| + | 1233427 |
| + | 1233429 |
| - | 1234479 |
| - | 1234991 |
| - | 1235081 |
| - | 1235083 |
| - | 1236065 |
| - | 1236520 |
| - | 1236521 |
| + | 1236582 |
| - | 1239731 |
| - | 1240191 |
| + | 1240281 |
| - | 1240685 |
| + | 1242238 |
| - | 1244836 |
| + | 1245569 |
| + | 1246843 |
| + | 1247154 |
| + | 1247736 |
| - | 1248368 |
| + | 1250221 |
| - | 1250593 |
| + | 1250861 |
| + | 1252122 |
| + | 1253640 |
| + | 1255954 |
| + | 1255955 |
| - | 1256030 |
| + | 1256092 |
| + | 1257413 |
| - | 1258180 |
| - | 1258183 |

| | |
|---|---|
| + | 1258281 |
| + | 1258282 |
| - | 1261388 |
| - | 1261390 |
| - | 1262645 |
| + | 1262746 |
| + | 1262747 |
| + | 1263167 |
| - | 1263290 |
| + | 1263342 |
| - | 1263872 |
| - | 1265004 |
| + | 1265006 |
| - | 1265006 |
| + | 1265009 |
| + | 1266193 |
| + | 1266357 |
| + | 1266598 |
| + | 1266959 |
| + | 1267018 |
| + | 1268282 |
| + | 1268283 |
| + | 1268376 |
| + | 1268764 |
| + | 1268866 |
| - | 1269047 |
| - | 1270113 |
| + | 1270560 |
| + | 1272686 |
| + | 1272688 |
| + | 1272689 |
| + | 1275749 |
| + | 1275750 |
| - | 1276869 |
| - | 1276870 |
| + | 1277634 |
| + | 1277637 |
| + | 1277881 |
| - | 1278414 |
| - | 1278415 |
| - | 1278932 |
| + | 1279033 |
| + | 1279036 |
| + | 1280408 |
| - | 1280506 |
| - | 1281004 |
| + | 1281024 |
| + | 1281026 |
| - | 1283351 |
| - | 1283353 |

| | |
|---|---|
| - | 1284800 |
| - | 1284802 |
| + | 1285555 |
| + | 1289272 |
| + | 1289273 |
| + | 1289547 |
| - | 1290579 |
| - | 1290581 |
| - | 1290976 |
| - | 1290977 |
| + | 1291186 |
| + | 1291187 |
| - | 1291807 |
| - | 1293991 |
| - | 1293994 |
| + | 1294085 |
| + | 1294088 |
| + | 1296315 |
| - | 1296540 |
| - | 1296542 |
| - | 1297623 |
| + | 1297624 |
| + | 1298353 |
| + | 1298354 |
| - | 1300143 |
| + | 1300396 |
| - | 1302460 |
| - | 1306194 |
| + | 1308753 |
| + | 1308764 |
| - | 1312276 |
| + | 1314337 |
| - | 1314338 |
| + | 1314424 |
| + | 1315836 |
| + | 1315838 |
| - | 1317430 |
| - | 1317431 |
| + | 1317510 |
| - | 1318927 |
| + | 1318987 |
| - | 1319689 |
| + | 1319879 |
| + | 1320676 |
| - | 1321703 |
| - | 1321704 |
| + | 1321805 |
| + | 1324862 |
| + | 1325011 |
| + | 1325866 |

| | |
|---|---|
| + | 1330774 |
| - | 1335373 |
| + | 1336601 |
| - | 1339091 |
| - | 1345298 |
| + | 1347282 |
| - | 1348369 |
| - | 1348953 |
| - | 1348955 |
| - | 1348957 |
| - | 1349195 |
| + | 1350056 |
| - | 1350452 |
| - | 1351137 |
| - | 1351138 |
| - | 1351283 |
| - | 1352744 |
| + | 1353011 |
| - | 1357833 |
| - | 1357834 |
| - | 1357837 |
| - | 1359339 |
| - | 1359344 |
| - | 1359422 |
| + | 1359433 |
| + | 1359719 |
| + | 1360365 |
| + | 1360366 |
| + | 1367576 |
| - | 1370951 |
| + | 1371316 |
| - | 1371903 |
| + | 1371978 |
| + | 1371985 |
| + | 1371987 |
| + | 1372753 |
| + | 1374847 |
| + | 1375244 |
| + | 1376299 |
| + | 1376327 |
| + | 1376581 |
| + | 1377206 |
| + | 1377208 |
| + | 1378223 |
| + | 1378224 |
| + | 1378225 |
| + | 1380951 |
| + | 1381776 |
| - | 1381905 |
| + | 1381994 |

| | |
|---|---|
| + | 1382432 |
| - | 1383171 |
| + | 1384239 |
| - | 1385928 |
| - | 1385930 |
| + | 1386013 |
| - | 1387029 |
| + | 1387188 |
| - | 1389108 |
| + | 1389118 |
| + | 1391680 |
| + | 1391681 |
| - | 1391871 |
| - | 1391872 |
| + | 1393723 |
| + | 1394751 |
| + | 1395605 |
| - | 1396051 |
| + | 1396205 |
| + | 1396772 |
| - | 1397202 |
| - | 1397355 |
| - | 1397769 |
| + | 1397909 |
| - | 1398149 |
| + | 1398153 |
| + | 1398424 |
| - | 1398955 |
| + | 1399380 |
| - | 1399491 |
| + | 1400643 |
| - | 1401540 |
| - | 1403115 |
| + | 1408932 |
| + | 1409334 |
| - | 1409774 |
| - | 1409775 |
| + | 1409886 |
| + | 1410607 |
| + | 1410609 |
| + | 1410859 |
| + | 1411775 |
| + | 1411776 |
| + | 1411867 |
| + | 1411870 |
| + | 1412600 |
| - | 1413060 |
| - | 1414901 |
| + | 1414957 |
| + | 1414959 |

| | |
|---|---|
| + | 1416036 |
| + | 1419184 |
| + | 1421321 |
| + | 1422275 |
| - | 1424721 |
| - | 1424722 |
| + | 1424800 |
| - | 1425257 |
| - | 1425259 |
| + | 1425276 |
| + | 1425603 |
| + | 1425605 |
| + | 1425607 |
| + | 1426847 |
| + | 1426852 |
| + | 1426854 |
| + | 1426857 |
| - | 1429904 |
| + | 1430543 |
| - | 1430554 |
| - | 1430555 |
| + | 1430622 |
| + | 1430635 |
| + | 1432020 |
| - | 1433016 |
| + | 1433170 |
| + | 1434665 |
| - | 1435279 |
| - | 1435281 |
| - | 1437808 |
| - | 1437927 |
| - | 1437989 |
| + | 1438836 |
| + | 1439242 |
| - | 1440297 |
| + | 1441281 |
| + | 1441359 |
| + | 1441647 |
| + | 1441737 |
| - | 1442620 |
| + | 1443327 |
| + | 1443626 |
| + | 1445273 |
| + | 1445547 |
| + | 1446577 |
| + | 1446806 |
| + | 1447226 |
| + | 1447230 |
| - | 1447884 |
| - | 1447885 |

| | |
|---|---|
| + | 1447982 |
| + | 1447983 |
| + | 1448335 |
| + | 1448337 |
| + | 1448916 |
| - | 1449826 |
| + | 1450428 |
| + | 1450671 |
| + | 1450676 |
| + | 1451264 |
| - | 1451522 |
| + | 1451862 |
| - | 1453554 |
| - | 1453555 |
| + | 1453653 |
| + | 1453654 |
| + | 1454278 |
| - | 1455002 |
| + | 1455082 |
| + | 1456087 |
| + | 1456766 |
| + | 1456850 |
| + | 1456851 |
| + | 1456852 |
| + | 1457079 |
| + | 1458293 |
| - | 1458325 |
| + | 1459340 |
| + | 1459343 |
| - | 1463507 |
| + | 1463614 |
| + | 1464205 |
| + | 1464206 |
| + | 1465032 |
| + | 1465674 |
| + | 1465675 |
| - | 1467445 |
| + | 1467704 |
| - | 1469044 |
| + | 1469971 |
| + | 1470530 |
| + | 1473070 |
| - | 1473073 |
| - | 1473192 |
| - | 1473427 |
| - | 1473428 |
| - | 1473429 |
| + | 1473573 |
| + | 1474095 |
| - | 1475058 |

| | |
|---|---|
| - | 1475060 |
| - | 1476469 |
| + | 1476855 |
| + | 1477234 |
| - | 1477952 |
| - | 1477954 |
| + | 1478038 |
| + | 1478991 |
| + | 1479145 |
| + | 1479884 |
| + | 1481311 |
| + | 1482206 |
| + | 1482208 |
| + | 1483449 |
| + | 1483553 |
| + | 1483555 |
| + | 1483557 |
| + | 1484049 |
| + | 1484051 |
| + | 1485059 |
| + | 1485410 |
| + | 1485427 |
| + | 1486010 |
| + | 1486995 |
| + | 1486997 |
| + | 1488942 |
| + | 1488945 |
| - | 1490611 |
| - | 1490613 |
| + | 1490740 |
| + | 1490742 |
| - | 1492077 |
| + | 1492177 |
| + | 1493718 |
| + | 1494748 |
| + | 1495485 |
| + | 1495692 |
| + | 1496131 |
| + | 1499864 |
| - | 1502098 |
| + | 1502930 |
| + | 1503558 |
| - | 1505164 |
| + | 1507459 |
| + | 1511267 |
| + | 1512076 |
| + | 1512077 |
| - | 1512243 |
| - | 1512244 |
| + | 1512346 |

| | |
|---|---|
| + | 1516110 |
| - | 1516256 |
| - | 1516257 |
| - | 1517621 |
| + | 1517801 |
| + | 1517802 |
| + | 1517803 |
| + | 1518303 |
| + | 1518306 |
| + | 1518504 |
| - | 1519858 |
| + | 1520501 |
| + | 1520502 |
| - | 1521311 |
| + | 1521315 |
| + | 1521323 |
| + | 1521325 |
| - | 1523531 |
| - | 1525038 |
| - | 1525042 |
| - | 1525237 |
| + | 1525349 |
| + | 1526508 |
| - | 1526773 |
| - | 1526776 |
| + | 1527210 |
| - | 1527286 |
| + | 1527510 |
| + | 1528115 |
| - | 1528217 |
| + | 1530719 |
| + | 1530720 |
| - | 1533734 |
| + | 1534070 |
| - | 1534670 |
| - | 1535834 |
| + | 1535889 |
| + | 1535890 |
| - | 1536896 |
| + | 1536979 |
| + | 1537402 |
| - | 1539456 |
| - | 1539756 |
| - | 1541638 |
| - | 1541640 |
| + | 1541743 |
| + | 1541862 |
| + | 1543530 |
| + | 1543870 |
| + | 1544087 |

| | |
|---|---|
| + | 1545383 |
| + | 1545640 |
| - | 1546032 |
| - | 1546033 |
| - | 1546034 |
| + | 1546091 |
| + | 1546092 |
| + | 1547981 |
| - | 1548613 |
| - | 1548617 |
| - | 1548858 |
| - | 1549212 |
| + | 1550631 |
| - | 1551298 |
| - | 1551299 |
| - | 1551300 |
| - | 1551569 |
| + | 1551917 |
| + | 1552371 |
| + | 1552372 |
| + | 1552374 |
| + | 1552844 |
| + | 1554155 |
| - | 1555081 |
| - | 1556618 |
| - | 1558157 |
| + | 1558557 |
| - | 1558996 |
| + | 1559118 |
| + | 1559219 |
| + | 1559486 |
| - | 1559735 |
| + | 1560800 |
| - | 1562838 |
| + | 1563426 |
| + | 1564046 |
| + | 1564081 |
| - | 1564807 |
| + | 1564853 |
| - | 1565737 |
| - | 1565738 |
| - | 1565898 |
| + | 1565915 |
| + | 1565917 |
| + | 1566656 |
| + | 1567622 |
| + | 1568406 |
| + | 1568407 |
| - | 1568566 |
| + | 1569443 |

| | |
|---|---|
| + | 1569770 |
| + | 1571858 |
| + | 1571859 |
| - | 1572840 |
| + | 1575101 |
| + | 1575107 |
| - | 1575140 |
| + | 1575234 |
| + | 1575238 |
| + | 1576109 |
| - | 1577132 |
| - | 1577274 |
| + | 1577380 |
| - | 1577460 |
| + | 1578336 |
| + | 1578479 |
| - | 1579398 |
| + | 1580062 |
| + | 1580063 |
| + | 1580065 |
| + | 1580560 |
| + | 1581183 |
| - | 1582660 |
| - | 1585312 |
| + | 1585639 |
| + | 1585986 |
| + | 1586247 |
| + | 1586338 |
| + | 1586456 |
| + | 1590379 |
| + | 1592052 |
| + | 1592055 |
| + | 1593612 |
| + | 1594076 |
| - | 1594673 |
| + | 1595510 |
| + | 1595738 |
| - | 1596048 |
| + | 1596189 |
| + | 1596307 |
| + | 1596447 |
| + | 1597364 |
| + | 1606502 |
| + | 1606504 |
| + | 1607308 |
| - | 1608779 |
| + | 1609113 |
| + | 1609282 |
| + | 1609283 |
| + | 1609284 |

| | |
|---|---|
| - | 1609364 |
| + | 1609549 |
| + | 1610203 |
| + | 1610227 |
| + | 1611056 |
| + | 1612179 |
| + | 1612482 |
| + | 1612483 |
| + | 1613066 |
| + | 1616709 |
| + | 1616711 |
| + | 1618153 |
| + | 1630104 |
| + | 1630105 |
| - | 1631545 |
| + | 1631910 |
| - | 1634731 |
| + | 1634774 |
| + | 1634776 |
| + | 1636265 |
| - | 1637873 |
| - | 1639372 |
| + | 1639717 |
| + | 1639816 |
| + | 1640661 |
| + | 1641594 |
| - | 1644483 |
| - | 1644485 |
| + | 1646373 |
| - | 1646408 |
| + | 1649096 |
| - | 1651221 |
| - | 1653943 |
| - | 1655863 |
| - | 1655864 |
| - | 1655866 |
| + | 1655976 |
| + | 1656035 |
| + | 1658529 |
| + | 1660254 |
| - | 1661165 |
| + | 1661931 |
| + | 1664017 |
| + | 1665309 |
| + | 1665315 |
| + | 1665321 |
| - | 1665725 |
| + | 1665851 |
| + | 1666536 |
| - | 1671498 |

| | |
|---|---|
| - | 1671500 |
| - | 1671676 |
| + | 1671789 |
| + | 1671792 |
| + | 1673586 |
| + | 1673588 |
| + | 1674197 |
| + | 1675139 |
| + | 1675941 |
| + | 1675943 |
| + | 1675945 |
| + | 1676479 |
| + | 1676481 |
| + | 1677436 |
| - | 1679490 |
| - | 1679938 |
| + | 1680380 |
| + | 1680388 |
| + | 1681587 |
| + | 1683506 |
| + | 1683518 |
| + | 1684886 |
| + | 1684997 |
| + | 1685230 |
| + | 1685436 |
| + | 1685805 |
| + | 1686175 |
| + | 1687162 |
| + | 1687163 |
| + | 1687262 |
| + | 1687265 |
| + | 1689264 |
| + | 1689665 |
| + | 1691053 |
| + | 1691212 |
| + | 1691214 |
| + | 1692754 |
| + | 1697613 |
| - | 1697988 |
| + | 1702043 |
| - | 1714109 |
| + | 1714621 |
| + | 1716478 |
| + | 1717474 |
| + | 1717475 |
| + | 1717821 |
| + | 1717822 |
| + | 1719775 |
| + | 1719776 |
| + | 1720319 |

| | |
|---|---|
| + | 1721109 |
| + | 1722705 |
| + | 1722829 |
| + | 1723516 |
| + | 1723824 |
| + | 1727066 |
| + | 1729056 |
| + | 1730722 |
| + | 1730902 |
| + | 1731600 |
| + | 1732938 |
| + | 1733132 |
| + | 1735857 |
| + | 1736088 |
| + | 1736090 |
| + | 1737638 |
| + | 1738820 |
| + | 1738822 |
| + | 1738854 |
| + | 1739341 |
| + | 1740637 |
| + | 1741153 |
| + | 1741860 |
| - | 1744129 |
| - | 1744133 |
| - | 1744331 |
| + | 1745408 |
| + | 1745936 |
| + | 1748356 |
| + | 1749377 |
| + | 1750243 |
| + | 1752191 |
| + | 1752192 |
| + | 1752560 |
| + | 1753949 |
| + | 1754740 |
| + | 1756350 |
| + | 1756961 |
| + | 1756962 |
| + | 1760414 |
| + | 1760810 |
| + | 1760815 |
| + | 1761550 |
| - | 1762767 |
| + | 1764539 |
| + | 1764616 |
| + | 1765819 |
| + | 1765821 |
| + | 1767146 |
| + | 1769809 |

| | |
|---|---|
| + | 1769810 |
| + | 1769811 |
| + | 1769813 |
| + | 1770429 |
| + | 1770431 |
| + | 1770820 |
| - | 1772416 |
| - | 1772682 |
| + | 1772768 |
| + | 1774196 |
| + | 1775447 |
| + | 1775705 |
| - | 1775756 |
| + | 1777730 |
| - | 1779190 |
| + | 1780404 |
| - | 1781155 |
| - | 1781821 |
| - | 1781822 |
| + | 1781906 |
| + | 1782645 |
| + | 1783175 |
| + | 1783674 |
| + | 1785375 |
| - | 1786017 |
| - | 1794720 |
| - | 1802770 |
| + | 1833660 |
| - | 1834800 |
| - | 1859874 |
| - | 1860393 |
| - | 1860859 |
| - | 1861299 |
| - | 1861300 |
| - | 1862007 |
| + | 1862905 |
| + | 1862907 |
| + | 1863386 |
| + | 1864202 |
| + | 1864203 |
| - | 1866025 |
| + | 1866361 |
| + | 1866363 |
| - | 1867135 |
| + | 1867347 |
| + | 1867349 |
| + | 1868116 |
| + | 1868404 |
| + | 1868569 |
| - | 1874010 |

| | |
|---|---|
| - | 1875257 |
| + | 1875268 |
| + | 1875271 |
| + | 1877934 |
| - | 1878651 |
| + | 1880053 |
| + | 1880455 |
| + | 1881409 |
| - | 1882150 |
| + | 1882439 |
| + | 1883140 |
| + | 1884162 |
| + | 1890832 |
| - | 1890946 |
| - | 1891036 |
| - | 1891727 |
| + | 1893350 |
| - | 1896054 |
| + | 1896238 |
| + | 1899555 |
| + | 1901864 |
| + | 1902169 |
| - | 1902242 |
| - | 1904258 |
| - | 1905027 |
| - | 1905592 |
| - | 1905593 |
| - | 1905594 |
| - | 1905595 |
| - | 1906738 |
| + | 1906759 |
| + | 1906984 |
| + | 1910211 |
| + | 1911502 |
| + | 1914051 |
| - | 1915272 |
| + | 1916636 |
| + | 1917421 |
| + | 1917422 |
| + | 1917505 |
| - | 1918284 |
| - | 1918285 |
| + | 1918372 |
| + | 1918531 |
| + | 1918728 |
| + | 1919433 |
| + | 1919435 |
| + | 1919834 |
| - | 1920066 |
| + | 1921983 |

| | |
|---|---|
| - | 1922253 |
| + | 1922263 |
| + | 1922503 |
| + | 1922504 |
| + | 1922505 |
| - | 1923054 |
| - | 1923117 |
| - | 1923118 |
| + | 1923994 |
| + | 1923997 |
| + | 1924447 |
| - | 1925233 |
| - | 1925441 |
| + | 1925548 |
| - | 1926459 |
| + | 1926608 |
| + | 1926610 |
| + | 1926612 |
| + | 1930055 |
| + | 1930540 |
| + | 1930752 |
| - | 1931867 |
| - | 1931869 |
| + | 1931882 |
| - | 1931921 |
| + | 1932084 |
| - | 1932529 |
| - | 1932530 |
| - | 1932532 |
| - | 1932534 |
| + | 1932648 |
| + | 1932651 |
| + | 1933176 |
| + | 1933177 |
| + | 1933261 |
| - | 1938456 |
| - | 1938521 |
| + | 1938859 |
| + | 1938860 |
| - | 1940320 |
| - | 1943688 |
| + | 1946239 |
| + | 1946873 |
| + | 1947229 |
| + | 1947635 |
| + | 1947637 |
| + | 1947639 |
| + | 1948406 |
| - | 1948462 |
| + | 1954630 |

| | |
|---|---|
| - | 1954960 |
| + | 1954977 |
| + | 1967079 |
| + | 1968124 |
| + | 1969545 |
| + | 1970459 |
| - | 1974054 |
| + | 1974316 |
| + | 1978906 |
| + | 1988330 |
| + | 1994333 |
| + | 1997203 |
| + | 1997561 |
| - | 1997665 |
| - | 1998056 |
| - | 1999450 |
| - | 2000856 |
| + | 2001354 |
| + | 2002176 |
| - | 2002464 |
| + | 2002601 |
| + | 2002603 |
| - | 2003990 |
| - | 2004520 |
| - | 2004585 |
| + | 2004645 |
| - | 2007446 |
| - | 2007448 |
| - | 2007450 |
| + | 2007490 |
| + | 2010974 |
| - | 2014688 |
| + | 2014723 |
| + | 2014738 |
| + | 2016349 |
| - | 2017746 |
| - | 2018290 |
| - | 2018427 |
| - | 2018486 |
| - | 2019302 |
| - | 2019305 |
| + | 2019402 |
| - | 2019422 |
| - | 2019792 |
| - | 2019794 |
| - | 2020114 |
| + | 2021575 |
| - | 2023320 |
| - | 2025283 |
| + | 2025366 |

| | |
|---|---|
| + | 2025368 |
| + | 2027483 |
| + | 2027486 |
| + | 2028267 |
| - | 2028666 |
| + | 2030171 |
| + | 2030174 |
| - | 2031197 |
| + | 2031249 |
| - | 2033712 |
| - | 2034435 |
| - | 2035926 |
| - | 2035934 |
| - | 2037453 |
| - | 2039408 |
| - | 2039826 |
| - | 2039827 |
| + | 2040597 |
| + | 2042156 |
| + | 2042429 |
| + | 2043160 |
| - | 2044951 |
| - | 2045865 |
| + | 2045895 |
| + | 2045989 |
| - | 2047187 |
| + | 2048482 |
| + | 2048483 |
| - | 2048968 |
| + | 2052375 |
| + | 2053990 |
| + | 2053992 |
| - | 2054033 |
| - | 2055337 |
| + | 2055650 |
| - | 2056148 |
| - | 2056149 |
| + | 2056250 |
| - | 2059653 |
| + | 2060789 |
| + | 2060791 |
| + | 2062113 |
| + | 2062114 |
| + | 2063026 |
| + | 2065005 |
| - | 2065741 |
| - | 2069168 |
| - | 2069680 |
| - | 2069682 |
| + | 2069820 |

| | |
|---|---|
| + | 2069821 |
| - | 2070127 |
| - | 2070128 |
| + | 2070176 |
| - | 2071033 |
| - | 2071114 |
| - | 2071115 |
| - | 2071332 |
| - | 2071441 |
| - | 2071882 |
| - | 2072053 |
| - | 2072248 |
| - | 2072652 |
| - | 2073031 |
| - | 2073582 |
| - | 2074605 |
| - | 2076074 |
| + | 2076128 |
| + | 2077725 |
| - | 2077999 |
| + | 2079093 |
| + | 2079095 |
| - | 2081123 |
| - | 2081124 |
| - | 2081125 |
| - | 2082488 |
| - | 2083033 |
| - | 2084040 |
| - | 2085132 |
| + | 2085267 |
| + | 2085268 |
| + | 2086481 |
| - | 2086635 |
| - | 2087036 |
| - | 2087550 |
| + | 2087661 |
| + | 2088231 |
| + | 2089368 |
| + | 2089370 |
| + | 2090544 |
| + | 2090700 |
| + | 2092522 |
| - | 2094839 |
| - | 2095817 |
| - | 2096111 |
| - | 2096114 |
| - | 2097013 |
| - | 2097093 |
| - | 2097634 |
| - | 2097635 |

| | |
|---|---|
| + | 2098068 |
| + | 2099033 |
| - | 2099862 |
| - | 2099956 |
| - | 2100375 |
| - | 2100377 |
| + | 2100459 |
| + | 2100461 |
| - | 2101024 |
| + | 2101825 |
| + | 2101951 |
| - | 2104611 |
| - | 2106307 |
| + | 2106428 |
| + | 2110928 |
| - | 2111730 |
| - | 2114683 |
| - | 2115647 |
| - | 2115738 |
| + | 2116538 |
| - | 2116957 |
| - | 2116958 |
| - | 2118304 |
| - | 2119265 |
| + | 2119365 |
| + | 2120410 |
| + | 2121909 |
| - | 2122301 |
| - | 2122979 |
| - | 2123947 |
| - | 2123949 |
| - | 2126321 |
| + | 2126453 |
| + | 2127744 |
| - | 2127780 |
| - | 2128043 |
| - | 2130027 |
| + | 2130148 |
| + | 2130151 |
| + | 2130424 |
| + | 2130871 |
| + | 2133108 |
| - | 2133324 |
| - | 2133325 |
| - | 2135494 |
| - | 2136053 |
| - | 2136247 |
| - | 2136248 |
| - | 2136875 |
| - | 2137623 |

| | |
|---|---|
| + | 2138196 |
| + | 2138607 |
| - | 2139122 |
| - | 2139123 |
| + | 2140161 |
| + | 2140993 |
| + | 2142085 |
| - | 2146487 |
| - | 2146850 |
| - | 2147472 |
| + | 2147510 |
| - | 2150461 |
| - | 2151283 |
| + | 2152300 |
| + | 2152911 |
| + | 2154032 |
| - | 2155009 |
| - | 2156709 |
| + | 2158416 |
| - | 2159189 |
| - | 2159190 |
| + | 2162041 |
| + | 2162042 |
| + | 2164102 |
| - | 2164895 |
| - | 2166554 |
| - | 2170266 |
| - | 2170746 |
| - | 2174793 |
| - | 2175517 |
| - | 2178652 |
| + | 2182343 |
| - | 2182374 |
| + | 2183952 |
| + | 2184342 |
| - | 2186837 |
| - | 2187326 |
| - | 2187329 |
| - | 2187330 |
| - | 2187342 |
| - | 2187503 |
| - | 2187956 |
| - | 2188147 |
| - | 2188150 |
| + | 2190677 |
| + | 2193787 |
| + | 2195143 |
| + | 2195148 |
| + | 2195953 |
| + | 2196160 |

| | |
|---|---|
| - | 2196632 |
| - | 2203667 |
| + | 2203679 |
| + | 2203738 |
| - | 2204065 |
| - | 2206603 |
| - | 2208126 |
| - | 2208300 |
| + | 2208374 |
| - | 2208570 |
| - | 2208813 |
| - | 2209000 |
| - | 2210191 |
| - | 2212502 |
| - | 2212503 |
| + | 2212535 |
| + | 2212538 |
| + | 2212541 |
| + | 2212835 |
| - | 2213974 |
| - | 2214953 |
| - | 2215085 |
| - | 2215882 |
| - | 2216613 |
| - | 2216728 |
| - | 2216813 |
| + | 2217339 |
| - | 2217681 |
| - | 2219484 |
| - | 2219485 |
| - | 2219525 |
| - | 2219568 |
| + | 2219744 |
| + | 2219905 |
| - | 2219981 |
| - | 2221126 |
| - | 2221458 |
| + | 2221646 |
| + | 2221800 |
| + | 2222320 |
| - | 2224790 |
| + | 2227159 |
| - | 2227579 |
| - | 2229197 |
| - | 2230042 |
| - | 2230046 |
| + | 2232697 |
| + | 2233234 |
| - | 2234346 |
| - | 2236831 |

| | |
|---|---|
| + | 2237525 |
| - | 2238454 |
| - | 2240143 |
| - | 2243112 |
| - | 2245469 |
| - | 2245591 |
| - | 2247590 |
| - | 2247689 |
| + | 2247876 |
| + | 2248389 |
| + | 2248390 |
| + | 2248391 |
| + | 2248992 |
| + | 2255939 |
| + | 2257051 |
| + | 2261053 |
| - | 2264585 |
| + | 2264891 |
| - | 2265410 |
| - | 2269728 |
| - | 2269730 |
| - | 2269733 |
| - | 2270333 |
| - | 2270334 |
| - | 2272001 |
| - | 2273392 |
| + | 2273533 |
| + | 2273534 |
| + | 2273882 |
| - | 2273884 |
| + | 2275750 |
| - | 2276113 |
| - | 2276896 |
| - | 2277446 |
| - | 2277580 |
| - | 2279711 |
| - | 2279713 |
| - | 2279715 |
| + | 2279928 |
| + | 2279948 |
| + | 2280193 |
| + | 2281800 |
| - | 2282977 |
| - | 2282978 |
| - | 2282979 |
| - | 2282981 |
| - | 2283126 |
| - | 2283751 |
| - | 2284611 |
| + | 2284721 |

| | |
|---|---|
| + | 2286306 |
| - | 2288092 |
| - | 2288094 |
| + | 2288097 |
| + | 2288100 |
| + | 2288102 |
| + | 2288662 |
| - | 2290451 |
| - | 2291650 |
| - | 2291651 |
| - | 2291762 |
| - | 2292342 |
| - | 2294074 |
| - | 2294076 |
| + | 2294261 |
| - | 2296563 |
| - | 2297928 |
| + | 2298636 |
| - | 2299182 |
| - | 2299183 |
| - | 2299184 |
| + | 2299232 |
| + | 2299294 |
| - | 2299345 |
| - | 2300685 |
| - | 2300686 |
| - | 2302618 |
| - | 2302946 |
| - | 2303883 |
| - | 2303885 |
| - | 2303889 |
| + | 2304260 |
| - | 2304354 |
| - | 2305059 |
| - | 2305221 |
| + | 2305272 |
| + | 2306335 |
| + | 2306449 |
| + | 2306451 |
| + | 2307817 |
| + | 2307820 |
| - | 2308403 |
| + | 2309042 |
| + | 2309699 |
| - | 2309764 |
| - | 2310158 |
| + | 2310867 |
| - | 2311909 |
| - | 2311910 |
| - | 2312487 |

| | |
|---|---|
| - | 2316172 |
| - | 2316407 |
| + | 2316555 |
| + | 2317231 |
| - | 2318001 |
| - | 2318449 |
| - | 2320209 |
| - | 2320211 |
| + | 2320972 |
| - | 2321098 |
| - | 2321888 |
| - | 2325682 |
| + | 2325791 |
| - | 2327942 |
| - | 2329445 |
| - | 2329447 |
| - | 2329839 |
| - | 2331267 |
| - | 2331720 |
| - | 2331723 |
| - | 2332102 |
| - | 2332105 |
| - | 2332412 |
| - | 2333215 |
| - | 2333235 |
| + | 2333769 |
| - | 2334645 |
| + | 2336345 |
| - | 2336848 |
| - | 2337016 |
| - | 2337485 |
| - | 2337763 |
| - | 2338532 |
| + | 2338554 |
| + | 2338555 |
| - | 2339193 |
| - | 2339194 |
| - | 2339498 |
| + | 2339963 |
| + | 2339964 |
| + | 2340940 |
| - | 2341203 |
| + | 2341377 |
| + | 2341378 |
| + | 2343214 |
| - | 2345120 |
| - | 2346355 |
| - | 2347581 |
| - | 2347583 |
| - | 2349299 |

| | |
|---|---|
| - | 2350160 |
| - | 2350798 |
| - | 2351773 |
| - | 2352494 |
| - | 2352968 |
| - | 2353003 |
| - | 2353829 |
| + | 2357671 |
| - | 2360512 |
| - | 2360581 |
| + | 2360597 |
| - | 2363071 |
| - | 2365160 |
| - | 2365695 |
| - | 2366310 |
| - | 2366312 |
| + | 2367706 |
| - | 2367755 |
| - | 2371504 |
| - | 2372000 |
| - | 2372001 |
| - | 2372002 |
| + | 2372532 |
| - | 2375670 |
| - | 2377619 |
| - | 2378143 |
| - | 2378243 |
| - | 2379900 |
| - | 2380309 |
| - | 2381848 |
| - | 2381850 |
| - | 2384446 |
| - | 2384447 |
| - | 2386034 |
| - | 2386036 |
| - | 2386038 |
| - | 2388850 |
| - | 2388852 |
| - | 2388853 |
| - | 2391629 |
| - | 2392214 |
| + | 2393237 |
| + | 2393258 |
| - | 2395955 |
| - | 2395958 |
| - | 2395961 |
| - | 2396745 |
| - | 2396747 |
| - | 2397704 |
| - | 2398134 |

| | |
|---|---|
| - | 2398941 |
| + | 2399911 |
| - | 2400610 |
| - | 2400891 |
| - | 2401581 |
| - | 2402022 |
| - | 2403204 |
| - | 2403396 |
| - | 2403397 |
| - | 2404167 |
| - | 2404171 |
| - | 2405056 |
| + | 2405599 |
| - | 2406614 |
| - | 2406739 |
| - | 2406869 |
| + | 2407568 |
| - | 2407758 |
| + | 2408793 |
| - | 2409506 |
| - | 2409508 |
| + | 2409648 |
| + | 2409650 |
| + | 2409651 |
| - | 2410890 |
| - | 2410891 |
| - | 2410892 |
| - | 2410893 |
| + | 2411049 |
| + | 2411051 |
| + | 2412057 |
| - | 2413501 |
| - | 2415262 |
| - | 2417351 |
| - | 2417928 |
| - | 2421375 |
| - | 2422065 |
| - | 2422184 |
| - | 2422185 |
| - | 2422273 |
| + | 2422456 |
| - | 2424868 |
| - | 2426450 |
| - | 2426633 |
| - | 2426635 |
| - | 2426824 |
| + | 2426845 |
| + | 2426847 |
| - | 2427848 |
| - | 2431637 |

| | |
|---|---|
| - | 2431640 |
| - | 2431883 |
| - | 2432169 |
| - | 2432691 |
| - | 2432900 |
| - | 2432901 |
| - | 2433558 |
| - | 2433559 |
| - | 2434206 |
| - | 2435863 |
| - | 2438330 |
| - | 2438335 |
| + | 2439095 |
| + | 2441309 |
| - | 2445025 |
| - | 2448454 |
| - | 2448483 |
| - | 2449621 |
| - | 2449749 |
| - | 2450317 |
| - | 2450886 |
| - | 2451387 |
| - | 2452842 |
| - | 2452843 |
| - | 2454035 |
| - | 2454258 |
| - | 2454261 |
| - | 2456879 |
| - | 2456943 |
| + | 2456966 |
| - | 2457352 |
| - | 2459104 |
| + | 2459298 |
| + | 2459301 |
| - | 2460528 |
| - | 2460529 |
| - | 2460530 |
| - | 2461941 |
| - | 2462225 |
| - | 2462226 |
| + | 2462337 |
| - | 2464097 |
| + | 2465941 |
| - | 2466595 |
| + | 2466691 |
| - | 2468528 |
| - | 2469543 |
| - | 2471799 |
| - | 2472772 |
| + | 2472817 |

| | |
|---|---|
| + | 2472881 |
| - | 2472914 |
| + | 2473586 |
| - | 2475728 |
| + | 2475816 |
| + | 2475936 |
| + | 2476943 |
| - | 2477955 |
| - | 2477957 |
| - | 2478202 |
| - | 2478759 |
| - | 2478956 |
| + | 2479099 |
| + | 2479102 |
| + | 2479103 |
| - | 2482226 |
| - | 2482228 |
| - | 2482229 |
| - | 2483362 |
| + | 2483571 |
| + | 2483572 |
| - | 2483777 |
| - | 2483780 |
| + | 2485060 |
| + | 2486589 |
| - | 2486646 |
| - | 2487921 |
| - | 2487922 |
| + | 2488818 |
| - | 2489325 |
| - | 2491848 |
| - | 2492849 |
| - | 2493527 |
| + | 2493635 |
| + | 2494628 |
| + | 2495402 |
| + | 2495883 |
| - | 2496070 |
| - | 2496594 |
| + | 2497230 |
| - | 2504692 |
| - | 2506929 |
| - | 2506931 |
| + | 2506989 |
| + | 2506990 |
| - | 2508518 |
| - | 2515931 |
| + | 2516202 |
| + | 2516414 |
| - | 2517741 |

| | |
|---|---|
| - | 2518869 |
| - | 2519030 |
| - | 2519032 |
| - | 2519033 |
| - | 2519034 |
| - | 2522803 |
| - | 2523362 |
| - | 2523677 |
| - | 2525740 |
| - | 2526555 |
| + | 2527042 |
| + | 2527619 |
| - | 2529707 |
| - | 2529709 |
| - | 2531238 |
| - | 2531240 |
| - | 2532258 |
| + | 2532952 |
| - | 2536050 |
| - | 2539348 |
| - | 2539349 |
| - | 2539587 |
| - | 2540252 |
| - | 2540402 |
| + | 2540909 |
| + | 2542576 |
| - | 2543905 |
| - | 2544830 |
| + | 2544922 |
| - | 2549665 |
| + | 2552313 |
| + | 2552322 |
| + | 2552324 |
| + | 2552634 |
| + | 2552637 |
| + | 2552638 |
| + | 2554834 |
| - | 2555309 |
| - | 2557230 |
| + | 2559626 |
| - | 2560096 |
| + | 2560350 |
| - | 2561471 |
| - | 2561472 |
| + | 2561560 |
| - | 2562972 |
| + | 2563159 |
| - | 2563350 |
| - | 2564439 |
| - | 2564440 |

| | |
|---|---|
| - | 2564441 |
| - | 2564442 |
| + | 2564610 |
| + | 2564611 |
| + | 2564612 |
| - | 2565582 |
| - | 2565590 |
| + | 2565674 |
| - | 2567324 |
| - | 2568033 |
| - | 2569088 |
| + | 2569639 |
| - | 2570554 |
| - | 2571824 |
| - | 2573457 |
| - | 2573458 |
| - | 2574596 |
| + | 2574614 |
| + | 2576169 |
| - | 2576225 |
| + | 2576337 |
| - | 2581658 |
| + | 2582016 |
| - | 2582334 |
| - | 2586098 |
| - | 2586170 |
| + | 2586779 |
| + | 2586781 |
| + | 2586783 |
| - | 2588451 |
| + | 2588454 |
| + | 2589052 |
| + | 2589053 |
| + | 2589660 |
| + | 2590781 |
| - | 2590818 |
| + | 2590925 |
| - | 2591959 |
| - | 2592924 |
| - | 2592927 |
| + | 2594259 |
| - | 2595538 |
| - | 2595539 |
| + | 2596454 |
| - | 2598477 |
| - | 2599364 |
| - | 2599365 |
| - | 2599923 |
| - | 2599925 |
| + | 2600062 |

| | |
|---|---|
| + | 2600156 |
| - | 2601529 |
| + | 2602969 |
| - | 2603354 |
| - | 2603840 |
| - | 2603842 |
| - | 2604026 |
| - | 2604088 |
| - | 2604317 |
| + | 2604328 |
| - | 2604663 |
| - | 2605632 |
| - | 2605635 |
| + | 2605855 |
| - | 2608915 |
| - | 2608917 |
| - | 2608918 |
| - | 2609636 |
| - | 2609970 |
| + | 2610028 |
| - | 2611375 |
| + | 2611441 |
| - | 2611480 |
| - | 2611484 |
| + | 2612441 |
| - | 2613207 |
| - | 2613280 |
| - | 2613403 |
| - | 2614087 |
| - | 2614669 |
| - | 2614742 |
| - | 2615521 |
| - | 2615602 |
| - | 2616948 |
| - | 2618573 |
| - | 2619817 |
| - | 2619818 |
| - | 2620607 |
| + | 2620629 |
| - | 2623588 |
| - | 2625941 |
| - | 2625942 |
| - | 2626580 |
| - | 2628605 |
| - | 2628791 |
| - | 2629682 |
| + | 2630798 |
| - | 2632785 |
| - | 2633488 |
| + | 2635744 |

| | |
|---|---|
| + | 2635746 |
| + | 2635747 |
| - | 2637169 |
| - | 2637170 |
| - | 2637171 |
| - | 2637331 |
| + | 2637352 |
| + | 2637581 |
| + | 2638489 |
| - | 2640475 |
| - | 2642528 |
| - | 2643861 |
| - | 2643863 |
| - | 2644799 |
| - | 2644908 |
| + | 2645957 |
| - | 2647278 |
| - | 2647360 |
| + | 2647372 |
| - | 2647663 |
| - | 2647664 |
| - | 2647666 |
| - | 2647949 |
| - | 2647950 |
| - | 2647951 |
| - | 2648672 |
| + | 2649069 |
| + | 2649789 |
| - | 2650852 |
| + | 2651598 |
| + | 2653578 |
| - | 2654109 |
| + | 2654436 |
| - | 2654906 |
| + | 2656024 |
| - | 2656757 |
| - | 2657667 |
| - | 2657670 |
| + | 2659180 |
| + | 2659183 |
| - | 2660113 |
| + | 2660136 |
| + | 2660255 |
| + | 2660257 |
| + | 2660956 |
| + | 2661075 |
| - | 2661475 |
| + | 2661923 |
| + | 2661925 |
| + | 2663489 |

| | |
|---|---|
| - | 2663971 |
| - | 2664304 |
| - | 2664439 |
| - | 2665074 |
| + | 2665554 |
| + | 2666494 |
| + | 2667684 |
| - | 2673394 |
| + | 2674118 |
| + | 2674120 |
| + | 2678182 |
| + | 2678193 |
| + | 2678526 |
| - | 2678564 |
| - | 2678586 |
| - | 2678588 |
| + | 2678730 |
| + | 2678731 |
| - | 2679075 |
| - | 2682324 |
| - | 2683022 |
| + | 2683079 |
| - | 2685518 |
| - | 2685838 |
| - | 2688903 |
| + | 2689968 |
| - | 2691215 |
| + | 2691565 |
| - | 2691617 |
| + | 2691620 |
| - | 2692836 |
| + | 2692882 |
| + | 2697040 |
| + | 2698640 |
| + | 2698862 |
| - | 2700621 |
| - | 2700622 |
| + | 2700698 |
| + | 2701560 |
| - | 2702080 |
| - | 2705202 |
| + | 2705333 |
| - | 2705826 |
| + | 2705896 |
| + | 2706387 |
| - | 2707407 |
| + | 2708582 |
| - | 2708785 |
| - | 2708786 |
| - | 2708787 |

| | |
|---|---|
| - | 2709458 |
| - | 2709459 |
| + | 2709861 |
| - | 2714339 |
| - | 2714342 |
| - | 2714402 |
| - | 2714622 |
| - | 2715427 |
| - | 2715451 |
| - | 2715452 |
| - | 2716894 |
| + | 2716904 |
| - | 2720334 |
| - | 2720336 |
| - | 2720338 |
| - | 2721856 |
| + | 2721873 |
| + | 2722252 |
| + | 2724651 |
| - | 2725319 |
| + | 2725791 |
| - | 2726163 |
| - | 2727181 |
| - | 2730268 |
| - | 2730854 |
| + | 2732383 |
| - | 2733575 |
| - | 2733578 |
| - | 2733579 |
| - | 2733581 |
| - | 2734358 |
| - | 2734360 |
| - | 2735499 |
| - | 2735501 |
| - | 2736620 |
| + | 2736843 |
| + | 2736844 |
| + | 2737418 |
| + | 2738281 |
| - | 2738377 |
| - | 2739296 |
| + | 2739444 |
| + | 2739447 |
| + | 2740428 |
| - | 2742146 |
| - | 2742822 |
| - | 2742854 |
| - | 2744047 |
| - | 2745518 |
| - | 2746020 |

| | |
|---|---|
| - | 2746488 |
| + | 2747618 |
| - | 2748275 |
| - | 2748848 |
| + | 2748933 |
| + | 2749059 |
| + | 2749524 |
| + | 2749633 |
| - | 2751204 |
| - | 2751747 |
| - | 2751749 |
| - | 2752594 |
| - | 2753703 |
| - | 2755828 |
| - | 2755842 |
| - | 2755957 |
| + | 2756239 |
| + | 2757466 |
| + | 2757468 |
| - | 2762862 |
| + | 2763839 |
| + | 2764659 |
| - | 2766494 |
| - | 2768732 |
| - | 2768735 |
| + | 2768790 |
| + | 2768791 |
| + | 2769802 |
| - | 2770909 |
| + | 2773340 |
| - | 2777756 |
| - | 2778505 |
| + | 2778871 |
| - | 2778872 |
| - | 2779367 |
| + | 2779527 |
| - | 2780279 |
| - | 2780477 |
| - | 2780979 |
| + | 2781016 |
| - | 2784945 |
| - | 2784947 |
| - | 2788055 |
| - | 2788056 |
| + | 2788653 |
| - | 2789716 |
| - | 2789896 |
| - | 2790757 |
| + | 2791054 |
| + | 2791055 |

| | |
|---|---|
| - | 2791896 |
| - | 2792083 |
| - | 2792085 |
| - | 2792205 |
| - | 2792206 |
| + | 2792221 |
| - | 2794099 |
| - | 2795825 |
| - | 2796985 |
| - | 2797545 |
| - | 2797548 |
| - | 2798146 |
| - | 2798166 |
| - | 2800579 |
| - | 2801108 |
| - | 2801110 |
| - | 2803064 |
| + | 2806061 |
| - | 2806554 |
| - | 2809339 |
| - | 2811050 |
| - | 2812217 |
| - | 2812218 |
| + | 2812317 |
| - | 2814018 |
| - | 2814691 |
| - | 2814692 |
| + | 2817849 |
| - | 2818161 |
| - | 2818162 |
| - | 2818163 |
| + | 2818176 |
| + | 2818177 |
| - | 2818390 |
| + | 2818457 |
| + | 2819462 |
| - | 2819744 |
| - | 2819745 |
| - | 2822034 |
| - | 2822765 |
| - | 2822767 |
| - | 2823607 |
| - | 2823609 |
| + | 2825528 |
| - | 2825795 |
| - | 2825796 |
| - | 2826195 |
| - | 2826196 |
| - | 2826664 |
| - | 2826817 |

| | |
|---|---|
| - | 2827474 |
| - | 2828567 |
| + | 2828703 |
| + | 2828704 |
| + | 2828708 |
| - | 2829197 |
| - | 2829198 |
| - | 2829352 |
| - | 2829514 |
| - | 2829516 |
| - | 2830177 |
| - | 2830436 |
| - | 2832136 |
| - | 2832827 |
| - | 2833215 |
| - | 2833534 |
| + | 2835017 |
| - | 2835821 |
| - | 2836257 |
| - | 2836790 |
| - | 2836792 |
| - | 2836825 |
| - | 2838153 |
| + | 2838458 |
| - | 2838802 |
| - | 2839913 |
| - | 2839914 |
| - | 2839915 |
| + | 2840036 |
| + | 2841377 |
| + | 2841574 |
| - | 2841661 |
| + | 2842840 |
| - | 2843853 |
| - | 2843979 |
| - | 2845475 |
| - | 2849497 |
| + | 2849532 |
| - | 2849626 |
| + | 2850505 |
| - | 2854611 |
| - | 2854613 |
| - | 2854615 |
| - | 2854618 |
| + | 2854624 |
| - | 2854803 |
| - | 2855930 |
| - | 2855935 |
| - | 2857086 |
| - | 2857202 |

| | |
|---|---|
| + | 2858238 |
| - | 2859279 |
| - | 2859282 |
| - | 2859843 |
| - | 2860099 |
| - | 2861801 |
| - | 2861802 |
| - | 2862995 |
| + | 2863067 |
| - | 2863182 |
| + | 2863776 |
| - | 2864340 |
| + | 2865160 |
| - | 2865906 |
| - | 2866814 |
| - | 2869618 |
| - | 2869620 |
| - | 2869627 |
| - | 2874159 |
| - | 2874354 |
| - | 2876437 |
| + | 2877062 |
| - | 2879199 |
| - | 2879200 |
| - | 2879206 |
| + | 2879271 |
| - | 2879293 |
| - | 2879700 |
| - | 2880801 |
| - | 2880803 |
| - | 2881190 |
| - | 2881858 |
| - | 2882891 |
| - | 2882894 |
| - | 2886122 |
| - | 2886124 |
| - | 2886126 |
| - | 2886295 |
| + | 2887375 |
| - | 2887624 |
| - | 2887768 |
| - | 2887772 |
| - | 2887774 |
| + | 2888291 |
| + | 2888576 |
| - | 2888850 |
| - | 2888852 |
| - | 2889011 |
| + | 2892565 |
| + | 2892567 |

| | |
|---|---|
| + | 2892568 |
| - | 2897453 |
| - | 2897455 |
| - | 2897605 |
| + | 2898242 |
| + | 2898891 |
| - | 2899929 |
| - | 2899932 |
| - | 2900101 |
| - | 2901359 |
| - | 2901886 |
| - | 2903129 |
| - | 2904528 |
| + | 2905019 |
| - | 2905054 |
| + | 2905403 |
| - | 2908087 |
| + | 2908544 |
| - | 2908827 |
| + | 2908930 |
| + | 2909010 |
| - | 2909465 |
| + | 2910745 |
| - | 2911077 |
| - | 2911078 |
| - | 2911081 |
| - | 2911087 |
| - | 2912917 |
| - | 2913368 |
| - | 2913369 |
| - | 2913583 |
| - | 2914847 |
| - | 2915247 |
| - | 2915248 |
| - | 2918585 |
| + | 2919702 |
| - | 2920359 |
| - | 2920362 |
| - | 2920603 |
| - | 2920937 |
| - | 2920938 |
| + | 2923650 |
| - | 2925052 |
| - | 2925056 |
| - | 2925058 |
| - | 2925925 |
| - | 2925926 |
| + | 2925992 |
| + | 2925998 |
| - | 2926706 |

| | |
|---|---|
| + | 2928544 |
| - | 2930791 |
| - | 2930793 |
| + | 2930796 |
| - | 2931602 |
| + | 2931663 |
| + | 2931958 |
| + | 2933154 |
| + | 2935226 |
| - | 2936317 |
| - | 2936486 |
| + | 2936580 |
| + | 2938228 |
| + | 2938861 |
| - | 2939523 |
| + | 2941424 |
| + | 2941425 |
| - | 2942270 |
| - | 2942460 |
| + | 2946783 |
| - | 2948165 |
| - | 2948972 |
| - | 2950139 |
| - | 2951334 |
| + | 2951403 |
| + | 2951458 |
| + | 2951459 |
| - | 2953607 |
| - | 2953610 |
| - | 2953613 |
| + | 2953778 |
| - | 2955673 |
| - | 2956734 |
| - | 2958287 |
| - | 2958367 |
| - | 2958369 |
| + | 2958386 |
| + | 2958388 |
| - | 2961492 |
| - | 2961493 |
| - | 2961494 |
| - | 2961768 |
| - | 2963645 |
| - | 2964901 |
| - | 2965167 |
| - | 2965412 |
| - | 2965595 |
| - | 2965597 |
| - | 2966279 |
| - | 2966280 |

| | |
|---|---|
| - | 2966931 |
| - | 2966932 |
| + | 2966953 |
| + | 2967209 |
| - | 2968100 |
| - | 2968756 |
| + | 2969058 |
| - | 2974083 |
| - | 2975881 |
| - | 2975882 |
| - | 2976062 |
| - | 2978592 |
| - | 2980773 |
| - | 2981073 |
| - | 2981074 |
| - | 2981526 |
| + | 2982189 |
| - | 2982463 |
| - | 2986651 |
| + | 2986675 |
| - | 2987163 |
| - | 2987640 |
| - | 2987641 |
| - | 2987924 |
| - | 2988612 |
| + | 2989632 |
| - | 2989640 |
| - | 2990958 |
| - | 2991239 |
| - | 2992492 |
| - | 2994422 |
| - | 2994649 |
| - | 2994650 |
| + | 2994747 |
| - | 2995363 |
| + | 2995664 |
| + | 2995666 |
| + | 2995956 |
| + | 2996678 |
| + | 2996827 |
| + | 2996951 |
| + | 2997155 |
| + | 2997156 |
| - | 2997250 |
| - | 2998648 |
| - | 3000277 |
| + | 3004290 |
| + | 3006508 |
| + | 3008859 |
| - | 3010625 |

| | |
|---|---|
| - | 3011119 |
| - | 3011475 |
| - | 3011478 |
| - | 3011481 |
| - | 3011740 |
| - | 3012769 |
| - | 3014419 |
| - | 3015054 |
| - | 3016390 |
| - | 3016394 |
| - | 3017764 |
| - | 3018224 |
| - | 3018226 |
| - | 3018228 |
| - | 3021073 |
| - | 3021074 |
| + | 3021163 |
| + | 3021164 |
| - | 3021164 |
| - | 3022283 |
| - | 3025346 |
| - | 3025671 |
| - | 3025823 |
| + | 3026666 |
| - | 3027905 |
| - | 3028137 |
| - | 3028140 |
| - | 3028144 |
| + | 3028222 |
| + | 3028225 |
| - | 3028312 |
| + | 3028342 |
| - | 3029325 |
| - | 3031513 |
| - | 3031514 |
| + | 3031528 |
| + | 3031534 |
| - | 3032090 |
| - | 3032092 |
| - | 3032479 |
| + | 3032761 |
| + | 3032776 |
| + | 3032917 |
| - | 3033060 |
| + | 3034941 |
| + | 3034945 |
| + | 3035546 |
| + | 3035548 |
| - | 3036527 |
| + | 3037599 |

| | |
|---|---|
| + | 3037723 |
| - | 3038168 |
| - | 3039961 |
| + | 3040051 |
| + | 3040982 |
| + | 3041493 |
| - | 3041920 |
| + | 3043185 |
| - | 3045331 |
| - | 3045334 |
| - | 3046565 |
| - | 3046567 |
| - | 3046674 |
| - | 3047160 |
| - | 3047392 |
| - | 3048040 |
| - | 3048089 |
| - | 3049684 |
| - | 3049685 |
| - | 3049829 |
| - | 3050622 |
| + | 3052449 |
| - | 3052618 |
| - | 3052619 |
| - | 3053120 |
| - | 3053489 |
| - | 3054048 |
| + | 3054623 |
| - | 3054637 |
| - | 3054638 |
| + | 3054698 |
| - | 3056344 |
| + | 3056454 |
| + | 3056918 |
| + | 3057344 |
| + | 3058593 |
| - | 3059490 |
| - | 3060215 |
| - | 3060216 |
| - | 3065492 |
| - | 3065493 |
| + | 3065674 |
| - | 3066326 |
| + | 3066419 |
| + | 3066421 |
| + | 3068873 |
| + | 3068875 |
| - | 3069231 |
| - | 3069232 |
| - | 3069233 |

| | |
|---|---|
| + | 3069249 |
| - | 3072125 |
| + | 3072195 |
| + | 3072289 |
| + | 3074964 |
| - | 3075199 |
| + | 3075384 |
| + | 3075835 |
| + | 3076698 |
| + | 3077270 |
| + | 3078636 |
| + | 3081966 |
| + | 3083419 |
| - | 3088261 |
| + | 3088307 |
| - | 3089169 |
| + | 3089983 |
| + | 3092259 |
| - | 3096107 |
| - | 3096588 |
| + | 3096732 |
| - | 3100272 |
| + | 3101619 |
| + | 3101772 |
| - | 3102538 |
| - | 3105381 |
| + | 3105440 |
| - | 3106051 |
| + | 3106141 |
| + | 3107147 |
| - | 3108507 |
| + | 3108592 |
| + | 3109706 |
| - | 3112113 |
| + | 3112123 |
| + | 3113529 |
| + | 3115274 |
| - | 3116195 |
| - | 3119482 |
| - | 3119873 |
| + | 3119968 |
| + | 3123122 |
| - | 3123705 |
| - | 3124015 |
| - | 3124134 |
| - | 3124741 |
| - | 3126596 |
| - | 3127715 |
| - | 3129342 |
| - | 3129345 |

| | |
|---|---|
| - | 3129346 |
| + | 3129491 |
| - | 3131758 |
| + | 3133044 |
| - | 3135491 |
| - | 3135493 |
| - | 3136706 |
| - | 3136708 |
| + | 3136780 |
| - | 3136884 |
| + | 3136895 |
| - | 3137402 |
| - | 3137404 |
| - | 3137997 |
| + | 3138077 |
| - | 3138502 |
| - | 3138912 |
| + | 3138957 |
| - | 3139257 |
| - | 3139296 |
| - | 3139297 |
| - | 3139381 |
| + | 3140811 |
| + | 3141285 |
| - | 3146079 |
| - | 3148257 |
| - | 3148461 |
| - | 3149834 |
| - | 3150272 |
| + | 3151569 |
| - | 3152742 |
| - | 3153815 |
| - | 3163197 |
| + | 3163220 |
| + | 3163341 |
| + | 3165105 |
| - | 3172320 |
| - | 3172676 |
| - | 3172786 |
| - | 3173040 |
| - | 3173590 |
| - | 3173935 |
| - | 3173956 |
| - | 3176916 |
| - | 3177385 |
| - | 3178816 |
| - | 3178911 |
| + | 3179093 |
| - | 3180521 |
| - | 3182585 |

| | |
|---|---|
| + | 3182672 |
| + | 3182694 |
| - | 3183818 |
| - | 3184415 |
| - | 3184847 |
| + | 3186712 |
| + | 3187420 |
| - | 3187466 |
| + | 3188175 |
| + | 3188177 |
| - | 3189832 |
| - | 3190045 |
| + | 3190683 |
| - | 3190768 |
| - | 3190770 |
| - | 3190875 |
| - | 3191731 |
| + | 3191981 |
| - | 3192486 |
| - | 3192932 |
| - | 3193540 |
| - | 3194539 |
| - | 3194839 |
| + | 3195379 |
| - | 3196753 |
| - | 3196756 |
| + | 3196879 |
| + | 3196881 |
| - | 3197365 |
| - | 3197772 |
| - | 3201839 |
| - | 3201841 |
| - | 3203964 |
| - | 3205818 |
| - | 3206079 |
| - | 3207348 |
| - | 3208178 |
| - | 3210298 |
| - | 3211695 |
| - | 3212457 |
| - | 3213603 |
| - | 3214299 |
| - | 3215346 |
| - | 3215649 |
| + | 3215854 |
| - | 3218210 |
| + | 3218222 |
| + | 3218971 |
| - | 3219831 |
| - | 3220103 |

| | |
|---|---|
| + | 3221931 |
| - | 3222115 |
| + | 3222116 |
| - | 3222122 |
| - | 3223054 |
| - | 3223404 |
| - | 3224650 |
| - | 3224700 |
| + | 3224754 |
| + | 3224756 |
| - | 3225612 |
| - | 3225613 |
| - | 3225615 |
| + | 3225710 |
| + | 3226843 |
| - | 3227279 |
| - | 3227466 |
| - | 3228381 |
| + | 3228717 |
| + | 3228718 |
| - | 3228719 |
| - | 3229359 |
| + | 3230037 |
| + | 3230039 |
| - | 3230287 |
| + | 3231386 |
| - | 3232467 |
| + | 3232477 |
| + | 3232582 |
| - | 3233700 |
| + | 3233891 |
| - | 3234552 |
| + | 3235896 |
| - | 3236442 |
| + | 3236461 |
| - | 3237006 |
| - | 3237007 |
| - | 3237010 |
| + | 3237128 |
| + | 3238143 |
| + | 3238145 |
| - | 3238242 |
| + | 3239659 |
| - | 3241620 |
| + | 3243043 |
| + | 3244142 |
| + | 3244742 |
| + | 3244744 |
| + | 3244745 |
| + | 3245741 |

| | |
|---|---|
| + | 3246447 |
| + | 3246527 |
| + | 3246528 |
| - | 3246612 |
| - | 3248732 |
| + | 3248918 |
| + | 3251921 |
| - | 3253203 |
| + | 3253218 |
| + | 3253302 |
| - | 3253473 |
| - | 3253475 |
| + | 3254562 |
| - | 3257040 |
| - | 3257305 |
| - | 3257974 |
| + | 3258645 |
| - | 3259299 |
| - | 3259301 |
| - | 3261115 |
| + | 3261423 |
| - | 3261466 |
| + | 3261840 |
| - | 3262282 |
| - | 3262353 |
| - | 3263770 |
| + | 3263825 |
| + | 3264456 |
| - | 3265233 |
| - | 3265234 |
| - | 3265297 |
| - | 3266153 |
| - | 3266154 |
| + | 3266169 |
| + | 3266641 |
| - | 3267851 |
| + | 3268745 |
| + | 3268859 |
| + | 3272095 |
| - | 3273183 |
| + | 3273547 |
| + | 3273709 |
| - | 3276045 |
| + | 3276068 |
| - | 3276495 |
| + | 3276817 |
| + | 3276940 |
| + | 3276944 |
| - | 3277677 |
| - | 3277773 |

| | |
|---|---|
| + | 3278300 |
| + | 3279547 |
| - | 3280588 |
| + | 3281204 |
| + | 3282208 |
| + | 3291371 |
| - | 3292053 |
| - | 3292404 |
| - | 3292406 |
| + | 3292923 |
| - | 3293427 |
| - | 3293429 |
| + | 3294240 |
| + | 3294242 |
| - | 3294338 |
| - | 3295218 |
| - | 3296326 |
| - | 3297448 |
| + | 3297974 |
| - | 3297991 |
| - | 3299685 |
| - | 3299688 |
| - | 3299889 |
| - | 3299892 |
| - | 3300939 |
| - | 3301443 |
| - | 3301444 |
| + | 3301549 |
| + | 3301556 |
| - | 3302876 |
| - | 3302998 |
| - | 3304381 |
| - | 3305419 |
| - | 3305711 |
| - | 3305993 |
| - | 3306104 |
| - | 3306928 |
| - | 3307590 |
| + | 3307602 |
| - | 3308272 |
| - | 3308273 |
| - | 3308621 |
| - | 3308623 |
| + | 3308799 |
| - | 3309391 |
| - | 3310321 |
| - | 3310322 |
| + | 3310399 |
| + | 3310400 |
| + | 3310404 |

| | |
|---|---|
| + | 3310461 |
| + | 3312120 |
| + | 3312122 |
| - | 3312754 |
| - | 3312755 |
| - | 3312875 |
| - | 3314786 |
| - | 3315925 |
| - | 3316200 |
| - | 3316816 |
| - | 3319031 |
| - | 3319247 |
| - | 3319583 |
| + | 3319841 |
| - | 3319932 |
| - | 3319935 |
| - | 3321344 |
| + | 3321572 |
| + | 3321578 |
| - | 3322072 |
| + | 3324652 |
| - | 3325684 |
| + | 3325695 |
| - | 3326778 |
| - | 3326780 |
| + | 3326968 |
| - | 3327166 |
| - | 3327167 |
| + | 3328681 |
| - | 3330460 |
| + | 3334747 |
| - | 3341004 |
| - | 3343697 |
| + | 3344089 |
| + | 3346192 |
| + | 3346271 |
| + | 3346274 |
| + | 3346712 |
| - | 3346722 |
| - | 3347240 |
| - | 3347440 |
| - | 3349889 |
| + | 3351070 |
| - | 3352145 |
| - | 3352721 |
| - | 3353367 |
| - | 3353934 |
| - | 3353935 |
| - | 3354968 |
| - | 3355040 |

| | |
|---|---|
| - | 3355323 |
| - | 3357133 |
| - | 3357509 |
| - | 3360826 |
| - | 3360883 |
| - | 3360884 |
| + | 3361132 |
| - | 3361182 |
| - | 3361183 |
| - | 3361184 |
| + | 3361633 |
| - | 3361695 |
| + | 3363050 |
| - | 3363729 |
| + | 3364272 |
| - | 3364534 |
| - | 3364535 |
| - | 3364536 |
| - | 3364537 |
| - | 3364989 |
| - | 3365381 |
| + | 3365995 |
| - | 3366535 |
| - | 3366538 |
| - | 3366954 |
| - | 3372464 |
| + | 3372525 |
| + | 3373718 |
| + | 3374432 |
| + | 3376797 |
| + | 3376799 |
| - | 3377419 |
| + | 3377838 |
| + | 3377839 |
| - | 3378626 |
| + | 3378787 |
| - | 3379212 |
| - | 3380065 |
| - | 3380067 |
| - | 3380069 |
| + | 3382088 |
| + | 3382172 |
| - | 3382525 |
| - | 3382527 |
| + | 3382610 |
| + | 3383535 |
| + | 3383538 |
| - | 3384013 |
| + | 3385107 |
| - | 3385606 |

| | |
|---|---|
| - | 3385683 |
| + | 3385695 |
| - | 3386898 |
| + | 3387302 |
| - | 3387997 |
| - | 3387999 |
| + | 3388007 |
| - | 3388289 |
| - | 3388825 |
| - | 3388827 |
| - | 3390440 |
| - | 3390443 |
| - | 3390710 |
| - | 3390712 |
| - | 3391992 |
| - | 3393254 |
| + | 3394503 |
| - | 3395205 |
| + | 3395866 |
| - | 3396546 |
| - | 3397765 |
| - | 3397766 |
| - | 3398954 |
| - | 3400399 |
| - | 3400400 |
| + | 3402514 |
| - | 3404688 |
| + | 3404809 |
| - | 3406570 |
| - | 3409102 |
| - | 3409458 |
| - | 3409997 |
| + | 3410395 |
| + | 3411116 |
| + | 3413053 |
| - | 3415263 |
| - | 3415264 |
| - | 3416570 |
| - | 3416888 |
| + | 3418067 |
| - | 3418311 |
| - | 3418312 |
| - | 3418314 |
| + | 3418423 |
| + | 3418426 |
| - | 3421380 |
| - | 3421383 |
| + | 3421733 |
| - | 3422142 |
| + | 3422306 |

| | |
|---|---|
| - | 3423161 |
| - | 3424361 |
| - | 3424372 |
| - | 3425175 |
| - | 3425177 |
| + | 3425247 |
| + | 3425249 |
| - | 3427611 |
| + | 3427771 |
| - | 3430308 |
| + | 3430371 |
| - | 3430473 |
| - | 3430478 |
| - | 3434200 |
| - | 3434473 |
| + | 3434593 |
| - | 3436523 |
| - | 3436749 |
| - | 3437230 |
| - | 3437498 |
| - | 3437501 |
| - | 3441052 |
| + | 3442368 |
| - | 3443848 |
| - | 3444226 |
| - | 3444230 |
| + | 3444328 |
| + | 3444331 |
| + | 3444715 |
| - | 3444986 |
| - | 3446156 |
| - | 3446157 |
| - | 3446920 |
| - | 3448565 |
| - | 3448858 |
| - | 3448973 |
| - | 3449661 |
| - | 3450594 |
| - | 3451200 |
| - | 3451766 |
| - | 3452108 |
| - | 3454640 |
| - | 3454775 |
| - | 3455354 |
| - | 3455363 |
| + | 3455530 |
| - | 3456552 |
| - | 3456554 |
| + | 3456641 |
| + | 3456642 |

| | |
|---|---|
| - | 3457537 |
| + | 3457595 |
| + | 3457599 |
| + | 3458032 |
| - | 3458152 |
| + | 3458305 |
| + | 3459056 |
| + | 3460215 |
| + | 3461624 |
| + | 3463506 |
| - | 3465136 |
| + | 3465657 |
| + | 3465778 |
| - | 3466382 |
| - | 3466517 |
| - | 3466753 |
| - | 3466792 |
| + | 3466918 |
| - | 3467384 |
| + | 3467585 |
| - | 3471046 |
| - | 3471049 |
| + | 3471097 |
| + | 3471236 |
| - | 3472272 |
| - | 3472520 |
| + | 3472597 |
| - | 3473290 |
| + | 3473326 |
| + | 3473327 |
| + | 3473330 |
| - | 3473699 |
| + | 3474217 |
| + | 3475582 |
| - | 3475964 |
| - | 3475968 |
| + | 3475987 |
| - | 3482174 |
| + | 3482602 |
| - | 3482769 |
| - | 3483872 |
| - | 3483873 |
| - | 3483984 |
| - | 3484770 |
| - | 3484772 |
| - | 3485248 |
| + | 3485642 |
| + | 3485643 |
| - | 3487887 |
| - | 3488900 |

| | |
|---|---|
| + | 3488923 |
| + | 3488924 |
| + | 3489952 |
| - | 3490057 |
| - | 3490761 |
| - | 3491593 |
| - | 3491595 |
| - | 3491597 |
| + | 3492555 |
| - | 3492714 |
| - | 3495767 |
| + | 3495768 |
| - | 3495769 |
| - | 3497366 |
| - | 3497642 |
| - | 3497644 |
| + | 3497652 |
| + | 3498482 |
| + | 3498568 |
| + | 3499501 |
| + | 3504707 |
| + | 3505510 |
| - | 3508556 |
| - | 3508557 |
| + | 3509644 |
| - | 3509690 |
| + | 3510687 |
| - | 3513112 |
| + | 3513645 |
| - | 3513823 |
| - | 3513824 |
| - | 3513825 |
| - | 3514946 |
| + | 3522693 |
| - | 3524653 |
| - | 3528240 |
| - | 3529911 |
| - | 3529912 |
| + | 3530066 |
| + | 3530067 |
| - | 3535486 |
| + | 3535813 |
| - | 3536845 |
| - | 3537911 |
| + | 3539953 |
| + | 3540792 |
| - | 3541854 |
| + | 3544620 |
| - | 3544689 |
| - | 3544994 |

| | |
|---|---|
| - | 3545605 |
| - | 3545994 |
| + | 3546201 |
| - | 3546885 |
| - | 3547793 |
| - | 3548071 |
| + | 3550199 |
| - | 3557711 |
| - | 3558749 |
| - | 3559563 |
| - | 3559565 |
| + | 3559599 |
| - | 3561993 |
| - | 3562225 |
| + | 3562352 |
| + | 3562541 |
| + | 3562542 |
| + | 3562815 |
| - | 3566659 |
| - | 3568183 |
| + | 3568246 |
| + | 3568251 |
| - | 3569066 |
| - | 3570196 |
| - | 3570710 |
| - | 3570744 |
| - | 3572975 |
| - | 3573116 |
| - | 3573117 |
| - | 3574235 |
| + | 3575313 |
| - | 3575971 |
| - | 3575972 |
| + | 3576008 |
| - | 3576010 |
| - | 3576272 |
| - | 3576914 |
| - | 3577166 |
| - | 3577648 |
| - | 3579580 |
| - | 3579582 |
| + | 3580518 |
| - | 3581440 |
| - | 3581896 |
| - | 3584973 |
| + | 3585047 |
| + | 3585108 |
| + | 3586589 |
| - | 3589424 |
| + | 3589586 |

| | |
|---|---|
| + | 3589587 |
| + | 3590507 |
| - | 3591533 |
| - | 3595218 |
| + | 3595328 |
| - | 3598153 |
| + | 3598169 |
| + | 3598784 |
| - | 3599304 |
| + | 3600721 |
| + | 3601889 |
| + | 3602036 |
| - | 3604743 |
| + | 3604925 |
| - | 3606126 |
| + | 3608953 |
| - | 3609008 |
| - | 3609326 |
| - | 3609977 |
| - | 3610252 |
| - | 3614579 |
| - | 3614966 |
| - | 3615375 |
| - | 3615429 |
| - | 3617032 |
| - | 3617499 |
| + | 3618337 |
| - | 3618631 |
| - | 3621569 |
| - | 3622120 |
| - | 3622207 |
| - | 3622209 |
| + | 3622320 |
| - | 3622705 |
| - | 3625546 |
| - | 3625632 |
| - | 3625659 |
| - | 3626108 |
| - | 3626110 |
| - | 3626466 |
| - | 3626917 |
| - | 3627004 |
| - | 3627864 |
| + | 3630209 |
| - | 3630908 |
| - | 3630910 |
| + | 3631573 |
| - | 3631608 |
| + | 3632132 |
| + | 3632481 |

| | |
|---|---|
| - | 3632633 |
| - | 3632906 |
| - | 3633449 |
| - | 3633954 |
| - | 3634783 |
| - | 3634888 |
| - | 3635995 |
| - | 3636299 |
| - | 3636748 |
| + | 3638348 |
| - | 3638520 |
| + | 3639801 |
| + | 3640344 |
| - | 3640592 |
| + | 3640658 |
| - | 3641093 |
| + | 3641239 |
| - | 3642135 |
| - | 3642137 |
| - | 3642224 |
| - | 3642859 |
| - | 3643154 |
| - | 3643283 |
| - | 3643575 |
| - | 3643576 |
| - | 3643615 |
| + | 3643627 |
| + | 3643964 |
| - | 3644537 |
| - | 3644539 |
| - | 3645803 |
| - | 3645928 |
| - | 3646339 |
| - | 3646567 |
| + | 3646575 |
| + | 3646620 |
| - | 3646659 |
| - | 3646662 |
| + | 3646718 |
| - | 3647699 |
| - | 3647701 |
| + | 3648405 |
| - | 3648977 |
| - | 3649379 |
| - | 3649783 |
| - | 3649802 |
| - | 3649805 |
| - | 3649806 |
| - | 3650147 |
| + | 3653463 |

| | |
|---|---|
| - | 3653547 |
| - | 3654663 |
| - | 3658936 |
| - | 3660342 |
| + | 3660691 |
| - | 3663109 |
| - | 3663130 |
| + | 3663197 |
| + | 3663257 |
| + | 3665417 |
| - | 3665422 |
| + | 3665547 |
| + | 3665549 |
| - | 3665784 |
| - | 3667832 |
| + | 3667844 |
| - | 3668193 |
| + | 3669968 |
| - | 3672493 |
| - | 3672496 |
| - | 3672497 |
| - | 3672721 |
| - | 3672723 |
| + | 3672795 |
| - | 3675531 |
| - | 3676056 |
| + | 3677854 |
| + | 3677957 |
| + | 3679014 |
| + | 3679198 |
| - | 3680555 |
| - | 3681090 |
| - | 3681092 |
| - | 3681096 |
| + | 3681231 |
| + | 3681296 |
| + | 3681721 |
| - | 3682704 |
| - | 3683147 |
| + | 3684061 |
| + | 3684562 |
| + | 3685745 |
| + | 3685978 |
| - | 3687506 |
| - | 3688575 |
| - | 3688695 |
| - | 3688699 |
| + | 3690053 |
| + | 3691887 |
| - | 3693781 |

| | |
|---|---|
| + | 3694090 |
| + | 3694091 |
| + | 3694213 |
| + | 3695284 |
| + | 3695286 |
| + | 3695288 |
| - | 3696784 |
| - | 3697523 |
| - | 3697822 |
| + | 3698636 |
| + | 3699151 |
| + | 3699599 |
| - | 3700664 |
| - | 3701203 |
| + | 3701379 |
| - | 3702963 |
| + | 3707106 |
| + | 3707962 |
| - | 3708724 |
| - | 3708725 |
| - | 3708727 |
| - | 3710167 |
| - | 3711347 |
| - | 3711394 |
| + | 3711457 |
| + | 3712510 |
| + | 3717972 |
| + | 3718512 |
| + | 3718777 |
| + | 3718778 |
| + | 3720896 |
| + | 3723385 |
| + | 3723659 |
| - | 3724089 |
| + | 3724644 |
| - | 3724941 |
| - | 3725182 |
| - | 3725595 |
| + | 3726199 |
| - | 3726583 |
| - | 3726588 |
| - | 3726841 |
| - | 3727269 |
| - | 3727422 |
| - | 3728197 |
| - | 3728199 |
| - | 3729407 |
| - | 3730862 |
| - | 3730956 |
| + | 3732873 |

| | |
|---|---|
| - | 3733304 |
| - | 3733724 |
| + | 3733808 |
| + | 3733810 |
| - | 3733958 |
| + | 3735494 |
| - | 3736952 |
| - | 3740571 |
| - | 3740921 |
| - | 3741033 |
| + | 3741054 |
| + | 3741147 |
| + | 3741148 |
| + | 3742635 |
| + | 3743435 |
| - | 3743653 |
| + | 3744197 |
| - | 3744222 |
| + | 3744305 |
| - | 3744315 |
| + | 3744660 |
| + | 3747160 |
| - | 3747171 |
| - | 3748310 |
| - | 3748322 |
| - | 3748324 |
| + | 3748804 |
| + | 3749009 |
| + | 3749010 |
| + | 3749600 |
| + | 3749984 |
| + | 3750536 |
| + | 3751698 |
| - | 3752208 |
| + | 3753013 |
| - | 3755804 |
| - | 3755893 |
| - | 3756541 |
| + | 3756743 |
| + | 3756745 |
| - | 3757623 |
| + | 3757906 |
| - | 3758036 |
| - | 3758505 |
| + | 3758522 |
| + | 3758524 |
| - | 3758648 |
| - | 3759627 |
| - | 3760207 |
| + | 3760225 |

| | |
|---|---|
| + | 3760227 |
| - | 3760567 |
| - | 3762472 |
| - | 3762474 |
| - | 3762476 |
| + | 3762574 |
| + | 3763118 |
| + | 3763119 |
| + | 3763120 |
| - | 3764980 |
| + | 3765015 |
| + | 3765017 |
| - | 3766087 |
| - | 3766629 |
| - | 3766630 |
| - | 3768100 |
| - | 3768970 |
| - | 3769377 |
| - | 3769947 |
| - | 3770321 |
| - | 3770323 |
| + | 3770536 |
| + | 3770785 |
| - | 3771449 |
| + | 3772077 |
| - | 3772162 |
| - | 3772233 |
| - | 3773390 |
| - | 3773923 |
| - | 3773924 |
| - | 3774182 |
| - | 3774583 |
| - | 3774882 |
| - | 3775294 |
| + | 3776806 |
| + | 3777579 |
| + | 3777686 |
| + | 3777891 |
| - | 3779660 |
| - | 3779789 |
| + | 3780176 |
| - | 3780423 |
| - | 3780424 |
| + | 3780545 |
| - | 3780646 |
| - | 3780650 |
| - | 3780756 |
| - | 3783828 |
| - | 3784715 |
| - | 3785569 |

| | |
|---|---|
| + | 3786881 |
| - | 3788252 |
| - | 3788253 |
| - | 3789103 |
| - | 3789105 |
| - | 3789106 |
| - | 3790481 |
| - | 3791744 |
| - | 3791746 |
| - | 3791747 |
| - | 3791946 |
| - | 3792279 |
| - | 3792280 |
| + | 3792341 |
| - | 3794066 |
| - | 3795116 |
| + | 3795463 |
| + | 3795910 |
| + | 3796999 |
| - | 3797627 |
| - | 3798185 |
| + | 3798269 |
| - | 3798654 |
| + | 3798721 |
| + | 3799404 |
| - | 3800048 |
| - | 3802244 |
| - | 3803357 |
| - | 3803359 |
| - | 3803364 |
| - | 3803365 |
| - | 3803368 |
| - | 3804884 |
| - | 3804968 |
| - | 3804970 |
| + | 3807511 |
| - | 3809480 |
| + | 3809536 |
| - | 3809965 |
| - | 3810055 |
| + | 3810065 |
| - | 3810612 |
| - | 3810615 |
| + | 3812076 |
| - | 3813147 |
| + | 3813196 |
| - | 3813778 |
| - | 3814344 |
| - | 3814345 |
| + | 3816147 |

| | |
|---|---|
| - | 3816561 |
| + | 3818441 |
| - | 3818510 |
| + | 3818844 |
| + | 3818873 |
| + | 3818875 |
| + | 3819231 |
| + | 3819234 |
| - | 3821517 |
| - | 3821519 |
| - | 3822428 |
| - | 3823099 |
| + | 3825396 |
| + | 3829195 |
| - | 3829977 |
| - | 3830641 |
| - | 3832256 |
| - | 3832258 |
| - | 3833545 |
| - | 3834342 |
| + | 3835630 |
| - | 3835815 |
| + | 3836013 |
| + | 3836331 |
| + | 3837083 |
| - | 3837727 |
| - | 3840593 |
| + | 3841955 |
| + | 3843808 |
| + | 3845965 |
| + | 3845967 |
| + | 3846908 |
| + | 3847009 |
| + | 3847043 |
| + | 3847319 |
| - | 3848074 |
| - | 3849702 |
| - | 3850123 |
| + | 3851612 |
| - | 3852116 |
| - | 3853034 |
| - | 3853415 |
| + | 3853436 |
| + | 3854192 |
| - | 3856976 |
| - | 3856977 |
| - | 3856978 |
| - | 3856983 |
| - | 3858918 |
| - | 3860862 |

| | |
|---|---|
| - | 3860864 |
| - | 3861298 |
| - | 3861301 |
| + | 3861409 |
| - | 3863274 |
| + | 3863799 |
| - | 3864295 |
| - | 3864296 |
| - | 3864824 |
| + | 3865030 |
| - | 3865259 |
| + | 3866336 |
| - | 3866404 |
| - | 3866527 |
| + | 3866556 |
| + | 3866565 |
| + | 3866856 |
| - | 3867350 |
| + | 3867455 |
| + | 3867457 |
| + | 3870574 |
| + | 3870576 |
| - | 3871345 |
| - | 3871347 |
| + | 3872282 |
| - | 3873520 |
| + | 3873774 |
| - | 3879505 |
| - | 3880551 |
| - | 3880553 |
| - | 3882101 |
| - | 3882751 |
| - | 3882752 |
| - | 3882888 |
| + | 3883085 |
| + | 3883088 |
| - | 3885658 |
| - | 3885868 |
| - | 3885970 |
| - | 3885972 |
| + | 3886922 |
| + | 3891499 |
| + | 3893497 |
| - | 3894433 |
| - | 3894442 |
| - | 3896153 |
| - | 3896156 |
| - | 3896157 |
| + | 3896262 |
| - | 3898678 |

| | |
|---|---|
| - | 3899209 |
| - | 3899211 |
| + | 3899225 |
| - | 3900629 |
| - | 3900844 |
| + | 3900939 |
| - | 3902154 |
| + | 3902164 |
| - | 3904175 |
| - | 3905174 |
| + | 3905306 |
| + | 3906001 |
| - | 3907002 |
| - | 3907005 |
| - | 3907443 |
| + | 3908492 |
| - | 3911315 |
| - | 3911656 |
| - | 3911860 |
| - | 3912244 |
| - | 3913548 |
| - | 3913549 |
| - | 3913640 |
| - | 3913642 |
| - | 3913644 |
| - | 3913903 |
| + | 3913915 |
| + | 3917095 |
| - | 3918319 |
| - | 3918321 |
| - | 3918322 |
| - | 3918324 |
| - | 3918509 |
| - | 3918614 |
| - | 3918616 |
| + | 3918745 |
| + | 3918746 |
| + | 3919983 |
| - | 3922270 |
| + | 3922273 |
| - | 3922450 |
| - | 3922987 |
| + | 3923138 |
| + | 3923141 |
| - | 3923827 |
| + | 3925750 |
| - | 3926089 |
| - | 3926418 |
| + | 3927707 |
| + | 3928050 |

| | |
|---|---|
| - | 3930073 |
| - | 3930588 |
| - | 3930590 |
| - | 3932839 |
| - | 3932840 |
| - | 3933147 |
| - | 3933180 |
| + | 3933190 |
| + | 3933192 |
| + | 3934334 |
| - | 3935544 |
| + | 3935554 |
| + | 3935571 |
| + | 3936573 |
| - | 3937042 |
| - | 3937043 |
| - | 3937044 |
| + | 3938229 |
| + | 3938231 |
| - | 3938241 |
| - | 3939464 |
| - | 3939693 |
| + | 3939837 |
| + | 3939840 |
| + | 3939841 |
| - | 3940991 |
| - | 3941882 |
| + | 3941944 |
| + | 3941947 |
| + | 3943778 |
| + | 3944219 |
| - | 3944703 |
| - | 3945445 |
| + | 3945671 |
| + | 3946354 |
| + | 3946355 |
| + | 3946453 |
| + | 3948361 |
| + | 3948525 |
| - | 3948693 |
| - | 3950622 |
| - | 3951734 |
| - | 3951795 |
| + | 3952016 |
| + | 3952017 |
| + | 3952071 |
| - | 3953258 |
| - | 3954411 |
| + | 3955231 |
| - | 3955783 |

| | |
|---|---|
| + | 3956363 |
| + | 3956448 |
| + | 3957332 |
| + | 3957335 |
| - | 3958853 |
| - | 3959521 |
| - | 3961921 |
| - | 3964003 |
| + | 3964026 |
| - | 3964900 |
| + | 3966383 |
| - | 3966439 |
| + | 3966725 |
| - | 3970878 |
| + | 3971008 |
| + | 3971015 |
| - | 3971872 |
| - | 3973300 |
| + | 3977334 |
| + | 3977349 |
| - | 3977768 |
| - | 3979390 |
| - | 3979392 |
| - | 3979516 |
| + | 3979711 |
| - | 3980845 |
| + | 3981198 |
| + | 3981199 |
| - | 3982484 |
| + | 3982947 |
| - | 3984056 |
| - | 3984057 |
| - | 3984058 |
| - | 3985265 |
| - | 3986273 |
| + | 3986400 |
| + | 3986793 |
| + | 3987301 |
| - | 3988970 |
| - | 3988971 |
| - | 3988972 |
| + | 3989093 |
| + | 3989094 |
| + | 3989192 |
| + | 3989194 |
| - | 3990994 |
| + | 3991017 |
| + | 3991635 |
| - | 3992183 |
| - | 3992422 |

| | |
|---|---|
| - | 3992552 |
| + | 3992654 |
| - | 3992688 |
| + | 3992698 |
| + | 3992970 |
| - | 3993099 |
| + | 3993307 |
| + | 3993864 |
| + | 3995056 |
| + | 3995057 |
| + | 3996388 |
| - | 3996731 |
| + | 3996742 |
| + | 3996744 |
| + | 3996890 |
| + | 3997194 |
| + | 3997750 |
| - | 3998477 |
| + | 3998486 |
| + | 3999141 |
| - | 4000348 |
| - | 4000847 |
| - | 4001608 |
| - | 4005023 |
| - | 4005313 |
| - | 4005435 |
| + | 4005521 |
| - | 4006039 |
| - | 4006060 |
| + | 4006914 |
| + | 4007443 |
| - | 4007712 |
| - | 4007714 |
| - | 4008120 |
| - | 4008124 |
| - | 4010235 |
| - | 4012603 |
| - | 4012789 |
| + | 4013194 |
| - | 4013734 |
| - | 4013736 |
| + | 4013740 |
| - | 4013899 |
| - | 4014524 |
| - | 4014526 |
| + | 4014657 |
| + | 4014659 |
| - | 4015416 |
| - | 4015419 |
| + | 4017211 |

| | |
|---|---|
| - | 4017451 |
| - | 4017819 |
| - | 4017921 |
| - | 4018857 |
| - | 4019149 |
| - | 4019606 |
| - | 4021856 |
| + | 4023411 |
| - | 4023641 |
| - | 4023939 |
| - | 4026185 |
| + | 4028375 |
| + | 4029068 |
| - | 4029891 |
| - | 4030583 |
| - | 4030630 |
| - | 4031151 |
| - | 4031646 |
| - | 4031732 |
| - | 4031746 |
| - | 4032282 |
| - | 4032284 |
| + | 4034822 |
| + | 4035606 |
| - | 4035816 |
| - | 4035820 |
| - | 4037182 |
| - | 4037578 |
| + | 4037603 |
| - | 4037656 |
| - | 4037662 |
| - | 4038755 |
| - | 4039213 |
| - | 4041059 |
| - | 4041060 |
| + | 4041450 |
| + | 4041451 |
| + | 4046189 |
| + | 4046484 |
| - | 4047158 |
| - | 4047862 |
| - | 4049440 |
| + | 4051174 |
| + | 4052259 |
| - | 4052304 |
| - | 4052306 |
| - | 4052308 |
| - | 4052309 |
| - | 4053367 |
| - | 4053369 |

| | |
|---|---|
| - | 4053370 |
| - | 4054312 |
| - | 4054313 |
| - | 4055494 |
| + | 4057701 |
| + | 4059890 |
| - | 4062223 |
| - | 4062226 |
| - | 4062343 |
| + | 4062720 |
| - | 4063586 |
| - | 4064530 |
| - | 4064532 |
| - | 4065019 |
| + | 4065036 |
| - | 4066066 |
| - | 4066067 |
| - | 4067021 |
| - | 4067023 |
| + | 4067130 |
| + | 4067131 |
| + | 4068613 |
| + | 4071151 |
| - | 4071253 |
| - | 4073002 |
| - | 4074717 |
| + | 4075145 |
| - | 4075235 |
| + | 4076339 |
| - | 4078164 |
| + | 4082009 |
| + | 4084425 |
| - | 4084574 |
| + | 4084727 |
| + | 4084770 |
| - | 4085275 |
| + | 4085409 |
| + | 4085571 |
| + | 4086376 |
| + | 4086655 |
| + | 4086657 |
| + | 4087263 |
| + | 4087377 |
| - | 4087377 |
| + | 4087944 |
| + | 4087945 |
| - | 4089266 |
| - | 4091403 |
| - | 4091405 |
| - | 4091752 |

| | |
|---|---|
| - | 4091753 |
| - | 4093696 |
| - | 4093927 |
| + | 4093945 |
| + | 4094454 |
| - | 4095509 |
| - | 4096934 |
| + | 4097007 |
| + | 4098571 |
| - | 4101040 |
| + | 4102182 |
| - | 4103696 |
| - | 4103697 |
| - | 4103698 |
| - | 4103699 |
| - | 4104217 |
| - | 4104355 |
| + | 4104415 |
| + | 4104416 |
| + | 4105234 |
| - | 4105871 |
| + | 4106173 |
| + | 4106553 |
| - | 4107960 |
| - | 4107963 |
| - | 4109580 |
| + | 4109682 |
| - | 4109732 |
| + | 4110255 |
| - | 4110576 |
| - | 4112021 |
| - | 4113262 |
| - | 4113378 |
| + | 4113379 |
| + | 4113381 |
| + | 4116547 |
| + | 4117052 |
| + | 4117054 |
| + | 4118600 |
| + | 4118601 |
| - | 4118778 |
| + | 4118884 |
| + | 4118886 |
| - | 4122693 |
| - | 4123062 |
| - | 4123068 |
| + | 4123160 |
| - | 4124846 |
| - | 4125741 |
| - | 4127425 |

| | |
|---|---|
| - | 4127672 |
| + | 4128019 |
| + | 4128020 |
| - | 4128145 |
| + | 4129632 |
| + | 4130036 |
| + | 4130161 |
| + | 4130178 |
| + | 4133458 |
| - | 4134636 |
| - | 4135211 |
| + | 4135326 |
| - | 4137049 |
| - | 4137662 |
| + | 4138825 |
| - | 4138960 |
| - | 4139581 |
| - | 4140206 |
| + | 4140218 |
| - | 4140559 |
| - | 4140698 |
| + | 4141118 |
| - | 4145553 |
| - | 4145554 |
| - | 4145555 |
| + | 4145713 |
| - | 4148254 |
| + | 4149253 |
| - | 4149678 |
| + | 4149723 |
| - | 4151861 |
| - | 4153283 |
| + | 4154005 |
| + | 4154662 |
| - | 4155052 |
| - | 4155293 |
| - | 4155294 |
| - | 4155296 |
| - | 4156769 |
| - | 4156774 |
| - | 4158868 |
| + | 4158981 |
| + | 4159443 |
| - | 4159469 |
| + | 4159555 |
| + | 4159556 |
| + | 4159558 |
| - | 4159744 |
| + | 4159763 |
| - | 4162211 |

| | |
|---|---|
| - | 4164125 |
| - | 4164127 |
| - | 4164156 |
| + | 4164576 |
| + | 4165155 |
| - | 4165334 |
| - | 4166709 |
| - | 4167006 |
| - | 4167008 |
| + | 4167081 |
| - | 4167885 |
| - | 4168055 |
| - | 4168163 |
| + | 4168166 |
| + | 4168168 |
| + | 4168170 |
| - | 4169936 |
| + | 4170016 |
| + | 4170266 |
| + | 4170809 |
| + | 4170981 |
| - | 4171793 |
| + | 4171810 |
| - | 4172546 |
| + | 4172820 |
| - | 4172847 |
| + | 4172925 |
| + | 4173086 |
| + | 4173719 |
| + | 4176205 |
| + | 4176890 |
| - | 4178170 |
| + | 4178250 |
| - | 4180476 |
| - | 4180478 |
| - | 4181271 |
| + | 4181618 |
| + | 4181957 |
| - | 4182640 |
| - | 4183178 |
| - | 4183952 |
| + | 4183957 |
| + | 4183959 |
| + | 4184180 |
| + | 4185767 |
| + | 4185855 |
| - | 4186493 |
| + | 4187265 |
| - | 4187616 |
| - | 4188655 |

| | |
|---|---|
| - | 4189181 |
| - | 4189183 |
| + | 4189380 |
| + | 4189778 |
| - | 4191158 |
| + | 4191168 |
| - | 4191169 |
| + | 4191170 |
| + | 4193312 |
| - | 4194077 |
| - | 4194196 |
| - | 4194198 |
| + | 4194342 |
| + | 4194883 |
| + | 4195318 |
| - | 4195459 |
| + | 4195535 |
| - | 4197184 |
| - | 4197185 |
| + | 4197437 |
| - | 4197767 |
| - | 4199798 |
| - | 4199801 |
| - | 4199803 |
| - | 4200995 |
| - | 4201186 |
| + | 4201877 |
| - | 4202282 |
| - | 4203358 |
| - | 4203360 |
| - | 4203361 |
| - | 4204279 |
| - | 4204541 |
| + | 4204736 |
| - | 4204869 |
| - | 4205439 |
| - | 4207194 |
| + | 4207375 |
| + | 4207597 |
| - | 4207819 |
| - | 4208783 |
| - | 4208785 |
| + | 4208847 |
| + | 4208942 |
| - | 4210676 |
| - | 4211815 |
| - | 4212331 |
| - | 4213014 |
| + | 4213213 |
| - | 4214655 |

| | |
|---|---|
| + | 4214786 |
| - | 4215493 |
| - | 4215496 |

**Table S2: Strains, plasmids, and oligonucleotides**

| Name | Description[1] | Source |
|---|---|---|
| | **Strains** | |
| *E. coli* JCB387 | Δ*nirB* Δ*lac* | (*1*) |
| *E. coli* T7 express | *fhuA2 lacZ*::T7 gene1 [lon] *ompT gal sulA11* R (*mcr73*::miniTn10--TetS)2 [*dcm*] R(*zgb*-210::Tn10--TetS) *endA1* Δ*mcrCmrr*)114::IS10 | Invitrogen |
| *B. subtilis* 168ca | *trpC*2 | (*2*) |
| | **Plasmids** | |
| pRW50 | Broad-host-range lac fusion vector for cloning promoters on *Eco*RI–*Hin*dIII fragments: contains the RK2 origin of replication and encodes tetracycline resistance. | (*3*) |
| pSR | pBR322-derived plasmid containing an *Eco*RI–*Hin*dIII cloning site upstream of the λoop transcription terminator. Encodes ampicillin resistance. | (*4*) |
| | **Oligonucleotides (5' to 3')** | |
| | *Amplification of $\sigma^{70}$ bound DNA flanked by* Eco*RI and* Hind*III sites for cloning in pRW50 or pSR* | |
| EWO0020 *wzxB* 1.1F | GGCTGCGAATTCacgttactttatctttactatctgc | This work |
| EWO0021 *wzxB* 1.1R | GCCCGAAGCTTCCTCCTttgtaagaacacttggtcctgaaaa | This work |
| EWO0022 *wzxB* 1.2F | GGCTGCGAATTCttgtaagaacacttggtcctgaaaa | This work |
| EWO0023 *wzxB* 1.2R | GCCCGAAGCTTCCTCCTacgttactttatctttactatctgc | This work |
| EWO0024 *yqiI* 1.1F | GGCTGCGAATTCcgcagtctgtagtggcggtcctgaac | This work |
| EWO0025 *yqiI* 1.1R | GCCCGAAGCTTCCTCCTatacctttaaattcaagtgctatatattc | This work |
| EWO0026 *yqiI* 1.2F | GGCTGCGAATTCatacctttaaattcaagtgctatatattc | This work |
| EWO0027 *yqiI* 1.2R | GCCCGAAGCTTTCCTCCTcgcagtctgtagtggcggtcctgaac | This work |
| EWO0028 *mcrB* 1.1F | GGCTGCGAATTCgtatgattcagtttttgacataggtg | This work |
| EWO0029 *mcrB* 1.1R | GCCCGAAGCTTCCTCCTcagtcctattacgcctgttcccaaaaag | This work |
| EWO0030 *mcrB* 1.2F | GGCTGCGAATTCcagtcctattacgcctgttcccaaaaag | This work |
| EWO0031 *mcrB* 1.2R | GCCCGAAGCTTCCTCCTgtatgattcagtttttgacataggtg | This work |
| EWO0032 *ygaQ* 1.1F | GGCTGCGAATTCcggttacacaatactaacttatttaac | This work |
| EWO0033 *ygaQ* 1.1R | GCCCGAAGCTTCCTCCTtgaaaaatcaatggcgcttaaatcatc | This work |
| EWO0034 *ygaQ* 1.2F | GGCTGCGAATTCtgaaaaatcaatggcgcttaaatcatc | This work |
| EWO0035 *ygaQ* 1.2R | GCCCGAAGCTTCCTCCTcggttacacaatactaacttatttaac | This work |
| EWO0036 *yagM* 1.1F | GGCTGCGAATTCagggatagccaaatccacaacttcc | This work |
| EWO0037 *yagM* 1.1R | GCCCGAAGCTTCCTCCTgagtcacaggtagtacagactactc | This work |
| EWO0038 *yeaI* 1.1F | GGCTGCGAATTCcgattagccagcgaactatggccg | This work |
| EWO0039 *yeaI* 1.1R | GCCCGAAGCTTCCTCCTcacaacaaacagtttactggaaacttc | This work |
| EWO0040 *yeaI* 1.2F | GGCTGCGAATTCcacaacaaacagtttactggaaacttc | This work |
| EWO0041 *yeaI* 1.2R | GCCCGAAGCTTCCTCCTcgattagccagcgaactatggccg | This work |
| EWO0042 *trkG* 1.1F | GGCTGCGAATTCttctttataactttcgttatattttttg | This work |
| EWO0043 *trkG* 1.1R | GCCCGAAGCTTCCTCCTtaaagggaatgcactaataacagaaaac | This work |
| EWO0044 *trkG* 1.2F | GGCTGCGAATTCtaaagggaatgcactaataacagaaaac | This work |
| EWO0045 *trkG* 1.2R | GCCCGAAGCTTCCTCCTttctttataactttcgttatattttttg | This work |
| EWO0046 *idnK* 1.1F | GGCTGCGAATTCggagtggtaaaacattaattggtag | This work |
| EWO0047 *idnK* 1.1R | GCCCGAAGCTTCCTCCTgttccagccagggaagtcgatcttc | This work |
| EWO0048 *idnK* 1.2F | GGCTGCGAATTCgttccagccagggaagtcgatcttc | This work |

| | | | |
|---|---|---|---|
| EWO0049 | *idnK* 1.2R | GCCCGAAGCTTCCTCCTggagtggtaaaacattaattggtag | This work |
| EWO0050 | *ycjW* 1.1F | GGCTGCGAATTCgaataaacatgggcatattgaccttc | This work |
| EWO0051 | *ycjW* 1.1R | GCCCGAAGCTTCCTCCTatgcgaaagcaaaattaagcagaaaatg | This work |
| EWO0052 | *ycjW* 1.2F | GGCTGCGAATTCatgcgaaagcaaaattaagcagaaaatg | This work |
| EWO0053 | *ycjW* 1.2R | GCCCGAAGCTTCCTCCTgaataaacatgggcatattgaccttc | This work |
| EWO0054 | *wcaD* 1.1F | GGCTGCGAATTCttcagatattgaaatgcccactccag | This work |
| EWO0055 | *wcaD* 1.1R | GCCCGAAGCTTCCTCCTgctcaatttggctcagcatcaaacag | This work |
| EWO0056 | *wcaD* 1.2F | GGCTGCGAATTCgctcaatttggctcagcatcaaacag | This work |
| EWO0057 | *wcaD* 1.2R | GCCCGAAGCTTCCTCCTttcagatattgaaatgcccactccag | This work |
| EWO0058 | *yeeL* 1.1F | GGCTGCGAATTCcttgccatatgtaatattgaggtg | This work |
| EWO0059 | *yeeL* 1.1R | GCCCGAAGCTTCCTCCTcaaaggtgaagataaagccagggc | This work |
| EWO0060 | *yeeL* 1.2F | GGCTGCGAATTCcaaaggtgaagataaagccagggc | This work |
| EWO0061 | *yeeL* 1.2R | GCCCGAAGCTTCCTCCTcttgccatatgtaatattgaggtg | This work |
| EWO0062 | *yigF* 1.1F | GGCTGCGAATTCtttctcatagaaccatttgttcgtg | This work |
| EWO0063 | *yigF* 1.1R | GCCCGAAGCTTCCTCCTtttgtcctggggttatggaaaaaag | This work |
| EWO0064 | *yigF* 1.2F | GGCTGCGAATTCtttgtcctggggttatggaaaaaag | This work |
| EWO0065 | *yigF* 1.2R | GCCCGAAGCTTCCTCCTtttctcatagaaccatttgttcgtg | This work |
| EWO0066 | *yigG* 1.1F | GGCTGCGAATTCcattgcctgaacaggcaaaatcttc | This work |
| EWO0067 | *yigG* 1.1R | GCCCGAAGCTTCCTCCTtactccattatctcgtcatcaacatg | This work |
| EWO0068 | *yigG* 1.2F | GGCTGCGAATTCtactccattatctcgtcatcaacatg | This work |
| EWO0069 | *yigG* 1.2R | GCCCGAAGCTTCCTCCTcattgcctgaacaggcaaaatcttc | This work |
| EWO0070 | *gadE* 1.1F | GGCTGCGAATTCtaaacccttcaaggttatcattgatac | This work |
| EWO0071 | *gadE* 1.1R | GCCCGAAGCTTCCTCCTaaaataaataggcgctttagcttttag | This work |
| EWO0072 | *gadE* 1.2F | GGCTGCGAATTCaaaataaataggcgctttagcttttag | This work |
| EWO0073 | *gadE* 1.2R | GCCCGAAGCTTCCTCCTtaaacccttcaaggttatcattgatac | This work |
| EWO0074 | *ybdO* 1.1F | GGCTGCGAATTCtcaacccctgctgatttagatattaattg | This work |
| EWO0075 | *ybdO* 1.1R | GCCCGAAGCTTCCTCCTcagccgttccgtcatttgcatgcc | This work |
| EWO0076 | *ybdO* 1.2F | GGCTGCGAATTCcagccgttccgtcatttgcatgcc | This work |
| EWO0077 | *ybdO* 1.2R | GCCCGAAGCTTCCTCCTtcaacccctgctgatttagatattaattg | This work |
| EWO0078 | *ybdO* 2.1F | GGCTGCGAATTCtcagcattctctgctgacatgagg | This work |
| EWO0079 | *ybdO* 2.1R | GCCCGAAGCTTCCTCCTcaataagtccgaactaaagaaaaac | This work |
| EWO0080 | *ybdO* 2.2F | GGCTGCGAATTCcaataagtccgaactaaagaaaaac | This work |
| EWO0081 | *ybdO* 2.2R | GCCCGAAGCTTCCTCCTtcagcattctctgctgacatgagg | This work |
| EWO0082 | *yehA* 1.1F | GGCTGCGAATTCtttaattttatggtatcgttataaag | This work |
| EWO0083 | *yehA* 1.1R | GCCCGAAGCTTCCTCCTttgttcaggggattttgcacttac | This work |
| EWO0084 | *yehA* 1.2F | GGCTGCGAATTCttgttcaggggattttgcacttac | This work |
| EWO0085 | *yehA* 1.2R | GCCCGAAGCTTCCTCCTtttaattttatggtatcgttataaag | This work |
| EWO0086 | *fepE* 1.1F | GGCTGCGAATTCgtggtgaaagagtcgctagaaaac | This work |
| EWO0087 | *fepE* 1.1R | GCCCGAAGCTTCCTCCTgttggcaatgtcgagtgaataattg | This work |
| EWO0088 | *fepE* 1.2F | GGCTGCGAATTCgttggcaatgtcgagtgaataattg | This work |
| EWO0089 | *fepE* 1.2R | GCCCGAAGCTTCCTCCTgtggtgaaagagtcgctagaaaac | This work |
| EWO0090 | *leuO* 1.1F | GGCTGCGAATTCtcagaacactgaacatcagctgcg | This work |
| EWO0091 | *leuO* 1.1R | GCCCGAAGCTTCCTCCTaattcccgggttgttggatgatttttg | This work |
| EWO0092 | *leuO* 1.2F | GGCTGCGAATTCaattcccgggttgttggatgatttttg | This work |
| EWO0093 | *leuO* 1.2R | GCCCGAAGCTTCCTCCTtcagaacactgaacatcagctgcg | This work |
| EWO0094 | *setC* 1.1F | GGCTGCGAATTCaagtatattcctcgcagatgaactg | This work |
| EWO0095 | *setC* 1.1R | GCCCGAAGCTTCCTCCTcatagcagaatcagtaatttacggtc | This work |
| EWO0096 | *setC* 1.2F | GGCTGCGAATTCcatagcagaatcagtaatttacggtc | This work |
| EWO0097 | *setC* 1.2R | GCCCGAAGCTTCCTCCTaagtatattcctcgcagatgaactg | This work |
| EWO0098 | *yagM* 2.1F | GGCTGCGAATTCcctcaaaggttgttctatgaataag | This work |
| EWO0099 | *yagM* 2.1R | GCCCGAAGCTTCCTCCTcagttatacgtgaaaggctatcctc | This work |
| EWO0100 | *yagM* 2.2F | GGCTGCGAATTCcagttatacgtgaaaggctatcctc | This work |
| EWO0101 | *yagM* 2.2R | GCCCGAAGCTTCCTCCTcctcaaaggttgttctatgaataag | This work |
| EWO0102 | *mcrC* 1.1F | GGCTGCGAATTCagtgtcctttgttttgacctggaatag | This work |
| EWO0103 | *mcrC* 1.1R | GCCCGAAGCTTCCTCCTgaaaattaccgggcattagcactcttc | This work |

| | | | |
|---|---|---|---|
| EWO0104 | *mcrC* 1.2F | GGCTGCGAATTCgaaaattaccgggcattagcactcttc | This work |
| EWO0105 | *mcrC* 1.2R | GCCCGAAGCTTCCTCCTagtgtcctttgttttgacctggaatag | This work |
| EWO0106 | *mcrC* 2.1F | GGCTGCGAATTCataaagtgaacgagcttcatctctg | This work |
| EWO0107 | *mcrC* 2.1R | GCCCGAAGCTTCCTCCTttccatcttaatcatgggaaaaccg | This work |
| EWO0108 | *mcrC* 2.2F | GGCTGCGAATTCttccatcttaatcatgggaaaaccg | This work |
| EWO0109 | *mcrC* 2.2R | GCCCGAAGCTTCCTCCTataaagtgaacgagcttcatctctg | This work |
| EWO0110 | *mcrB* 2.1F | GGCTGCGAATTCctccaagaaatgcaaaccagggaatag | This work |
| EWO0111 | *mcrB* 2.1R | GCCCGAAGCTTCCTCCTaaccctggattgaaaaatttattaag | This work |
| EWO0112 | *mcrB* 2.2F | GGCTGCGAATTCaaccctggattgaaaaatttattaag | This work |
| EWO0113 | *mcrB* 2.2R | GCCCGAAGCTTCCTCCTctccaagaaatgcaaaccagggaatag | This work |
| EWO0114 | *trkG* 2.1F | GGCTGCGAATTCtatcggatattttaataactatttg | This work |
| EWO0115 | *trkG* 2.1R | GCCCGAAGCTTCCTCCTaacagaaagaaacgaagttcaatatc | This work |
| EWO0116 | *trkG* 2.2F | GGCTGCGAATTCaacagaaagaaacgaagttcaatatc | This work |
| EWO0117 | *trkG* 2.2R | GCCCGAAGCTTCCTCCTtatcggatattttaataactatttg | This work |
| EWO0118 | *trkG* 3.1F | GGCTGCGAATTCaggtctgtatggagtttcttttttc | This work |
| EWO0119 | *trkG* 3.1R | GCCCGAAGCTTCCTCCTtgcagtagccccaaaacctaatccc | This work |
| EWO0120 | *trkG* 3.2F | GGCTGCGAATTCtgcagtagccccaaaacctaatccc | This work |
| EWO0121 | *trkG* 3.2R | GCCCGAAGCTTCCTCCTaggtctgtatggagtttcttttttc | This work |
| EWO0122 | *idnK* 2.1F | GGCTGCGAATTCgtctttataaaaagaatgaaacagg | This work |
| EWO0123 | *idnK* 2.1R | GCCCGAAGCTTCCTCCTcccgacgctgcattcgcgcgagaatag | This work |
| EWO0124 | *idnK* 2.2F | GGCTGCGAATTCcccgacgctgcattcgcgcgagaatag | This work |
| EWO0125 | *idnK* 2.2R | GCCCGAAGCTTCCTCCTgtctttataaaaagaatgaaacagg | This work |
| EWO0126 | *yqiI* 2.1F | GGCTGCGAATTCgaatattttatgaatgttttctg | This work |
| EWO0127 | *yqiI* 2.1R | GCCCGAAGCTTCCTCCTataagttacaccgaaagtataagag | This work |
| EWO0128 | *yqiI* 2.2F | GGCTGCGAATTCataagttacaccgaaagtataagag | This work |
| EWO0129 | *yqiI* 2.2R | GCCCGAAGCTTCCTCCTgaatattttatgaatgttttctg | This work |
| EWO0130 | *ygaQ* 2.1F | GGCTGCGAATTCttaaagatccagtaacaaaagaacg | This work |
| EWO0131 | *ygaQ* 2.1R | GCCCGAAGCTTCCTCCTgcattccatttaaacgcttttc | This work |
| EWO0132 | *ygaQ* 2.2F | GGCTGCGAATTCgcattccatttaaacgcttttc | This work |
| EWO0133 | *ygaQ* 2.2R | GCCCGAAGCTTCCTCCTttaaagatccagtaacaaaagaacg | This work |
| EWO0134 | *evgS* 1.1F | GGCTGCGAATTCcctaatgaactttatcattttcttattc | This work |
| EWO0135 | *evgS* 1.1R | GCCCGAAGCTTCCTCCTtttttccgtcaggcgcagaacttcg | This work |
| EWO0136 | *evgS* 1.2F | GGCTGCGAATTCtttttccgtcaggcgcagaacttcg | This work |
| EWO0137 | *evgS* 1.2R | GCCCGAAGCTTCCTCCTcctaatgaactttatcattttcttattc | This work |
| EWO0138 | *evgS* 2.1F | GGCTGCGAATTCaaagcactctcggattccttaccg | This work |
| EWO0139 | *evgS* 2.1R | GCCCGAAGCTTCCTCCTaaagggtgagtcactgttttctaatg | This work |
| EWO0140 | *evgS* 2.2F | GGCTGCGAATTCaaagggtgagtcactgttttctaatg | This work |
| EWO0141 | *evgS* 2.2R | GCCCGAAGCTTCCTCCTaaagcactctcggattccttaccg | This work |
| EWO0142 | *evgS* 3.1F | GGCTGCGAATTCatatacacacaggtatttgaaattg | This work |
| EWO0143 | *evgS* 3.1R | GCCCGAAGCTTCCTCCTtgcattaattagatcacgcgtttcag | This work |
| EWO0144 | *evgS* 3.2F | GGCTGCGAATTCtgcattaattagatcacgcgtttcag | This work |
| EWO0145 | *evgS* 3.2R | GCCCGAAGCTTCCTCCTatatacacacaggtatttgaaattg | This work |
| EWO0146 | *yibA* 1.1F | GGCTGCGAATTCgatatcggagcatttatactcgggc | This work |
| EWO0147 | *yibA* 1.1R | GCCCGAAGCTTCCTCCTtttcttgcatcgctgagccgttgac | This work |
| EWO0148 | *yibA* 1.2F | GGCTGCGAATTCtttcttgcatcgctgagccgttgac | This work |
| EWO0149 | *yibA* 1.2R | GCCCGAAGCTTCCTCCTgatatcggagcatttatactcgggc | This work |
| EWO0150 | *yibA* 2.1F | GGCTGCGAATTCgggttttatctgttttatgcgatgag | This work |
| EWO0151 | *yibA* 2.1R | GCCCGAAGCTTCCTCCTcggaagttataatttcattgtcatc | This work |
| EWO0152 | *yibA* 2.2F | GGCTGCGAATTCcggaagttataatttcattgtcatc | This work |
| EWO0153 | *yibA* 2.2R | GCCCGAAGCTTCCTCCTgggttttatctgttttatgcgatgag | This work |
| EWO0154 | *elaD* 1.1F | GGCTGCGAATTCgccgaatgaagtccagttattcccc | This work |
| EWO0155 | *elaD* 1.1R | GCCCGAAGCTTCCTCCTttttctttatcatagcctagtgcac | This work |
| EWO0156 | *elaD* 1.2F | GGCTGCGAATTCttttctttatcatagcctagtgcac | This work |
| EWO0157 | *elaD* 1.2R | GCCCGAAGCTTCCTCCTgccgaatgaagtccagttattcccc | This work |
| EWO0158 | *sfmD* 1.1F | GGCTGCGAATTCcggcaatacaggaagtgatattttc | This work |

| | | |
|---|---|---|
| EWO0159 *sfmD* 1.1R | GCCCGAAGCTTCCTCCTtgccgaatcgttagctgggccgccg | This work |
| EWO0160 *sfmD* 1.2F | GGCTGCGAATTCtgccgaatcgttagctgggccgccg | This work |
| EWO0161 *sfmD* 1.2R | GCCCGAAGCTTCCTCCTcggcaatacaggaagtgatattttc | This work |

*Mutation of shared $\sigma^{70}$ promoter elements*

| | | |
|---|---|---|
| EWO0162 *wzxB*1.1R | GCCCGAAGCTTcctcctttgtaagaacacttggtcctgaaaatttcg<br>gtatattcggtttttgccaagcgactatgctatacccgataatgtttgttg | This work |
| EWO0163 *wzxB*1.1R | GCCCGAAGCTTcctcctttgtaagaacacttggtcctgaaaatttcg<br>gtatattcggtttttgccaagcgactatcgtatatatgataatgtttgttg | This work |
| EWO0164 *wzxB*1.2F | GGCTGCGAATTCttgtaagaacacttggtcctgaaaatttcggtata<br>ttcggtttttgccaagcgactatgctatacccgataatgtttgttgaatatgg | This work |
| EWO0165 *wzxB*1.2F | GGCTGCGAATTCttgtaagaacacttggtcctgaaaatttcggtata<br>ttcggtttttgccaagcgactatcgtatatatgataatgtttgttgaatatgg | This work |
| EWO0166 *yigG*1.1 | GGCTGCGAATTCcattgcctgaacaggcaaaatcttcggctatcat<br>tgtgatgatagagatgatatatactgctcctgtaccaaaaacataag | This work |
| EWO0167 *yigG*1.2R | GCCCGAAGCTTcctcctcattgcctgaacaggcaaaatcttcggcta<br>tcattgtgatgatagagatgatatatactgctcctgtaccaaaaacataag | This work |
| EWO0168 *yqiI*2.1R | GCCCGAAGCTTcctcctataagttacaccgaaagtataagagttttga<br>ttataaaa-Δ-acctgatgctaacaacatcattatatttgc | This work |
| EWO0169 *yqiI*2.2F | GGCTGCGAATTCataagttacaccgaaagtataagagttttgattata<br>aaa-Δ-acctgatgctaacaacatcattatatttgcccatgc | This work |
| EWO0170 *ygaQ*1.1F | GGCTGCGAATTC-Δ-acttatttaacccaaaatatcataaaaaagcc<br>gttaccaattacatggaatatctggtaac | This work |
| EWO0171 *ygaQ*1.2R | GCCCGAAGCTTcctcct-Δ-acttatttaacccaaaatatcataaaaaa<br>gccgttaccaattacatggaatatctggtaacttgtc | This work |

*Amplification of canonical promoters flanked by* Eco*RI and* Hind*III sites*
*for cloning in pRW50 or pSR*

| | | |
|---|---|---|
| EWO0177 *fixA* 1.1F | GGCTGCGAATTCtgaggaacttaacaatattg | This work |
| EWO0178 *fixA* 1.1R | GCCCGAAGCTTCCTCCTatctccagaaatcatgaagg | This work |
| EWO0179 *fixA* 1.2F | GGCTGCGAATTCatctccagaaatcatgaagg | This work |
| EWO0180 *fixA* 1.2R | GCCCGAAGCTTCCTCCTtgaggaacttaacaatattg | This work |
| EWO0181 *araC* 1.1F | GGCTGCGAATTCcgccgtgcaaataatcaatg | This work |
| EWO0182 *araC* 1.1R | GCCCGAAGCTTCCTCCTtcttttactggctcttctcg | This work |
| EWO0183 *araC* 1.2F | GGCTGCGAATTCtcttttactggctcttctcg | This work |
| EWO0184 *araC* 1.2R | GCCCGAAGCTTCCTCCTcgccgtgcaaataatcaatg | This work |
| EWO0185 *ilvIH* 1.1F | GGCTGCGAATTCtattcttattaccccgtgtttatg | This work |
| EWO0186 *ilvIH* 1.1R | GCCCGAAGCTTCCTCCTgataagcgatcggacgaccatc | This work |
| EWO0187 *ilvIH* 1.2F | GGCTGCGAATTCgataagcgatcggacgaccatc | This work |
| EWO0188 *ilvIH* 1.2R | GCCCGAAGCTTCCTCCTtattcttattaccccgtgtttatg | This work |
| EWO0189 *apt* 1.1F | GGCTGCGAATTCtgatgaaaagcaagaaaagc | This work |
| EWO0190 *apt* 1.1R | GCCCGAAGCTTCCTCCTactcacggcgcgttttaaacg | This work |
| EWO0191 *apt* 1.2F | GGCTGCGAATTCactcacggcgcgttttaaacg | This work |
| EWO0192 *apt* 1.2R | GCCCGAAGCTTCCTCCTtgatgaaaagcaagaaaagc | This work |
| EWO0193 *cstA* 3.1F | GGCTGCGAATTCactccgatttacatggttgc | This work |
| EWO0194 *cstA* 3.1R | GCCCGAAGCTTCCTCCTtgctcccattacagagagcac | This work |
| EWO0195 *cstA* 3.2F | GGCTGCGAATTCtgctcccattacagagagcac | This work |
| EWO0196 *cstA* 3.2R | GCCCGAAGCTTCCTCCTactccgatttacatggttgc | This work |
| EWO0197 *asnB1*.1F | GGCTGCGAATTCtgatatcgaatacgccaaaaattg | This work |
| EWO0198 *asnB1*.1R | GCCCGAAGCTTCCTCCTtcaccattacgtttttatttttc | This work |
| EWO0199 *asnB1*.2F | GGCTGCGAATTCtcaccattacgtttttatttttc | This work |
| EWO0200 *asnB1*.2R | GCCCGAAGCTTCCTCCTtgatatcgaatacgccaaaaattg | This work |

| | | |
|---|---|---|
| EWO0201 *gltA1*.1F | GGCTGCGAATTCaataactgtcccgaatgaattg | This work |
| EWO0202 *gltA1*.11R | GCCCGAAGCTTCCTCCTtcatctaatttgacaatcattc | This work |
| EWO0203 *gltA1*.12F | GGCTGCGAATTCtcatctaatttgacaatcattc | This work |
| EWO0204 *gltA1*.12R | GCCCGAAGCTTCCTCCTaataactgtcccgaatgaattg | This work |
| EWO0205 *ompA2*.1F | GGCTGCGAATTCgttaaatccttcaccggggg | This work |
| EWO0206 *ompA2*.1R | GCCCGAAGCTTCCTCCTatacaagactttttttcatatg | This work |
| EWO0207 *ompA2*.2F | GGCTGCGAATTCatacaagactttttttcatatg | This work |
| EWO0208 *ompA2*.2R | GCCCGAAGCTTCCTCCTgttaaatccttcaccggggg | This work |
| EWO0209 *tdK* 8.1F | GGCTGCGAATTCaagggagaacgcatataccc | This work |
| EWO0210 *tdK* 8.1R | GCCCGAAGCTTCCTCCTcccgcattcattgcggaatag | This work |
| EWO0211 *tdK* 8.2F | GGCTGCGAATTCcccgcattcattgcggaatag | This work |
| EWO0212 *tdK* 8.2R | GCCCGAAGCTTCCTCCTaagggagaacgcatataccc | This work |
| EWO0213 *osmB* 1.1F | GGCTGCGAATTCacagccgcggtcattttttg | This work |
| EWO0214 *osmB* 1.1R | GCCCGAAGCTTCCTCCTcgtgatataaccctgcgcgcgag | This work |
| EWO0215 *osmB* 1.2F | GGCTGCGAATTCcgtgatataaccctgcgcgcgag | This work |
| EWO0216 *osmB* 1.2R | GCCCGAAGCTTCCTCCTacagccgcggtcattttttg | This work |
| EWO0217 *hisB* 1.1F | GGCTGCGAATTCaaccaactacattctggcgc | This work |
| EWO0218 *hisB* 1.1R | GCCCGAAGCTTCCTCCTcgctggctttcttcacgggttc | This work |
| EWO0219 *hisB* 1.2F | GGCTGCGAATTCcgctggctttcttcacgggttc | This work |
| EWO0220 *hisB* 1.2R | GCCCGAAGCTTCCTCCTaaccaactacattctggcgc | This work |
| EWO0221 *cirA* 2.1F | GGCTGCGAATTCttttatgcaggtgatcatcc | This work |
| EWO0222 *cirA* 2.1R | GCCCGAAGCTTCCTCCTcaattccatttccctgacaaatc | This work |
| EWO0223 *cirA* 2.2F | GGCTGCGAATTCcaattccatttccctgacaaatc | This work |
| EWO0224 *cirA* 2.2R | GCCCGAAGCTTCCTCCTttttatgcaggtgatcatcc | This work |
| EWO0225 *bcp* 1.1F | GGCTGCGAATTCgcggacgcagcaaatattgag | This work |
| EWO0226 *bcp* 1.1R | GCCCGAAGCTTCCTCCTtcggtgcgatatcaccggctttc | This work |
| EWO0227 *bcp* 1.2F | GGCTGCGAATTCtcggtgcgatatcaccggctttc | This work |
| EWO0228 *bcp* 1.2R | GCCCGAAGCTTCCTCCTgcggacgcagcaaatattgag | This work |
| EWO0229 *pheL* 1.1F | GGCTGCGAATTCattgagtgtatcgccaacgc | This work |
| EWO0230 *pheL* 1.1R | GCCCGAAGCTTCCTCCTctcccattcaggggaaggtaaaaaag | This work |
| EWO0231 *pheL* 1.2F | GGCTGCGAATTCctcccattcaggggaaggtaaaaaag | This work |
| EWO0232 *pheL* 1.2R | GCCCGAAGCTTCCTCCTattgagtgtatcgccaacgc | This work |
| EWO0233 *cysJ* 1.1F | GGCTGCGAATTCgtgtcgtcatgcgtcgttatg | This work |
| EWO0234 *cysJ* 1.1R | GCCCGAAGCTTCCTCCTaggttagtcgatttggttattag | This work |
| EWO0235 *cysJ* 1.2F | GGCTGCGAATTCaggttagtcgatttggttattag | This work |
| EWO0236 *cysJ* 1.2R | GCCCGAAGCTTCCTCCTgtgtcgtcatgcgtcgttatg | This work |
| EWO0237 *agrR* 1.1F | GGCTGCGAATTCctttcataacattatttcag | This work |
| EWO0238 *agrR* 1.1R | GCCCGAAGCTTCCTCCTatgctttaaatgctttaactag | This work |
| EWO0239 *agrR* 1.2F | GGCTGCGAATTCatgctttaaatgctttaactag | This work |
| EWO0240 *agrR* 1.2R | GCCCGAAGCTTCCTCCTctttcataacattatttcag | This work |
| EWO0241 *rpsJ* 1.1F | GGCTGCGAATTCgattgggagcattgttaggtag | This work |
| EWO0242 *rpsJ* 1.1R | GCCCGAAGCTTCCTCCTgagagataacccgaaggctg | This work |
| EWO0243 *rpsJ* 1.2F | GGCTGCGAATTCgagagataacccgaaggctg | This work |
| EWO0244 *rpsJ* 1.2R | GCCCGAAGCTTCCTCCTgattgggagcattgttaggtag | This work |
| EWO0245 *ivbL* 1.1F | GGCTGCGAATTCgcagttggtagtagttttgc | This work |
| EWO0246 *ivbL* 1.1R | GCCCGAAGCTTCCTCCTaaacgtgatcaacccctcaattttcc | This work |
| EWO0247 *ivbL* 1.2F | GGCTGCGAATTCaaacgtgatcaacccctcaattttcc | This work |
| EWO0248 *ivbL* 1.2R | GCCCGAAGCTTCCTCCTgcagttggtagtagttttgc | This work |
| EWO0249 *dnaN* 3.1F | GGCTGCGAATTCtcgccagcgccatcgccatc | This work |
| EWO0250 *dnaN* 3.1R | GCCCGAAGCTTCCTCCTacttgctggcattgcaggaaaaac | This work |
| EWO0251 *dnaN* 3.2F | GGCTGCGAATTCacttgctggcattgcaggaaaaac | This work |
| EWO0252 *dnaN* 3.2R | GCCCGAAGCTTCCTCCTtcgccagcgccatcgccatc | This work |
| EWO0253 *trxA* 2.1F | GGCTGCGAATTCttaaatgtgttttgctcatag | This work |
| EWO0254 *trxA* 2.1R | GCCCGAAGCTTCCTCCTtatataactccacaggaataag | This work |
| EWO0255 *trxA* 2.2F | GGCTGCGAATTCtatataactccacaggaataag | This work |

| | | | |
|---|---|---|---|
| EWO0256 | *trxA* 2.2R | GCCCGAAGCTTCCTCCTttaaatgtgttttgctcatag | This work |
| EWO0257 | *rrsE* 2.1F | GGCTGCGAATTCacagccggttcggttgaagag | This work |
| EWO0258 | *rrsE* 2.1R | GCCCGAAGCTTCCTCCTttcgagtgcccacacagattg | This work |
| EWO0259 | *rrsE* 2.2F | GGCTGCGAATTCttcgagtgcccacacagattg | This work |
| EWO0260 | *rrsE* 2.2R | GCCCGAAGCTTCCTCCTacagccggttcggttgaagag | This work |
| EWO0261 | *adiY* 1.1F | GGCTGCGAATTCgctaaagcaaagcgataccg | This work |
| EWO0262 | *adiY* 1.1R | GCCCGAAGCTTCCTCCTtttttttgcctgttatttatc | This work |
| EWO0263 | *adiY* 1.2F | GGCTGCGAATTCtttttttgcctgttatttatc | This work |
| EWO0264 | *adiY* 1.2R | GCCCGAAGCTTCCTCCTgctaaagcaaagcgataccg | This work |
| EWO0265 | *valS* 1.1F | GGCTGCGAATTCcgtattcaggttgaaaccag | This work |
| EWO0266 | *valS* 1.1R | GCCCGAAGCTTCCTCCTgatatattgattagtctgcg | This work |
| EWO0267 | *valS* 1.2F | GGCTGCGAATTCgatatattgattagtctgcg | This work |
| EWO0268 | *valS* 1.2R | GCCCGAAGCTTCCTCCTcgtattcaggttgaaaccag | This work |
| EWO0269 | *argeE* 2.1F | GGCTGCGAATTCgttgaatacgctgattgtgg | This work |
| EWO0270 | *argeE* 2.1R | GCCCGAAGCTTCCTCCTtgcggatgcaaatcggagattaac | This work |
| EWO0271 | *argeE* 2.2F | GGCTGCGAATTCtgcggatgcaaatcggagattaac | This work |
| EWO0272 | *argeE* 2.2R | GCCCGAAGCTTCCTCCTgttgaatacgctgattgtgg | This work |
| EWO0273 | *argU* 1.1F | GGCTGCGAATTCggtcgttcacttgttcagcaac | This work |
| EWO0274 | *argU* 1.1R | GCCCGAAGCTTCCTCCTacctgcggcccacgacttag | This work |
| EWO0275 | *argU* 1.2F | GGCTGCGAATTCacctgcggcccacgacttag | This work |
| EWO0276 | *argU* 1.2R | GCCCGAAGCTTCCTCCTggtcgttcacttgttcagcaac | This work |
| EWO0277 | *rpsU* 2.1F | GGCTGCGAATTCctcctcacccttataaaagtc | This work |
| EWO0278 | *rpsU* 2.1R | GCCCGAAGCTTCCTCCTcggcatgtgcctctcacctttg | This work |
| EWO0279 | *rpsU* 2.2F | GGCTGCGAATTCcggcatgtgcctctcacctttg | This work |
| EWO0280 | *rpsU* 2.2R | GCCCGAAGCTTCCTCCTctcctcacccttataaaagtc | This work |
| EWO0281 | *bolA* 2.1F | GGCTGCGAATTCacgaaataatcgcccctggtaaaag | This work |
| EWO0282 | *bolA* 2.1R | GCCCGAAGCTTCCTCCTctagccgcttttaccgtttc | This work |
| EWO0283 | *bolA* 2.2F | GGCTGCGAATTCctagccgcttttaccgtttc | This work |
| EWO0284 | *bolA* 2.2R | GCCCGAAGCTTCCTCCTacgaaataatcgcccctggtaaaag | This work |
| EWO0285 | *guaB* 1.1F | GGCTGCGAATTCgtcgtcaaacgtcagagcttc | This work |
| EWO0286 | *guaB* 1.1R | GCCCGAAGCTTCCTCCTcgcccttcggggatagcaag | This work |
| EWO0287 | *guaB* 1.2F | GGCTGCGAATTCcgcccttcggggatagcaag | This work |
| EWO0288 | *guaB* 1.2R | GCCCGAAGCTTCCTCCTgtcgtcaaacgtcagagcttc | This work |
| EWO0289 | *aroK* 1.1F | GGCTGCGAATTCaagacagcaaaatgccgcctgaatg | This work |
| EWO0290 | *aroK* 1.1R | GCCCGAAGCTTCCTCCTaagcggtaatgtttttacgctgaacg | This work |
| EWO0291 | *aroK* 1.2F | GGCTGCGAATTCaagcggtaatgtttttacgctgaacg | This work |
| EWO0292 | *aroK* 1.2R | GCCCGAAGCTTCCTCCTaagacagcaaaatgccgcctgaatg | This work |
| EWO0293 | *ssuE* 1.1F | GGCTGCGAATTCtacccgccagggtgatgacacgcatac | This work |
| EWO0294 | *ssuE* 1.1R | GCCCGAAGCTTCCTCCTctttagtttattttcagaaaaagatacac | This work |
| EWO0295 | *ssuE* 1.2F | GGCTGCGAATTCctttagtttattttcagaaaaagatacac | This work |
| EWO0296 | *ssuE* 1.2R | GCCCGAAGCTTCCTCCTtacccgccagggtgatgacacgcatac | This work |
| EWO0297 | *tsK* 2.1F | GGCTGCGAATTCgtatgccactgtttgaaaatccc | This work |
| EWO0298 | *tsK* 2.1R | GCCCGAAGCTTCCTCCTgcaatcgattacgtaaatgatagaac | This work |
| EWO0299 | *tsK* 2.2F | GGCTGCGAATTCgcaatcgattacgtaaatgatagaac | This work |
| EWO0300 | *tsK* 2.2R | GCCCGAAGCTTCctcctgtatgccactgtttgaaaatccc | This work |
| EWO0301 | *gdhA* 1.1F | GGCTGCGAATTCctcgttattaatttgctttcctggg | This work |
| EWO0302 | *gdhA* 1.1R | GCCCGAAGCTTCCTCCTtgtctgatccatagatataaaaccc | This work |
| EWO0303 | *gdhA* 1.2F | GGCTGCGAATTCtgtctgatccatagatataaaaccc | This work |
| EWO0304 | *gdhA* 1.2R | GCCCGAAGCTTCCTCCTctcgttattaatttgctttcctggg | This work |
| EWO0305 | *fepB* 1.1F | GGCTGCGAATTCagccctcaccctggaagggag | This work |
| EWO0306 | *fepB* 1.1R | GCCCGAAGCTTCCTCCTatgtcaactcttgaggtaacgc | This work |
| EWO0307 | *fepB* 1.2F | GGCTGCGAATTCatgtcaactcttgaggtaacgc | This work |
| EWO0308 | *fepB* 1.2R | GCCCGAAGCTTCCTCCTagccctcaccctggaagggag | This work |
| EWO0309 | *put* 1.1F | GGCTGCGAATTCgcaaaaaatgtgagagagtgcaacc | This work |
| EWO0310 | *put* 1.1R | GCCCGAAGCTTCCTCCTacggggtaacagagtttatgttttacc | This work |

| | | |
|---|---|---|
| EWO0311 *put* 1.2F | GGCTGCGAATTCacggggtaacagagtttatgttttacc | This work |
| EWO0312 *put* 1.2R | GCCCGAAGCTTCCTCCTgcaaaaaatgtgagagagtgcaacc | This work |
| EWO0313 *mtlA* 1.1F | GGCTGCGAATTCctatatttatgtgattgatatcacac | This work |
| EWO0314 *mtlA* 1.1R | GCCCGAAGCTTCCTCCTgtttgctgtcgcgcagg | This work |
| EWO0315 *mtlA* 1.2F | GGCTGCGAATTCgtttgctgtcgcgcagg | This work |
| EWO0316 *mtlA* 1.2R | GCCCGAAGCTTCCTCCTctatatttatgtgattgatatcacac | This work |
| EWO0317 *purR* 1.1F | GGCTGCGAATTCaatctcccgtcatttataatgataag | This work |
| EWO0318 *purR* 1.1R | GCCCGAAGCTTCCTCCTaaaagtgttgcggtacgccgg | This work |
| EWO0319 *purR* 1.2F | GGCTGCGAATTCaaaagtgttgcggtacgccgg | This work |
| EWO0320 *purR* 1.2R | GCCCGAAGCTTCCTCCTaatctcccgtcatttataatgataag | This work |
| D49724 | ggttggacgcccggcatagtttttcagcaggtcgttg | (5) |

[1]for oligonucleotides, sequences in upper case do not anneal to the DNA template and contain restriction sites for *Eco*RI or *Hin*dIII. Underlined sequences introduce mutations and the symbol Δ indicates a short deletion.
